# Semantic Reversal Anomalies under the Microscope: Task and Modality Influences on Language-Associated Event-Related Potentials

**DOI:** 10.1101/788976

**Authors:** Louise Kyriaki, Matthias Schlesewsky, Ina Bornkessel-Schlesewsky

**Affiliations:** University of South Australia, Cognitive and Systems Neuroscience Research Hub, University of South Australia, GPO Box 2471, Adelaide SA 5001, Australia

## Abstract

Semantic reversal anomalies (SRAs) – sentences where an implausibility is created by reversing participant roles – have attracted much attention in the literature on the electrophysiology of language. In spite of being syntactically well-formed but semantically implausible, these sentences unexpectedly elicited a monophasic P600 effect in English and Dutch rather than an N400 effect. Subsequent research revealed variability in the presence/absence of an N400 effect to SRAs depending on the language examined and the choice of verb type in English. However, most previous studies employed the same presentation modality (visual) and task (acceptability judgement). Here, we conducted two experiments and three statistical analyses to investigate the influence of stimulus modality, task demand, and statistical choices on event-related potential (ERP) response patterns to SRAs in English. We reproduced a previous study’s procedure and analysis (Bourguignon et al., 2012), and further introduced between-subjects factors of task type and modality, using mixed effects modelling to analyse the data. We observed an N400 effect to typical English SRAs (agent subject verbs, e.g. *the fries will eat the boys*), which contrasts existing literature and was not predicted by existing theories that account for SRA processing. Task demand modulated the ERPs elicited by SRAs, while auditory presentation led to increased comprehension accuracy and a more broadly distributed ERP. Finally, the statistical methods used influenced the presence/absence of ERP effects. Our results suggest a sensitivity of language-related ERP patterns to methodological parameters and we conclude that future experiments should take this into careful consideration.

## 1. Introduction

Semantic reversal anomalies (SRAs) and the ERPs they elicit have played an important role in building psycholinguistic theory over the past fifteen years. SRAs arise when the role assignments of two nouns are reversed in a syntactically well-formed sentence, leading to an implausible interpretation (Kim & Osterhout, 2005). For example, the sentence *“The pill will swallow the child with hot tea”* becomes implausible at the verb, *“swallow”* (Bourguignon, Drury, Valois, & Steinhauer, 2012). While *“pill”* is an acceptable direct object (i.e. *“The child will swallow the pill…”*), its inanimacy makes it unsuitable for the subject role, causing a linguistic violation (Bourguignon et al., 2012).

Early SRA experiments observed a P600 rather than the expected N400 to such semantically anomalous verbs. These findings challenged the beliefs of a one-to-one mapping of ERPs onto linguistic processes, where the N400 was thought to reflect semantic processing and the P600 was thought to reflect syntactic processing (e.g. Hagoort, Brown, & Groothusen, 1993; Kutas & Hillyard, 1980; Osterhout & Holcomb, 1992, 1993). In response, two distinct interpretations arose: (1) that very basic assumptions about the timecourse of sentence processing must be rethought, or (2), that the underlying functions which the N400 are thought to reflect must be re-evaluated.

Supporting the first interpretation, Kim and Osterhout (2005) suggested that in SRAs, “semantic attraction” dominates combinatory analysis and overwhelms unambiguous syntactic cues, leading to a semantic analysis. This “semantic attraction” hypothesis conflicts with both modular and interactive views of sentence comprehension (Fodor, 1983; Marslen-Wilson, 1975), and requires a reanalysis of the timecourse of sentence processing. Other studies challenge this interpretation, showing that a P600 can also be elicited by SRA sentences without semantic attraction (Kolk et al., 2003; Van Herten et al., 2005). An alternate interpretation is that the N400 indexes lexical preactivation (Stroud & Phillips, 2012) or the retrieval of lexical information from memory (Brouwer, Fitz, & Hoeks, 2012; Lau, Phillips, & Poeppel, 2008) rather than a more complex aspect of sentence processing. This interpretation suggests that the absence of an N400 effect is due to word and context priming, where the facilitation of verb retrieval is approximately equal for semantically plausible and anomalous phrases (Brouwer et al., 2012). This has implications for how the N400 can be used to conceptualise sentence processing, and is not supported by recent studies which observed an N400 response to SRAs (Bornkessel-Schlesewsky et al., 2011; Bourguignon et al., 2012).

The above interpretations share one similarity in that they depend on the findings of a monophasic P600 elicited by SRAs. However, more recent results suggest a variability of the ERP response to SRAs both across and within languages. Across languages, the presence versus absence of the N400 has been attributed to the relative reliance of a language on particular sentence features for interpretation (Bornkessel-Schlesewsky & Schlesewsky, 2019). In languages which do not rely on word order for interpretation and have a higher reliance on animacy or morphological case marking (“sequence-independent” languages, e.g. German, Turkish, Mandarin Chinese), an N400 is elicited by SRAs (Bornkessel-Schlesewsky et al., 2011)^1^. English and Dutch, which have been the predominant focus of existing SRA investigations, rely minimally on animacy and more heavily on word order as the main cue to sentence interpretation (i.e. they are “sequence-dependent” languages; see MacWhinney, Bates, & Kliegl, 1984). Therefore the lack of N400 effects observed in initial English and Dutch SRA experiments (Hoeks, Stowe, & Doedens, 2004; Kim & Osterhout, 2005; Kolk, Chwilla, Van Herten, & Oor, 2003; Kuperberg, Sitnikova, Caplan, & Holcomb, 2003; van Herten, Chwilla, & Kolk, 2006; Van Herten, Kolk, & Chwilla, 2005) may actually reflect cross-linguistic differences in the cues used for sentence interpretation. Bornkessel-Schlesewsky and Schlesewsky (2019) propose that this can be accounted for by assuming that the N400 reflects precision-weighted prediction error responses. In the predictive coding literature, the term precision is used to index the (un)certainty of a prediction and/or the (un)certainty surrounding the sensory input (Feldman & Friston, 2010) and precision-weighting plays a crucial role in determining the balance between model predictions and sensory input (cf. Adams, Stephan, Brown, Frith, & Friston, 2013). According to Bornkessel-Schlesewsky and Schlesewsky’s account, precision in language depends on the relevance of particular linguistic features for interpretation (cf. the Competition Models’ notion of cue validity; e.g. MacWhinney et al., 1984). Thus, in a language such as German or Mandarin Chinese, in which non-word-order cues such as animacy have a higher precision (cue validity) than in English or Dutch, an animacy-induced SRA elicits an N400 effect, while this is not the case in a sequence-dependent language due to the lower precision of the animacy cue.

The relevance of a particular cue for sentence interpretation can also change depending on the construction of a sentence, resulting in a differential ERP response even within the same language. To date, this type of manipulation has been accomplished using verb type (Bornkessel-Schlesewsky et al., 2011; Bourguignon et al., 2012). An Icelandic experiment found that the N400 and P600 are differentially elicited depending on the combination of verb class and case marking (Bornkessel-Schlesewsky et al., 2011). When “alternating” verbs are used, role assignment is determined by word order and which noun is placed in the subject position, similarly to sentence processing in English or Dutch. When “non-alternating” verbs are used, role assignment is determined by a combination of verb class and case, similarly to German, Turkish, and Mandarin Chinese. Bornkessel-Schlesewsky et al. (2011) found that SRAs involving alternating verbs elicited a P600, patterning with English and Dutch. SRAs involving non-alternating verbs elicited an N400-late positivity pattern, as in German, Turkish and Mandarin Chinese (Bornkessel-Schlesewsky et al., 2011).

In English, Bourguignon et al. (2012) reported novel findings where SRAs involving experiencer subject verbs (ESVs; i.e. *The gifts have loved the children…)* elicited a biphasic N400-late positivity in contrast to typical SRA constructions (using agent subject verbs; ASVs) which elicited a P600, as expected. They suggested that thematic aspect plays a role in SRA processing, and ESVs are less informative to processing because they can be mapped to both subject and object positions (Bourguignon et al., 2012). They interpreted the N400 observed in the ESV condition as reflecting lexical identification of the verb (Bourguignon et al., 2012). If found to be a stable and replicable effect, the English verb-dependent N400 response to SRAs could have significant implications for current psycholinguistic theories. To investigate the verb-class effects on the presence or absence of the N400 and P600, we aim to reproduce the findings of Bourguignon et al. (2012).

Verb-class dependent findings in Icelandic and English provide evidence against the suggestions of Brouwer et al. (2012), Kim & Osterhout (2005) and Stroud & Phillips (2012). When subject and object can be mapped to more than one plausible thematic role (Icelandic alternating verbs, English ESVs) the ERP response is modulated. It is evident that the N400 and P600 do not map one-to-one onto basic lexical processes, but index more complex processing during sentence comprehension. Further, the ERPs elicited by SRAs differ based on the relative prominence of cues to sentence processing, whether that be across languages (e.g. English versus German and the relative prominence of animacy) or within languages (e.g. across verb types in English). These differences highlight the importance of investigating SRAs and the ERPs they elicit in a range of circumstances. Much of the current research has employed a similar experimental environment regarding the choice of languages investigated, tasks used, and stimulus presentation methods. SRAs should be examined in a range of experimental conditions in order to illuminate the generalisability of existing results to aid in the development of existing sentence processing theories (Bornkessel-Schlesewsky, Schlesewsky, Small, & Rauschecker, 2015; Friederici, 2011, 2012; Kim & Osterhout, 2005; Kuperberg, et al., 2003; van de Meerendonk, Kolk, Chwilla & Vissers, 2009).

### 1.1. Task effects

A monophasic P600 has consistently been reported at the verb position of SRAs in sequence-dependent languages (English and Dutch; see Hoeks et al., 2004; Kim & Osterhout, 2005; Kolk et al., 2003; Kuperberg et al., 2003; van Herten et al., 2006; Van Herten et al., 2005), with all but one existing study using visual stimulus presentation combined with an acceptability judgement task^2^. Given the sensitivity of late positivities to task effects and experimental environment, the observed P600 is likely modulated by the design of these experiments (Sassenhagen, Schlesewsky, & Bornkessel-Schlesewsky, 2014). Some suggest the P600 to linguistically deviant stimuli may be a domain-general, late P3 component in response to target events (Coulson, King, & Kutas, 1998; Kutas, Neville, & Holcomb, 1987; Kutas, Van Petten, & Kluender, 2006; Sassenhagen et al., 2014; Van Herten et al., 2005).

An active judgement task increases the saliency of SRA violation verbs, as that is the point where the sentence becomes “unacceptable” and a judgement can be made (Schacht, Sommer, Shmuilovich, Martíenz, & Martín-Loeches, 2014). It has been shown that the P600 response to linguistic violations is larger when combined with an active judgement task compared to passive comprehension (Osterhout, Allen, Mclaughlin, & Inoue, 2002; Osterhout & McKinnon, 1996), a lexical decision (Roehm, Bornkessel-Schlesewsky, Rösler, & Schlesewsky, 2007) identifying the letter case (Gunter & Friederici, 1999), or a probe verification task (Schacht et al., 2014). Thus, in typical delayed-response sentence-processing experiments, a positivity is expected following a task-relevant violation ((Sassenhagen et al., 2014; see Kotz, Frisch, Von Cramon, & Friederici, 2003 for an alternate view).

It is likely that the P600 in SRA experiments is influenced by task demands and stimulus salience, reflecting domain-general rather than higher-order language processes. Bourguignon et al. (2012) also recommended that their verb-type manipulation be retested without an overt judgement task, to determine if the ERP differences between ASV and ESV persist. To address the possibility of P600 effects being task-related, we included a delayed comprehension question task in the present study. When questions cannot be prepared for during comprehension, a task-dependent P600 should be diminished in this condition.

### 1.2. Modality effects

The modality of linguistic stimuli has also been found to influence ERP amplitude, latency, and topography (Domalski, Smith, & Halgren, 1991; Hagiwara, Soshi, Ishihara, & Imanaka, 2007; Holcomb, Coffey, & Neville, 1992; Holcomb & Neville, 1990; Wolff, Schlesewsky, Hirotani, & Bornkessel-Schlesewsky, 2008). In a semantic priming and lexical decision task, Holcomb & Neville (1990) found significant cross-modal differences in the topography of N400s to unrelated words, where visual ERPs were right-lateralised and auditory ERPs were spread across hemispheres. However, others have found no differences between auditory and visual N400s elicited by semantic violations (although the low number of electrodes limits topographical conclusions; Balconi & Pozzoli, 2004). A third modality to consider is sign language, where ERPs elicited by American Sign Language semantic anomalies show differences in latency and morphology compared to both auditory and visual English stimuli (Neville et al., 1997; Neville, Mills, & Lawson, 1992). In the same way that co-articulation effects in the speech stream can provide predictive information about upcoming words, the transition phrase prior to a critical sign provides predictive information in sign language (Hosemann, Herrmann, Steinbach, Bornkessel-Schlesewsky, & Schlesewsky, 2013). This suggest that auditory and sign language may pattern together, as the continuous input stream provides information differently to visual stimulus presentation. When reading, word information is available instantly, and the commonly used method in ERP studies is rapid serial visual presentation (RSVP). RSVP, where single words are presented sequentially on a screen, inherently differs from natural auditory language. When presenting stimuli visually, the RSVP parameters (i.e. the duration that the word is displayed for and/or the duration between each word) may influence the timing of resultant ERPs (Hagoort & Brown, 2000).

Current neurobiological models of sentence processing are based upon the auditory system, and do not make predictions about reading processes (Bornkessel-Schlesewsky & Schlesewsky, 2013; Bornkessel-Schlesewsky, Schlesewsky, Small, & Rauschecker, 2015; Friederici, 2011, 2012). The auditory system may be more sensitive in detecting irregularities or violations in the speech stream compared to reading, as auditory language processing abilities are developed earlier than visual language processing (Holcomb & Neville, 1990). Supporting this, research into semantic borderline anomalies (violations on the borderline of detection) found that violations were detected at a higher rate when stimuli were presented auditorily compared to visually (Bohan, 2007). It is unclear whether we can confidently generalise ERP findings from the visual modality (specifically RSVP) to broader models of natural sentence processing. To investigate the influence of stimulus modality on ERP effects, we developed both a visual and auditory version of the present study for comparison.

### 1.3. The present study

The present study comprises two experiments partially reproducing that of Bourguignon et al. (2012), who used a 2 x 2 design with factors Verb Type (ASV versus ESV) and Animacy (of the Subject Noun). In Experiment 1, we added a between-subjects factor of Task (Judgement or Comprehension) thus employing a 2 x 2 x 2 design (Animacy x Verb Type x Task). Experiment 2 matched Experiment 1, using auditory stimulus presentation rather than visual. Example stimuli are outlined in Table 1. The aims of the present study are: 1) attempt to replicate Bourguignon et al.’s 2012 novel finding of differential ERP patterns to SRAs across verb classes in English, and 2) systematically examine the effects of task and modality on the ERPs elicited by SRAs. Bourguignon et al. (2012) was selected for reproduction in the current study due to their novel study design which manipulated verb type, and the novel findings of an N400 in English SRAs. We predict (H1) that our study will reproduce the behavioural and ERP results of Bourguignon et al. (2012). The main findings of Bourguignon et al. (2012) were: (a) Participants effectively discriminated grammatical from ungrammatical (violation) sentences and were more accurate for ASV than ESV. (b) At the first noun, a significant N400 was elicited for inanimate compared to animate Subject Nouns, most prominent at the midline. (c) At the verb, animacy reversals elicited an N400 effect for ESV only, and elicited a P600 effect for both ASV and ESV.

**Table 1.** Sentence conditions included in the present study. Target words are indicated in **bold** and violations are marked by an asterisk (*). Adapted from Bourguignon et al. (2012).

| Critical Conditions |  |
| --- | --- |
| <u>Agent Subject Verbs (ASV)</u> | <u>Experiencer Subject Verbs (ESV)</u> |
| *Animate – grammatical* | *Animate – grammatical* |
| The boys will **eat** the fries too quickly. | The children will **love** the gifts of the orphanage. |
| The student will **write** the answer on the form. | The judges will **despise** the movies at the festival. |
| *Inanimate – ungrammatical* | *Inanimate – ungrammatical* |
| The fries will * **eat** the boys too quickly. | The gifts will * **love** the children of the orphanage. |
| The answer will * **write** the student on the form. | The movies will * **despise** the judges at the festival. |
| Filler Conditions |  |
| <u>Experiencer Object Verbs (EOV)</u> | <u>Semantic Anomalies (SEA)</u> |
| *Animate – grammatical* | *Animate – grammatical* |
| My achievements will **gladden** her parents a great deal. | The peasants will **peel** the carrots in the kitchen. |
| The toys will **amuse** the kids for many weeks. | The senate will **abolish** the laws on taxes. |
| *Inanimate – ungrammatical* | *Inanimate – ungrammatical* |
| Her parents will * **gladden** my achievements a great deal. | The carrots will * **peel** the peasants in the kitchen. |
| The kids will * **amuse** the toys for many weeks. | The laws will * **abolish** the senate on taxes. |
| <u>Phrase Structure Violations (PSV)</u> |  |
| *Noun phrase first – grammatical* | *Verb phrase first – grammatical* |
| The man chose **the rabbit** to adopt for his kids. | The man chose **to adopt** the rabbit for his kids. |
| The girl helped **the bird** to collect a pile of sticks. | The girl wanted **to collect** a bird from the woods. |
| *Noun phrase first – ungrammatical* | *Verb phrase first – ungrammatical* |
| The man chose **to rabbit** the adopt for his kids. | The man chose **the adopt** to rabbit for his kids. |
| The girl wanted **to bird** a collect from the woods. | The girl helped **the collect** to bird a pile of sticks. |

#### 1.3.1. Experiment 1: Task

We included the between-subjects factor of task to compare the ERPs elicited in a judgement versus comprehension task. The crucial difference between tasks is that the comprehension question could not be predicted prior to presentation. The judgement task employed the same question after each sentence, therefore a decision could be made during sentence presentation. If the P600 observed in SRA studies relates to task-relevance, it should be diminished when the task is unpredictable. We hypothesise (H2) that a P600 will be elicited at the Verb with an Animacy effect and that the amplitude of this P600 will be larger for the Judgement compared to Comprehension condition.

#### 1.3.2. Experiment 2: Modality

Experiment 2 matched Experiment 1, but sentence stimuli were presented auditorily via loudspeakers rather than with RSVP. To test the predictions of current models of sentence processing which are based on auditory processing, we developed an auditory stimulus set with the same sentences as Experiment 1. We hypothesise (H3) that the topography of the N400 effect will differ across modality, with the auditory condition showing more broadly distributed ERP topography.

## 2. Experiment 1: Visual modality

### 2.1. Methods

#### 2.1.1. Participants

Forty-eight right-handed (Edinburgh Handedness Inventory), native English-speaking adults (32 female, mean age = 23.2 ± 0.77, age range 18-40) with normal vision and no history of psychiatric, neurological, cognitive, or language disorders participated after giving informed consent. All participants had normal or corrected-to-normal vision. Participants were offered an honorarium for their participation.

#### 2.1.2. Stimuli

Stimuli were adapted from Bourguignon et al. (2012) with minor alterations listed below. See Bourguignon et al. (2012, pg. 181-183) for further information. Table 1 provides example stimuli, and Appendix A lists all sentence material.

##### 2.1.2.1. Tense alteration

The stimuli created by Bourguignon et al. (2012) were in the present perfect tense *(“The hikers have used the compass… “).* They aimed to have natural sounding sentences for both verb types while including a functional category (i.e. the auxiliary *“has/have”)* to minimize carry-over ERP effects from the Subject Noun to the Verb. The aspect of the present perfect tense varied across the critical conditions (ASV and ESV), potentially confounding results (for further discussion see Bourguignon et al., 2012; Brennan & Pylkkänen, 2010). To avoid this while keeping natural sentences with a functional auxiliary, ASV, ESV, and filler Experiencer Object Verb (EOV) stimuli were adapted to the future indefinite tense (i.e. *“The hikers will use the compass”).* This allowed the consistent use of the auxiliary *“will”* for all critical stimuli, rather than using either *“has”* and *“have”.* An additional motivation for the aspectual change was that we perceived the present perfect tense as sounding unnatural for certain sentences. This was particularly salient for ESV sentences, which describe an internal state where it may be uncommon to use present perfect tense (e.g. present perfect “The children have loved the gifts” versus future “The children will love the gifts”).

##### 2.1.2.2. Re-norming

Word frequency of the adapted future indefinite verbs was recalculated using the British National Corpus (*The British National Corpus, version 3,* 2007). ASV and ESV did not differ in orthographic length or frequency. Animate and Inanimate ESV and ASV Subject Nouns also did not differ in orthographic length or frequency (all*p’s* > .1).

##### 2.1.2.3. Lists

The two complementary stimuli lists were replicated, with eight pseudorandomised versions developed for each task conditions (300 sentences per list, 16 randomisations total). Parameters for pseudorandomisation of the Judgement task condition were: minimum distance of two between condition, no more than three consecutive violations, minimum distance of ten between item number, minimum difference of two between item number. Additional parameters for pseudorandomisation of the Comprehension task condition were: no more than three of the same question type consecutively, no more than two of a correct answer type consecutively, and no more than three repetitions of correct answer location (left/right of screen) consecutively.

##### 2.1.2.4. Comprehension task

The Comprehension task comprised three question types: (1) probing for the Subject Noun, (2) probing for the Object Noun, or (3) a filler yes/no probe question. Below the comprehension question, two answers were presented on the screen, either two nouns or “yes” and “no”. See Table 2 for example comprehension questions. Question type was pseudorandomly allocated to sentence condition and balanced within each presentation list. For each sentence condition within a list, 10 of each question type were allocated to correct sentences and 10 of each to violation sentences. The exception was the phrase structure violation (PSV) filler condition, where the syntax violation made it difficult to create Undergoer and Actor questions, and was allocated a higher ratio of yes/no probe questions^3^.

**Table 2.** Example question and response choices for the Judgement and Comprehension task conditions, including sub-conditions of the comprehension task.

| Task condition | Task subtype | Question presented | Answer choice presented |
| --- | --- | --- | --- |
| Judgement | N/A | Good? | N/A |
|  | Subject probe | Who/what will smoke? | Pipe / Man |
| Comprehension | Object probe | Who/what will be composed? | Writers / Poems |
|  | Yes/no probe | Was there a stage? | Yes / No |

Note. Answer choice indicates the possible answers shown on the left and right side of the screen.

#### 2.1.3. Procedure

The procedure matched that of Bourguignon et al. (2012) with minor alterations as below. The eye-blink prompt was represented by the symbol “!!!”. The font of stimuli was altered to the monospaced font Courier New (size 26), to allow for accurate calculation of spacing between the two comprehension answer choices. Participants were seated 70cm from the computer monitor, and were not shown their electroencephalography (EEG) artefacts. Sentence presentation and the judgement task condition matched that of Bourguignon et al. (2012, pg. 183). The comprehension condition presented a question with two potential answers below. Participants were instructed to select the answer they deemed correct by pressing either the left or right mouse button to select the left- or rightside answer. Prior to the main experiment, participants undertook eight unrelated practice trials (two with linguistic violations) to become familiar with the procedure. The entire testing session lasted between 2 and 2.5 hours.

#### 2.1.4. EEG recording and preprocessing

EEG was continuously recorded from 64 cap-mounted Ag/AgCl active electrodes online referenced to the left mastoid (Acticap, Brain Products, Gilching, Germany). Horizontal and vertical eye movements and blinks were monitored with electrode pairs placed above/below the left eye and at the outer canthi of both eyes. Impedance for each electrode was kept below 5 kOhm. All channels were amplified using a Brain Products ActiChamp amplifier at a 500 Hz sampling rate. Offline data pre-processing and averaging was carried out with MNE-Python version 0.18.2 (Gramfort et al., 2014). A copy of the raw data was bandpass filtered from 0.1 to 30 Hz (zero-phase, hamming windowed finite impulse response (FIR filter; 16,501 sample filter length; 0.1-7.5 Hz transition bandwidth). Electrooculography (EOG) artefacts were identified and removed using the ‘mne.preprocessing.find_eog_events’ function. This function automatically identifies peaks in the EOG channels, and removes the surrounding 500ms from the data. Epochs with peak-to-peak amplitude over 75 microvolts or under 5 microvolts were excluded.

Data for Experiment 1 were analysed via two pipelines: (1) reproducing the original analysis conducted by Bourguignon et al. (2012), and (2) using mixed effect models (MEMs) which can specify random effect structures and examine the data trial-by-trial (Bates, Mächler, Bolker, & Walker, 2014). By utilising random effect structures in MEMs, we are able to index individual-level variance (e.g. by participants, by stimulus items) as random deviations from the fixed effects of interest. For example, including a random intercept by-participants allows each participant to have a different baseline EEG response rather than grouping the data for all participants onto one intercept. For further discussion relating to the application of MEMs onto electrophysiological investigations of language processing, see Payne, Lee, & Federmeier (2015) and Alday, Schlesewsky, & Bornkessel-Schlesewsky (2017).

The reproduction analysis examined the Judgement task subset of participants (*n* = 23). A digital phase FIR band-pass filter (0.4 - 30 Hz) was applied and individual average ERPs were computed for each condition at each electrode in epochs from −200ms to 1100 ms relative to target word onset including a 100ms pre-stimulus baseline. A response-contingent analysis was conducted, where only trials followed by a correct response were analysed. Average amplitude was calculated for the 300-500ms (N400), 700-900ms and 900-1100ms (P600) time-windows. The later P600 timewindow was originally included by Bourguignon et al. (2012) in order to check for differences in the auxiliary positioned prior to the verb. For the MEM analysis, data for both task conditions were included, with all trials included irrespective of behavioural response. The FIR band-pass filter was set at 0.1 – 30 Hz, and trial-by-trial mean prestimulus voltage (−200 to 0 ms) was included as a covariate in the statistical analyses rather than applying a baseline correction (see Alday, 2017). with all other settings were unchanged.

#### 2.1.5. Data analysis

R version 3.5.2 (R Core Team, 2018) was used for all statistical analyses with the packages *tidyverse* version 1.2.1 (Wickham, 2017), *car* version 3.0.2 (Fox & Weisberg, 2019), *lme4* version 1.1.21 (Bates, Maechler, Bolker & Walker, 2015), *effects* version 4.1.0 (Fox, 2019), *reshape2* version 1.4.3 (Wickham, 2007) and *ez* version 4.4.0 (Lawrence, 2016). Plots were created in R using packages *ggplot2* version 3.3.0 (Wickham et al., 2020) and *ggpubr* version 0.3.0 (Kassambara, 2020). Raw data and all analysis scripts are available via the Open Science Framework.

##### 2.1.5.1. Behavioural data

For the reproduction analysis, we subjected judgement acceptability ratings to a global ANOVA with factors Animacy of Subject Noun (Animate or Inanimate) and Verb Type (ASV or ESV). Animate Subject Nouns corresponded to grammatical sentences, while inanimate corresponded to ungrammatical sentences in ASV and ESV (this was opposite for control condition EOV). For the MEM analysis, mixed effects regression models were conducted separately for each task group (Judgement or Comprehension). Model structures included Animacy and Verb Type. For the Comprehension group, Question Type (Actor or Undergoer probe) was also included. Intercepts were grouped by Participant ID. Judgement accuracy (Correct or Incorrect) was specified as the dependent variable. Model structures did not include random intercepts by Item, or random slopes as failure to converge indicated over specification (see Bates et al., 2015 for discussion on developing model structures that are not maximal). Categorical variables were encoded with sum encoding (ANOVA-style encoding; see Appendix B for further details).

##### 2.1.5.2. EEG data

For the reproduction, global ANOVAs for the ERP data included factors Animacy and Verb Type. Forty-two electrodes were analysed in each time window separately for lateral and midline electrodes. The midline included Fz, FCz, Cz, CPz, Pz, and POz, reflected by the factor Anterior-Posterior (6 levels). Lateral electrodes included 36 electrodes (18 over each hemisphere) along three columns of six electrodes each: (1) medial (F1/2, FC1/2, C1/C2, CP1/2, P1/2, PO1/2); (2) intermediate (F3/4, FC3/4, C3/4, CP3/4, P3/4, PO3/4); (3) lateral (F5/6, FC5/6, C5/6, CP5/6, P5/6, PO5/6). Thus, topographical factors included in the ANOVAs were Hemisphere (2 levels), Column (3 levels) and Anterior-Posterior (6 levels). Significant interactions (*p* < .05) were followed by step-down analyses. The Greenhouse-Geisser correction is reported where necessary.

For the MEM analysis, MEMs fit by restricted maximum likelihood were undertaken for average amplitude in each ERP time window. Fifty-three electrodes were analysed, with the midline containing 13 (F1/z/2, FC1/2, C1/z/2, CP1/z/2, Pz, POz) electrodes. Lateral electrodes included 30 electrodes (15 over each hemisphere). Electrodes were assigned to the left, midline, or right laterality and anterior, central or posterior location sagitality. For the left laterality, anterior contained electrodes F7, F5, F3, FT7, FC5 and FC3, central contained T7, C5, C3, TP7, CP5 and CP3, while posterior contained PO7, PO3 and O1. For the right laterality, anterior contained electrodes F8, F6, F4, FT8, FC6 and FC4, central contained T8, C6, C4, TP8, CP6 and CP4, while posterior contained PO8, PO4 and O2. Model structures included Animacy (Animate or Inanimate), Verb Type (ASV or ESV), Laterality (Left, Midline, Right), Sagitality (Anterior, Central, Posterior). Scaled Prestimulus Amplitude was included in the model as a main effect. Intercepts were grouped by Participant ID and Item. Model structures did not include random slopes as failure to converge indicated over specification. Categorical variables were encoded with sum encoding (ANOVA-style encoding; see Appendix B for further details and model summaries).

### 2.2. Results

#### 2.2.1. Behavioural data

##### 2.2.1.1. Reproduction

Acceptability rates for grammatical sentences were 89.67% for ASV and 80.98% for ESV, and for ungrammatical sentences were 7.91% for ASV and 15.48% for ESV. A repeated measures ANOVA included a significant main effect of Animacy (F (1,21) = 1194.65; *p* < .001), showing that participants effectively discriminated grammatical and ungrammatical sentences. A significant Animacy x Verb Type interaction (F (1,21) = 62.81; *p* < .001) showed that discrimination was more successful in the ASV condition compared to the ESV condition. Follow up ANOVAs were conducted separately for grammatical and nongrammatical conditions, each showing a significant effect of Verb Type. This showed the higher accuracy of ASV over ESV held for accepting grammatical (F (1,21) = 11.63; *p* = .003) and rejecting ungrammatical sentences (F (1,21) = 20.98; *p* < .001).

##### 2.2.1.2. MEM

Response accuracy was analysed rather than acceptance ratings, and coded by correct acceptance, incorrect acceptance, correct rejection, or incorrect rejection. The task groups were analysed separately, as the Comprehension group included the additional factor of Question Type. For the Judgement condition, a significant interaction effect of Animacy x Verb Type was observed (χ^2^(1) = 4.95, *p* =.026), where accuracy was higher for ASV than ESV. For Comprehension, a significant effect of Animacy x Verb Type x Question Type was observed (χ^2^(1) = 434.63,*p* <.001). When probed for the Actor (e.g. “Who/what will eat?”), participants performed similarly for grammatical and ungrammatical ASV sentences, but were significantly more accurate for grammatical compared to ungrammatical ESV sentences. When probed for the Undergoer (e.g. “Who/what will be eaten?), participants were most accurate for grammatical compared to ungrammatical sentences. See Table 3 for response accuracy percentages.

**Table 3.**
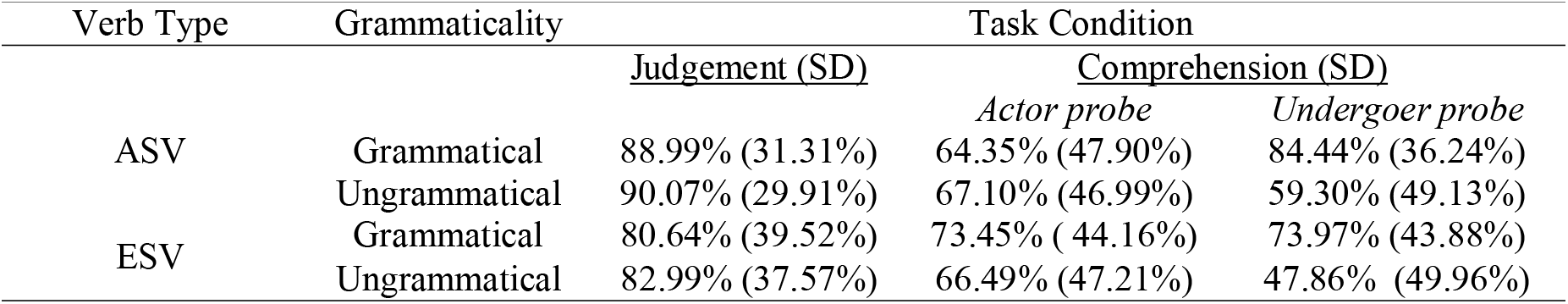
Participant response accuracy to ASV and ESV sentences, by Verb Type, Grammaticality, and Task for Experiment 1 (visual modality).

| Verb Type | Grammaticality | Task Condition |  |  |
| --- | --- | --- | --- | --- |
|  |  | Judgement (SD) | Comprehension (SD) |  |
|  |  |  | Actor probe | Undergoer probe |
| ASV | Grammatical | 88.99% (31.31%) | 64.35% (47.90%) | 84.44% (36.24%) |
|  | Ungrammatical | 90.07% (29.91%) | 67.10% (46.99%) | 59.30% (49.13%) |
| ESV | Grammatical | 80.64% (39.52%) | 73.45% (44.16%) | 73.97% (43.88%) |
|  | Ungrammatical | 82.99% (37.57%) | 66.49% (47.21%) | 47.86% (49.96%) |

#### 2.2.2. ERP data

##### 2.2.2.1. Reproduction

For the Subject Noun, a negative deflection was observed in the N400 time-window (300-500ms) for inanimate compared to animate Subject Nouns in ASV and ESV sentences. A global ANOVA showed a main effect of Animacy on the midline (F(1,21) = 8.84; *p* = .007) with a follow-up ANOVA showing this was most significant at posterior regions (e.g. Pz, F(1,21) = 11.39; *p* = .003). An Animacy x Column x Anterior-Posterior interaction (F(3.84,80.60) = 3.53, *p* = .011) indicated that the N400 was more prominent near the midline (F1/2 columns, F(1,21) = 8.73, *p* = .008; F3/4 columns, F(1,21) = 9.91, *p* = .005) compared to the lateral electrodes (F5/6 columns, F(1,21) = 4.84, *p* = .039). The N400 effect was also largest at central and posterior regions (e.g. CP row, F(1,21) = 11.53; *p* = .003 and P row, F(1,21) = 10.72; *p* = .004). No statistically significant Animacy effects were observed in the 700-900 or 900-1100ms time-windows, as reported in Bourguignon et al. (2012), who posited that this reflects an absence of difference in the auxiliary preceding the verb.

In the verb N400 time-window, an ANOVA including both sentence conditions did not show a significant Animacy x Verb Type interaction for the midline or lateral electrode factors (F(1,21) = 3.78, *p* = .065), with visual inspection of the ERP plots suggesting a small negativity for ungrammatical ESV sentences. In the verb 700-900ms time-range, an Animacy x Verb Type effect was observed, where violations elicited a positivity for ASV but not ESV (F(1,21) = 13.16, *p* = 0.02). An interaction of Animacy x Column x Anterior Posterior (F(4.15, 87.21) = 3.64, *p* = .008) was observed, but follow-up analyses showed no significant effects. At lateral electrodes, an interaction of Animacy x Verb Type x Column x Hemisphere was also observed (F(1.30, 27.34) = 9.13, *p* = .003). Follow-up analyses were conducted for each verb type separately, with no significant effects found for the ESV condition. The ASV condition showed a significant Animacy x Column effect (F(1.10,23.12) = 23.73,*p* < .001) where the positivity was largest at medial compared to lateral electrodes. The ASV condition also showed a significant main effect of Animacy on Hemisphere (F(1,21) = 7.07, *p* = .015), with the positivity larger in the right hemisphere. See Figure 1 for ERP plots at the Subject Noun and the verb analysed using the Bourguignon et al. (2012) pipeline, displayed with and without a −100ms to 0ms pre-stimulus baseline correction.

**Figure 1.**
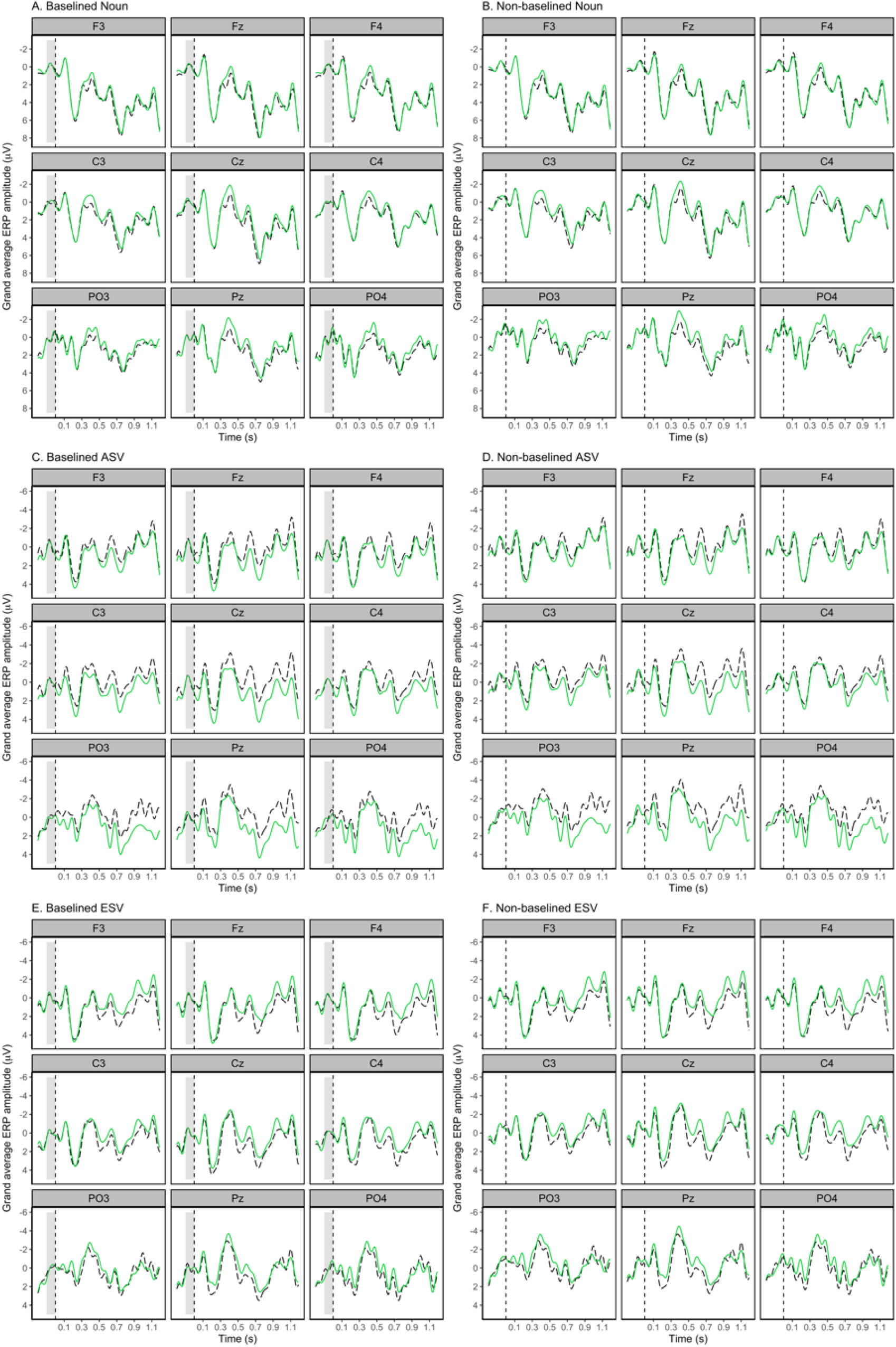
Grand average waveform of the visual ERPs analysed using the procedure reported in Bourguignon et al. (2012), with no baseline correction applied (A-C, baseline shaded grey) and with a baseline correction of −100ms to 0ms pre-stimulus onset as conducted in the original study (D-F). A and D show the ERP time-locked to noun onset at the Subject Noun of Agent- and Experiencer-Subject Verb sentences (ASV and ESV, conditions collapsed), where black/dashed lines represent animate Subject Nouns and green/solid lines represent inanimate Subject Nouns. A significant N400 effect is observed for inanimate compared to animate Subject Nouns. B and E show the ERP time-locked to verb onset for the ASV condition, with C and F time-locked to verb onset for the ESV condition. For the verb plots, black/dashed lines represent correct/grammatical sentences while green/solid lines represent incorrect/ungrammatical sentences. At the verb, no significant effects were found in the N400 time-window for ASV or ESV, and in the P600 time-window, a significant effect was observed for the ASV condition only. Negativity is plotted upwards and waveforms are time-locked to the onset of the noun or verb with a −200ms to 0ms prestimulus interval. Here, the visual difference between the baseline-correct and non-baseline-corrected ERP waveforms appears minimal.

##### 2.2.2.2. MEM

In the Subject Noun N400 time-window, a main effect of Animacy (χ^2^(1) = 149.50, *p* <.001) showed that Inanimate Subject Nouns elicited a negative effect compared to Animate Subject Nouns. An Animacy x Task effect (χ^2^(1) = 121.12, *p* <.001) was observed, which was larger for the Judgement compared to Comprehension condition. An interaction of Animacy x Laterality (χ^2^(2) = 22.42, *p* <.001) shows that the N400 was larger in the right hemisphere. In the P600 time-window, we observed an Animacy x Task effect (χ^2^(1) = 44.90, *p* <.001). This interacted with Laterality (Animacy X Task x Laterality, χ^2^(2) = 8.21, *p* =.016) and Sagitality (Animacy x Task x Sagitality, χ^2^(2) = 7.27, *p* =.026). In the later window (900-1100ms) a main effect of Animacy (χ^2^(2) = 63.95,*p* <.001) was observed. However, the interactions in the 700-900 and 900-1100ms windows did not resolve to simple effects in any topographical region upon visual inspection of the model plots (see Figure 2 for ERP plots and Figure 3 for model plots; see Appendix B for model summaries and Appendix C for ERP plots including 95% confidence intervals by subject and by item for all analyses reported).

**Figure 2.**
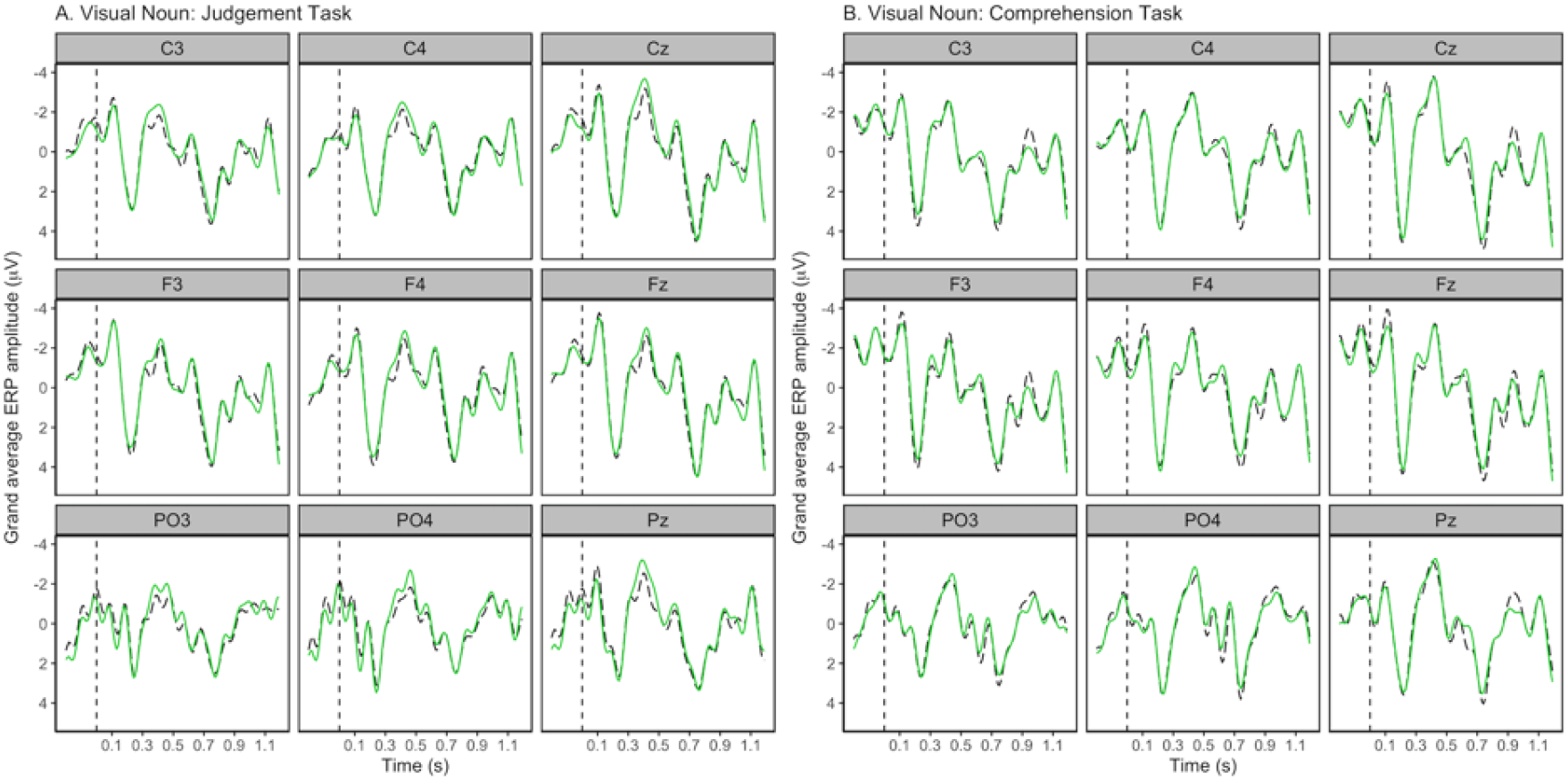
Grand average waveform of the visual ERPs elicited at the Subject Noun of Agent- and Experiencer-Subject Verb sentences (conditions collapsed) for the Judgement and Comprehension task conditions. Black/dashed lines represent animate Subject Nouns while green/solid lines represent inanimate Subject Nouns. Negativity is plotted upwards and waveforms are time-locked to the onset of the noun with a −200ms to 0ms prestimulus interval. Inanimate Subject Nouns elicited an N400 effect relative to animate Subject Nouns, and this effect was larger for the Judgement condition compared to comprehension condition. No significant effects in the 700-900 or 900-1100ms time windows were observed.

**Figure 3.**
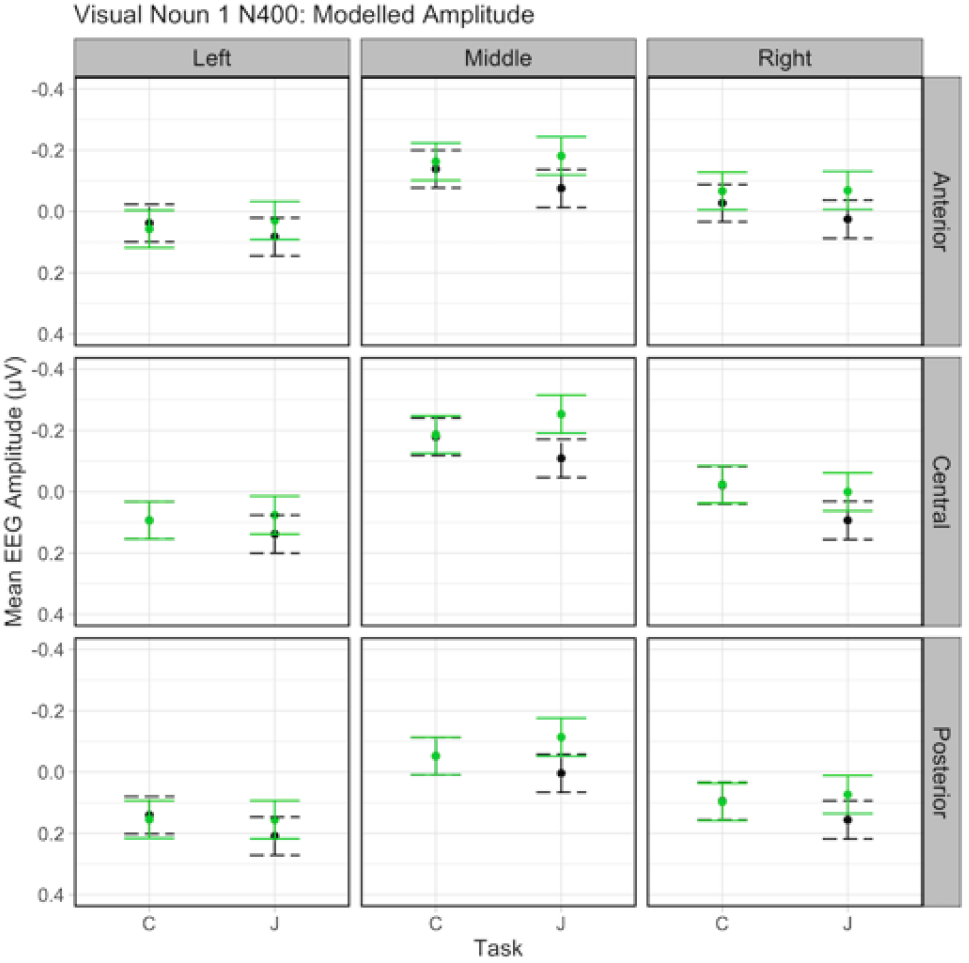
Interaction of Subject Noun animacy, task type and region of interest on mean EEG amplitude for the Subject Noun N400 time window in the visual experiment. Black/dashed lines represent animate Subject Nouns and green/solid lines represent inanimate Subject Nouns. Bars indicate 95% confidence intervals. Effects are shown for nine topographical regions of interest. C = comprehension task, J = judgement task.

For the Verb, in the 300-500ms time-window, an interaction of Animacy x Verb Type x Task (χ^2^(1) =117.46, *p* <.001) indicated that for ASV sentences, violations elicited an N400 for the Judgement condition only. For ESV sentences, an N400 was elicited for the Comprehension but not Judgement condition. This interacted with Laterality (Animacy x Verb Type x Task x Laterality, χ^2^(2) = 24.44, *p* <.001) and Sagitality (Animacy x Verb Type Task x Sagitality, χ^2^(2) = 5.88, *p* =.053) and the N400 effect was largest for electrodes at middle/right Laterality and anterior/central Sagitality. In the 700-900ms time-window, an interaction of Animacy x Verb Type x Task (χ^2^(1) =22.42, *p* <.001) indicated a significant positivity for violation ASV sentences in the Comprehension group. An interaction with Laterality (Animacy x Verb Type x Task x Laterality (¿(12.87) =11.70, *p* =.002), showed left-lateralisation of this effect. Visual inspection of the ERP plots (see Figure 4) suggests a positivity in this time-window for violation ASV sentences in the Judgement group, but this was not statistically significant (although a significant interaction was observed in the reproduction analysis). Model plots are included in Figure 5.

**Figure 4.**
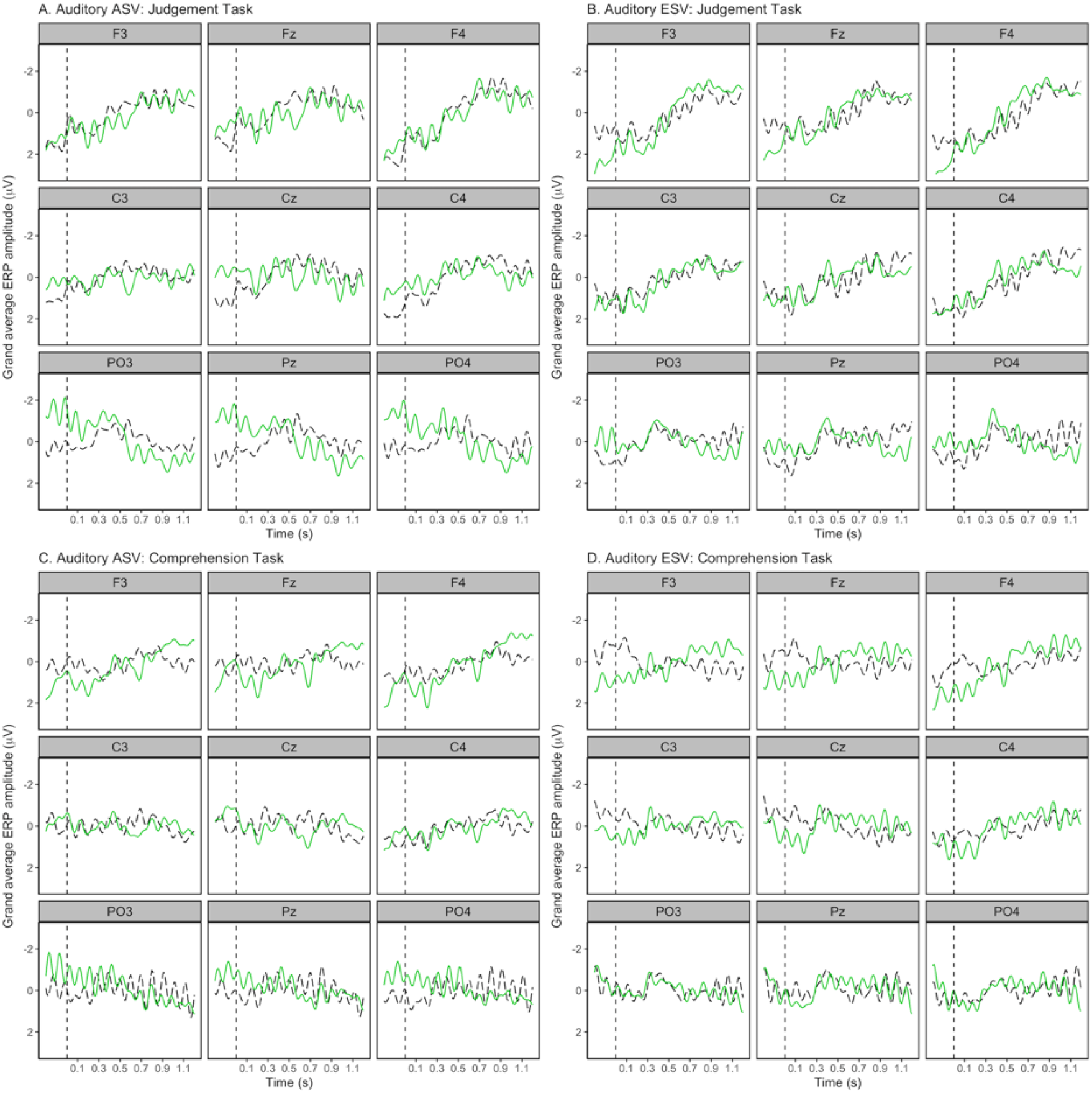
Grand average waveform of the visual ERPs elicited at the verb of Agent Subject Verb (ASV) and Experiencer Subject Verb (ESV) sentences for the Judgement and Comprehension task conditions. Black/dashed lines represent correct/grammatical sentences while green/solid lines represent incorrect/ungrammatical sentences. Negativity is plotted upwards and waveforms are time-locked to the onset of the verb with a −200ms to 0ms prestimulus interval. In the N400 time-window, ASV violations elicited a significant effect for the Judgement condition only, while ESV violations elicited a significant effect for the Comprehension condition only. In the P600 time-window, ASV violations elicited a significant effect for the Comprehension group only, with visual inspection of the plots showing a non-significant positivity for the Judgement group also. No effects were observed in the P600 time-window for ESV violations.

**Figure 5.**
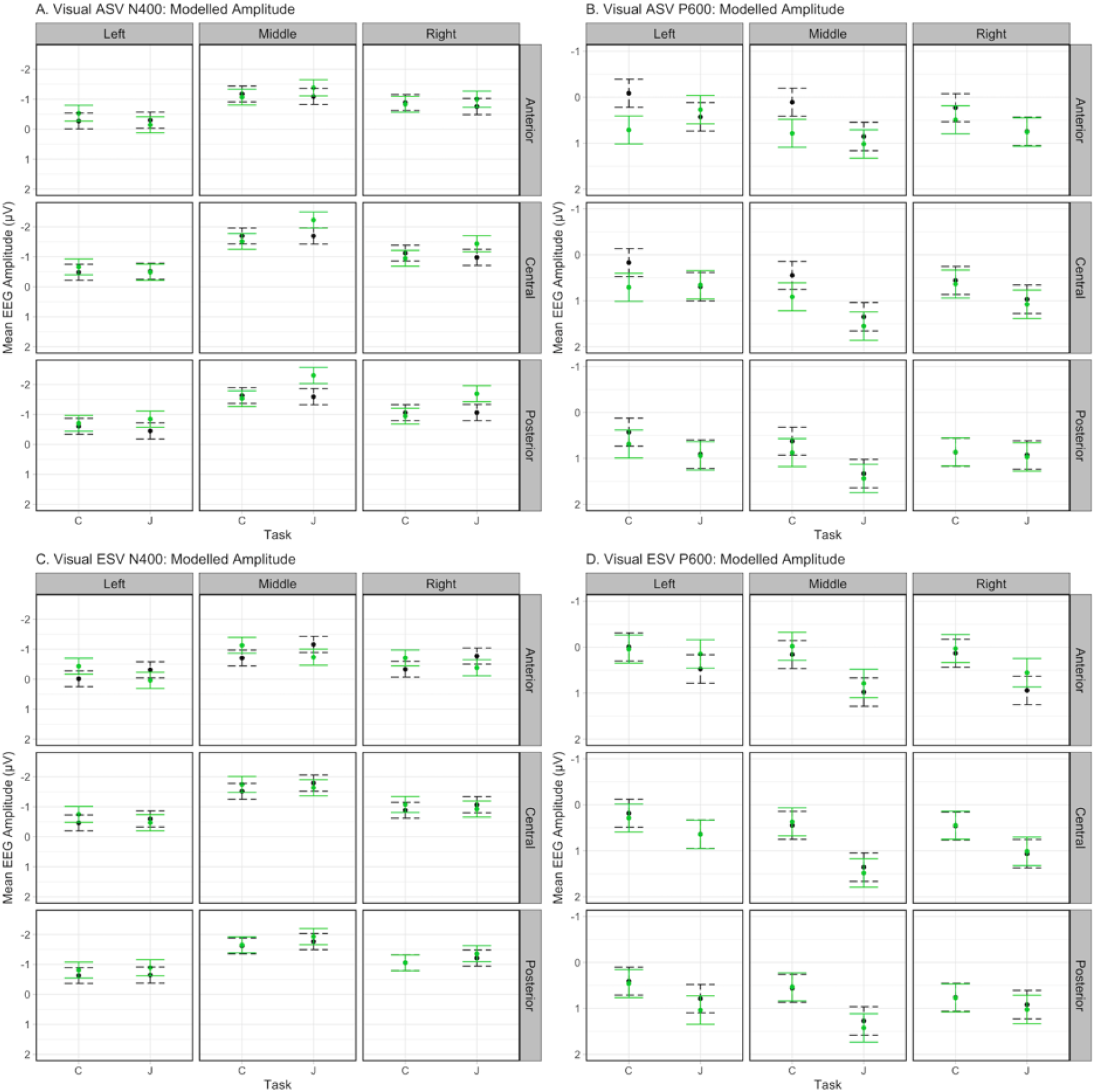
Interaction of sentence grammaticality, task type and region of interest on mean EEG amplitude for the Agent Subject Verb (ASV) and Experiencer Subject Verb (ESV) N400 and P600 time windows in the visual experiment. Black/dashed lines represent correct/grammatical sentences while green/solid lines represent incorrect/ungrammatical sentences. Bars indicate 95% confidence intervals. Effects are shown for nine topographical regions of interest.

## 3. Experiment 1: Visual modality summary

Using ANOVAs, we reproduced Bourguignon et al.’s behavioural results and N400 effect for inanimate Subject Nouns. At the verb, we did not find a significant Animacy x Verb Type interaction in the N400 time-window, where Bourguignon et al. observed an N400 effect to ESV violations. In the verb P600 time-window, we observed a P600 to violations in the ASV condition only, rather than both verb conditions as expected.

The MEM analysis showed the same behavioural results and ERP effects at the Subject Noun. The N400 at the Subject Noun was larger for the Judgement compared to the Comprehension condition. At the verb, ASV violations elicited an N400 for the Judgement condition, while ESV violations elicited an N400 for the Comprehension condition. The lack of N400 effect for ESV violations in the Judgement condition was unexpected. In the Verb P600 time window, a significant positivity was observed for violation ASV sentences in the Comprehension condition only. We did not find a P600 effect for either task group in the ESV condition.

## 4. Experiment 2: Auditory modality

### 4.1. Methods

#### 4.1.1. Participants

Forty-eight right-handed (Edinburgh Handedness Inventory), native English-speaking adults (27 female, mean age = 24.9 ± 0.8, age range 18-39) participated after giving informed consent. Participants fulfilled the same inclusion criteria as Experiment 1, with the added criterion of no self-reported hearing impairments, and had not participated in Experiment 1. They were offered an honorarium for participation.

#### 4.1.2. Stimuli

Stimuli were recorded in a silent room using professional recording equipment (16-bit 44.1 kHz Wav format) by a female native English speaker. Initial recording was split over two consecutive days, and the speaker returned on a third day to re-record a subset of stimuli that possessed some external disturbance, poor voice quality or intonation. All stimuli were recorded in a natural and neutral prosody that did not give cues to violations or the nature of the sentence. Individual sentences were extracted from the larger recording chunks manually. The intensity of all audio files was averaged to a mean value to normalise the intensity of each individual file. The duration and pitch of recordings was not altered to retain the naturalness of stimuli.

##### 4.1.2.1. Acoustic analyses

Speech-analysis program Praat (Boersma, 2006) was used to measure the duration, intensity, and fundamental frequency (F0)/pitch of each word.

###### 4.1.2.1.1. Duration

Duration analyses were performed for the Subject Noun, auxiliary, and verb of critical sentences. Mean and standard deviation values are presented in Table 4. ANOVAs were computed to determine significant differences across conditions. At the Subject Noun, a significant difference *(p* = .002) in duration between ASV and ESV was observed where Subject Nouns were longer in violation ESV sentences than in violation and control ASV sentences. For the auxiliary (always the word “will”), a significant Verb Type x Violation interaction was observed *(p* = .017). There was a significant difference between ASV and ESV *(p* = .02) but not between control and violation sentences either between or within conditions *(p* <.05)^4^. No significant differences were observed in duration between ASV and ESV verbs (all *p*’s < .1).

**Table 4.** Mean and standard deviation word duration (milliseconds) of the Subject NP, auxiliary, and verb of ASV and ESV sentences in Experiment 2.

| Verb Type | Grammaticality | Word Duration |  |  |
| --- | --- | --- | --- | --- |
|  |  | <u>Subject NP (SD)</u> | <u>Auxiliary (SD)</u> | <u>Verb (SD)</u> |
| ASV | Grammatical | 453.33 (124.50) | 153.67 (33.30) | 387.00 (121.74) |
|  | Ungrammatical | 451.33 (110.86) | 176.67 (43.76) | 397.00 (102.72) |
| ESV | Grammatical | 492.67 (134.91) | 190.17 (70.70) | 380.17 (110.45) |
|  | Ungrammatical | 519.17 (156.15) | 185.00 (74.07) | 374.33 (114.29) |

###### 4.1.2.1.2. Intensity (dB)

ANOVAs were computed for the critical conditions to observe the effect of condition and violation on the intensities of the Subject Noun, auxiliary, and verb. Table 5 shows the mean and standard deviation intensities. No significant difference in intensity was found between conditions for the Subject Noun or auxiliary. For the verb, a significant effect of condition *(p* = .017) indicated that ESV had a higher mean sound intensity than ASV, although mean values differ by only 0.5 dB.

**Table 5.** Mean and standard deviation word duration intensity (dB) of the Subject NP, auxiliary, and verb of ASV and ESV sentences in Experiment 2.

| Verb Type | Grammaticality | Word |  |  |
| --- | --- | --- | --- | --- |
|  |  | <u>Subject NP (SD)</u> | <u>Auxiliary (SD)</u> | <u>Verb (SD)</u> |
| ASV | Grammatical | 71.98 (1.29) | 70.97 (1.82) | 70.18 (1.67) |
|  | Ungrammatical | 71.95 (1.28) | 70.58 (1.60) | 70.19 (1.48) |
| ESV | Grammatical | 71.66 (1.23) | 70.95 (1.36) | 70.71 (1.45) |
|  | Ungrammatical | 71.73 (1.09) | 70.67 (1.27) | 70.59 (1.33) |

###### 4.1.2.1.3. Fundamental frequency/pitch (Hz)

As the fundamental frequency (F_0_) differences between words gives rise to the perception of intonation or prosody, the minimum, maximum, and mean F0 values were measured. These values were analysed using ANOVAs for the Subject Noun, auxiliary, and verb. No significant differences in pitch mean were found (all*p*’s <. 1). For pitch minimum, a significant difference was found at the Subject Noun, where control nouns had a higher mean pitch minimum than violation *(p* = .018). For pitch maximum, a significant difference across conditions was found at the auxiliary *(p* = .03) where ASV had a higher mean pitch maximum than ESV. See Table 6 for mean and standard deviations of F_0_.

**Table 6.** Mean, minimum, maximum and standard deviations of Fundamental Frequency (Hz) of the Subject NP, auxiliary, and verb of ASV and ESV sentences in Experiment 2.

| Word | Verb Type | Grammaticality | Measure |  |  |
| --- | --- | --- | --- | --- | --- |
|  |  |  | <u>Mean (SD)</u> | <u>Min (SD)</u> | <u>Max (SD)</u> |
| Subject NP | ASV | Grammatical | 167.47 (6.80) | 148.43 (8.35) | 182.03 (22.11) |
|  |  | Ungrammatical | 167.49 (8.31) | 143.20 (21.34) | 180.88 (10.15) |
|  | ESV | Grammatical | 168.51 (9.58) | 147.49 (9.83) | 181.58 (11.83) |
|  |  | Ungrammatical | 168.71 (7.21) | 145.47 (15.39) | 187.72 (23.37) |
| Auxiliary | ASV | Grammatical | 159.49 (7.19) | 151.56 (8.34) | 168.50 (9.41) |
|  |  | Ungrammatical | 159.47 (16.13) | 146.38 (22.87) | 172.59 (25.52) |
|  | ESV | Grammatical | 157.25 (7.64) | 149.42 (7.93) | 164.57 (10.24) |
|  |  | Ungrammatical | 159.42 (6.69) | 151.26 (10.58) | 168.04 (9.93) |
| Verb | ASV | Grammatical | 155.93 (7.63) | 145.71 (14.94) | 168.38 (8.90) |
|  |  | Ungrammatical | 154.76 (6.97) | 141.39 (18.57) | 169.74 (12.07) |
|  | ESV | Grammatical | 154.26 (6.26) | 138.06 (21.06) | 166.75 (13.00) |
|  |  | Ungrammatical | 155.40 (6.43) | 144.05 (13.41) | 169.16 (24.24) |

### 4.2. Procedure

The procedure was matched to Experiment 1, with the only change being the presentation of sentence stimuli via loudspeakers (mono front speaker format). Each trial comprised a 200ms blink prompt, followed by a fixation cross presented for 500ms pre-stimulus onset. While the audio was presented, the fixation cross remained onscreen. Judgement and comprehension questions were presented visually as in Experiment 1.

### 4.3. Data analysis

Data analysis was identical to that of the MEM analysis for Experiment 1.

### 4.4. Results

#### 4.4.1. Behavioural data

For the Judgement condition, significant main effects of Animacy (χ^2^(1) = 13.37,*p* <.001) and Verb Type (χ^2^(1) = 5528.11, *p* < .001) were observed, with accuracy significantly higher for ASV than ESV, and higher for grammatical than ungrammatical. For the Comprehension condition, significant effects of Animacy x Verb Type (χ^2^(1) = 607.47, *p* <.001), Animacy x Question Type (χ^2^(1) = 1464.64, *p* <.001), and Verb Type x Question Type were observed (χ^2^(1) = 1309.76, *p* <.001). When probed for the Actor, participants were more accurate in for grammatical sentences compared to ungrammatical sentences. When probed for the Undergoer participants were most accurate for grammatical compared to ungrammatical sentences. Accuracy percentages are reported in Table 7.

**Table 7.** Participant response accuracy to ASV and ESV sentences, by Verb Type, Grammaticality, and Task for Experiment 2 (auditory modality).

| Verb Type | Grammaticality | Task Condition |  |  |
| --- | --- | --- | --- | --- |
|  |  | Judgement | Comprehension |  |
|  |  |  | Actor probe | Undergoer probe |
| ASV | Grammatical | 92.40% (26.50%) | 86.32% (34.37%) | 90.71% (29.03%) |
|  | Ungrammatical | 92.71% (26.00%) | 77.21% (42.00%) | 72.11% (44.85%) |
| ESV | Grammatical | 86.72% (33.93%) | 83.33% 37.27% | 81.10% (39.19%) |
|  | Ungrammatical | 86.99% (33.65%) | 78.80% (40.87%) | 63.84% (48.04%) |

#### 4.4.2. ERP data

##### 4.4.2.1. Subject Noun

In the N400 time-window, an interaction effect of Animacy x Task (χ^2^(1) = 30.92, *p* < .001) showed that inanimate Subject Nouns elicited a negative effect compared to animate, larger for the Judgement compared to the Comprehension condition. An Animacy x Sagitality interaction (χ^2^(2) = 114.68, *p* <.001) indicated that this negativity was maximal at posterior and central regions. In the P600 time-window, there was an interaction of Animacy X Task x Laterality (χ^2^(2) = 36.87,*p* <.001). In the later window (9900-1100ms) the same interaction was observed (χ^2^(2) = 10.26, *p* =.006), along with an interaction of Animacy X Task X Sagitality (χ^2^(2) = 23.40, *p* <.001). Interactions in the 700-900 and 900-1100ms windows did not resolve to simple effects upon visual inspection of the ERP (see Figure 6) or model plots (see Figure 7).

**Figure 6.**
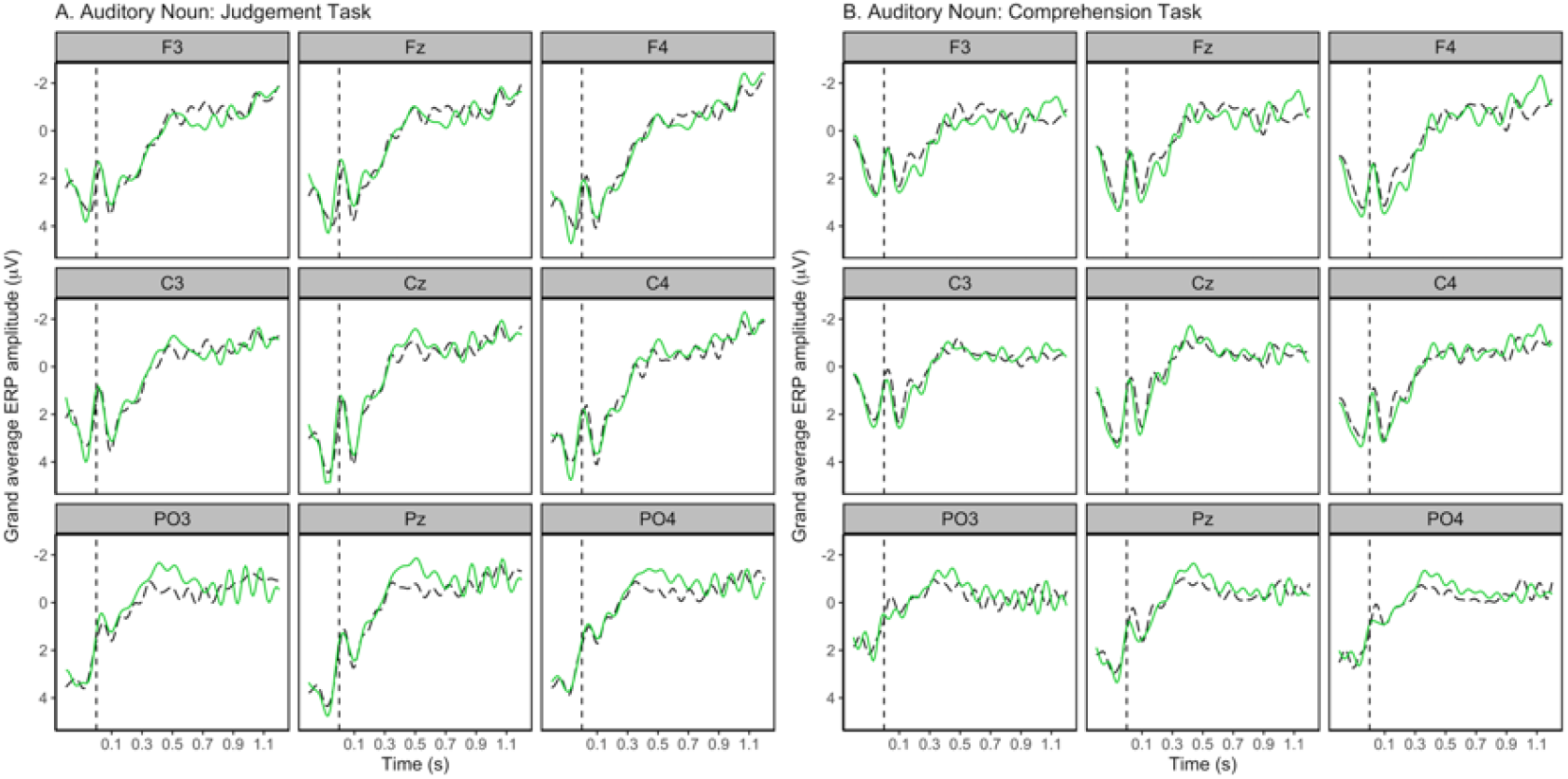
Grand average waveform of the auditory ERPs elicited at the Subject Noun of Agent Subject Verb and Experiencer Subject Verb sentences (conditions collapsed) Judgement and Comprehension task conditions. Black/ dashed lines represent animate Subject Nouns while green/solid lines represent inanimate Subject Nouns. Negativity is plotted upwards and waveforms are time-locked to the onset of the noun with a −200ms to 0ms prestimulus interval. Inanimate Subject Nouns elicited an N400 effect relative to animate Subject Nouns, and this effect was larger for the Judgement condition compared to Comprehension condition. No significant effects in the 700-900 or 900-1100ms time windows were observed.

**Figure 7.**
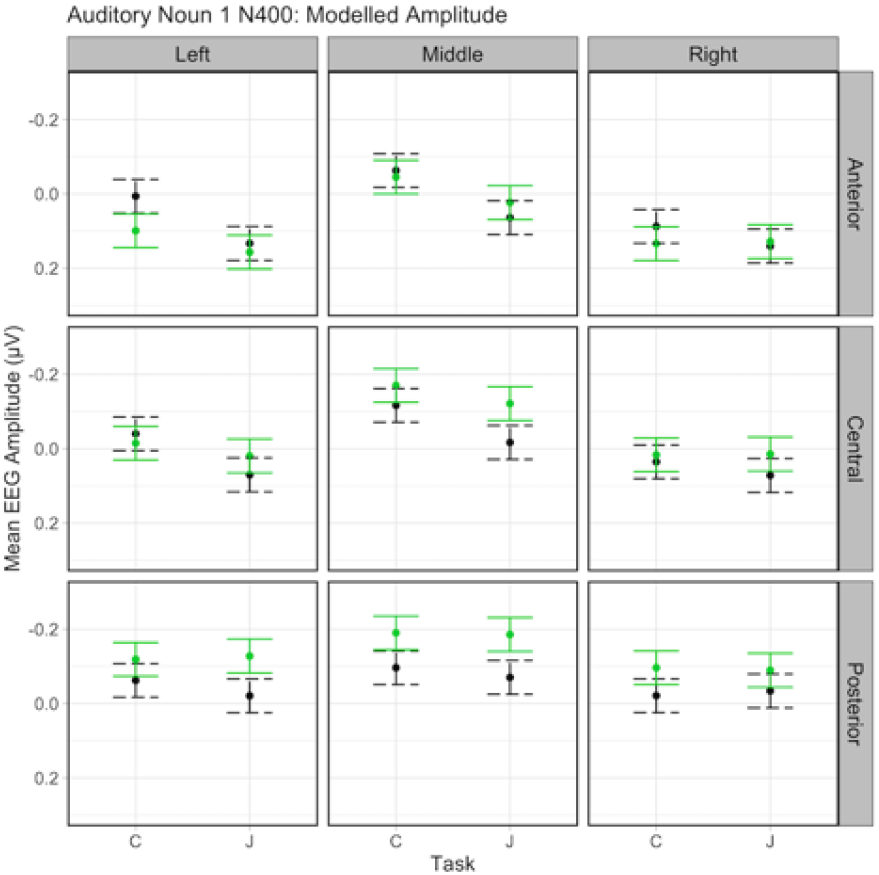
Interaction of Subject Noun animacy, task type and region of interest on mean EEG amplitude for the Subject Noun N400 time window in the auditory experiment. Black/dashed lines represent animate Subject Nouns and green/solid lines represent inanimate Subject Nouns. Bars indicate 95% confidence intervals. Effects are shown for nine topographical regions of interest. C = comprehension task, J = judgement task.

##### 4.4.2.2. Verb

In the 300-500ms time-window, an interaction effect of Animacy x Verb Type x Task (χ^2^(1) =10.61, *p* =.001) was observed. There were also interactions of Animacy x Task x Laterality (χ^2^(2) =27.90, *p* <.001), Verb Type x Task x Laterality (χ^2^(2) =14.36, *p* <.001), Animacy x Task x Sagitality (χ^2^(2) =23.243, *p* <.001), Verb Type x Task x Sagitality (χ^2^(2) =9.25, *p* =.01). ERP and model plots show no N400 effect for the Comprehension condition. For the Judgement condition, violations for ASV and ESV elicited an N400 effect, most prominent in the right hemisphere. Visual inspection of the ERPs suggests this effect was larger for ESV verb than ASV verbs (see Figure 8 for ERP plots and Figure 9 for model plots).

**Figure 8.**
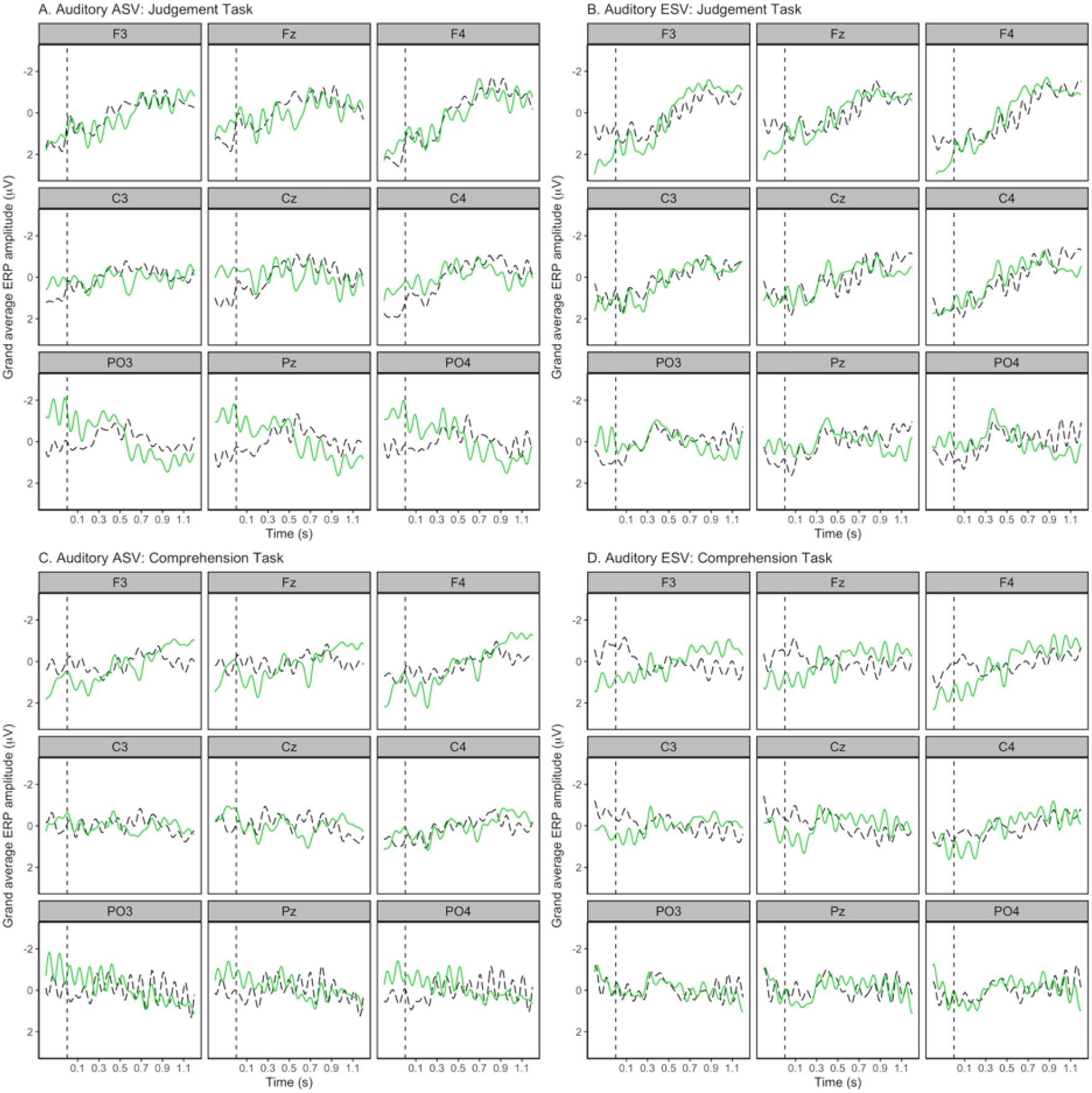
Grand average waveform of the auditory ERPs elicited at the Verb of Agent Subject Verb (ASV) and Experiencer Subject Verb (ESV) sentences for the Judgement and Comprehension task conditions. Black/dashed lines represent correct/grammatical sentences while green/solid lines represent incorrect/ungrammatical sentences. Negativity is plotted upwards and waveforms are time-locked to the onset of the verb with a −200ms to 0ms prestimulus interval. In the N400 time-window, ASV and ESV violations elicited a significant effect for the Judgement condition only. In the P600 time-window, ASV violations elicited a significant effect for the Judgement and Comprehension groups. No significant effects were observed in the P600 time-window for ESV violations, although visual inspection of the plots shows an ongoing negativity toward ESV violations in the Comprehension group.

**Figure 9.**
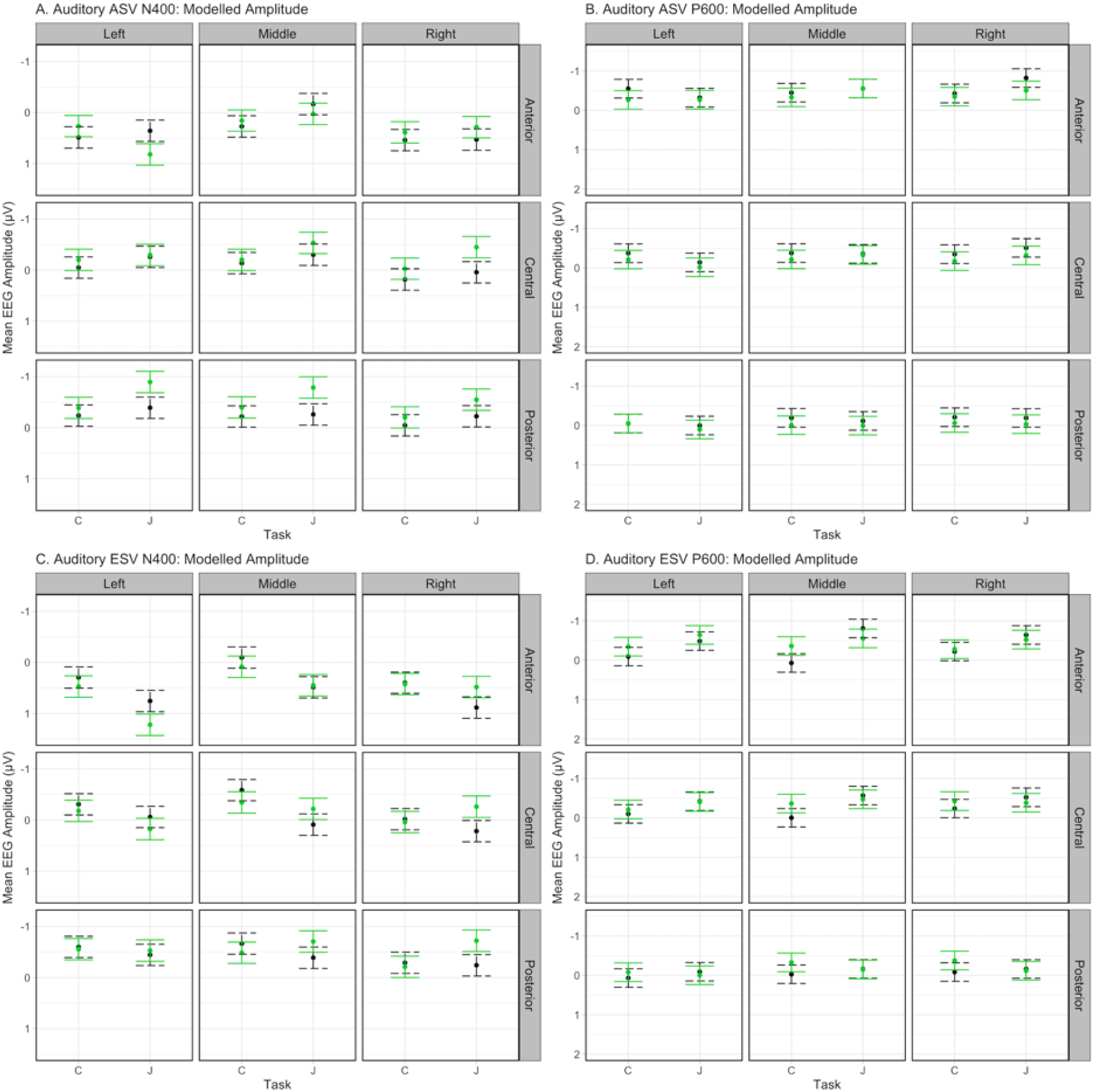
Interaction of sentence grammaticality, task type and region of interest on mean EEG amplitude for the Agent Subject Verb (ASV) and Experiencer Subject Verb (ESV) N400 and P600 time windows in the auditory experiment. Black/dashed lines represent correct/grammatical sentences while green/solid lines represent incorrect/ungrammatical sentences. Bars indicate 95% confidence intervals. Effects are shown for nine topographical regions of interest.

In the 700-900ms time-window, there was an interaction effect of Animacy x Verb Type x Task (χ^2^(1) =21.12, *p* <.001), as well as interactions of Verb Type x Task x Laterality (χ^2^(1) =18.44, *p* =.001) and Verb Type x Task x Sagitality C(^2^(1) =9.95, *p* =.007). Violations at the verb elicited a positivity for ASV, maximal in the right hemisphere and larger for the Judgement compared to the Comprehension condition. Visual inspection of the ERPs (Figure 7) show an ongoing negativity towards violation ESV sentences in the Comprehension group.

Prior to verb onset, visual inspection of the ERP data shows a difference in prestimulus amplitude between correct and violation sentences. To investigate this, we plotted the ERP data time-locked to onset of the Subject Noun and continuing past the onset of the verb. This showed that the prestimulus negativity prior to verb onset was the continuous negativity elicited by inanimate Subject Nouns, which was reinforced at the auxiliary. The ERP overlap across sentence constituents was expected, with similar phenomena observed in previous studies of auditory sentence processing (Choudhary et al., 2009; Wolff et al., 2008). See Appendix C for these ERP plots, and Appendix B for model summaries.

## 5. Experiment 2: Auditory modality summary

We replicated the behavioural results of Experiment 1’s MEM analysis. The ERP results at the Subject Noun also showed the same pattern as Experiment 1 where inanimate Subject Nouns elicited an N400 effect, larger for the Judgement condition. At the verb, no effects were found in the N400 time-window for the Comprehension condition. For the Judgement condition, we found an N400 effect to ASV and ESV violations, with visual inspection of the ERP plots showing it appeared larger for ESV. Violations at the verb also elicited a P600 effect for violation ASV sentences in both task groups. No significant effects were found in the verb P600 window for ESV.

## 6. Discussion

This paper aimed to reproduce the novel findings of Bourguignon et al. (2012), where verb class modulated the ERP response to English SRAs. Here, we investigated the effect of task demand and stimulus modality on the ERPs elicited by SRAs. We found that SRAs in English elicited both N400 and P600 effects at the critical verb position, and the N400 effect was elicited in both agent- and experiencer-subject verb types. Neither the reproduction ANOVA analysis nor the mixed effects model (MEM) analysis showed a P600 effect to ESV violations. Both task and modality modulated the ERPs elicited in a quantitative and qualitative manner.

### 6.1. Behavioural Results: Auditory input increases comprehension accuracy

Response accuracy was higher for the auditory over visual modality, Judgement over Comprehension task, ASV over ESV verb type, and grammatical over ungrammatical sentences. In the Judgement condition, a higher accuracy for ASV is likely due to violation saliency being stronger than in ESV, as suggested by Bourguignon et al. (2012). Comprehension task accuracy was much higher for the auditory compared to visual modality, with mean accuracy scores between 3.41% and 21.97% higher for auditory, following the modality-specific pattern of semantic anomalies reported by Bohan (2007), where accuracy was higher in response to auditory stimuli. When response accuracy is aggregated by item, variance was higher in the visual experiment compared to auditory, suggesting that additional (e.g. prosodic) cues in the auditory stimuli may have aided comprehension for items which were more difficult to interpret when presented visually (see Appendix D for by-subject and by-item variance plotted for each modality using the method proposed by Allen et al., 2019).

### 6.2. ERP results

#### 6.2.1. The subject animacy N400

As hypothesised, we observed an N400 effect for inanimate compared to animate Subject Nouns in Experiments 1 and 2. For Experiment 1 (visual), the reproduction analysis found the effect most prominent near the midline, while the MEM analysis found the effect right-lateralised. For Experiment 2 (auditory), the N400 was largest at posterior and central regions. The N400 effect likely reflects the increased processing cost incurred by an inanimate noun occurring in the position most commonly filled by the sentential subject in English (Bornkessel-Schlesewsky et al., 2011).

#### 6.2.2. The “verb type-independentN400”

For ASV and ESV, we observed an N400 effect at the verb for violation conditions, modulated by task and modality (see Table 8 for an overview of ERP effects across experiments). We had predicted an N400 for ESV violations, and replicated the results reported by Bourguignon et al. (2012), though not in the reproduction ANOVA analysis. Visual ESV violations elicited an N400 for the Comprehension condition only, while auditory ESV violations elicited an N400 in the Judgement condition only.

**Table 8.**
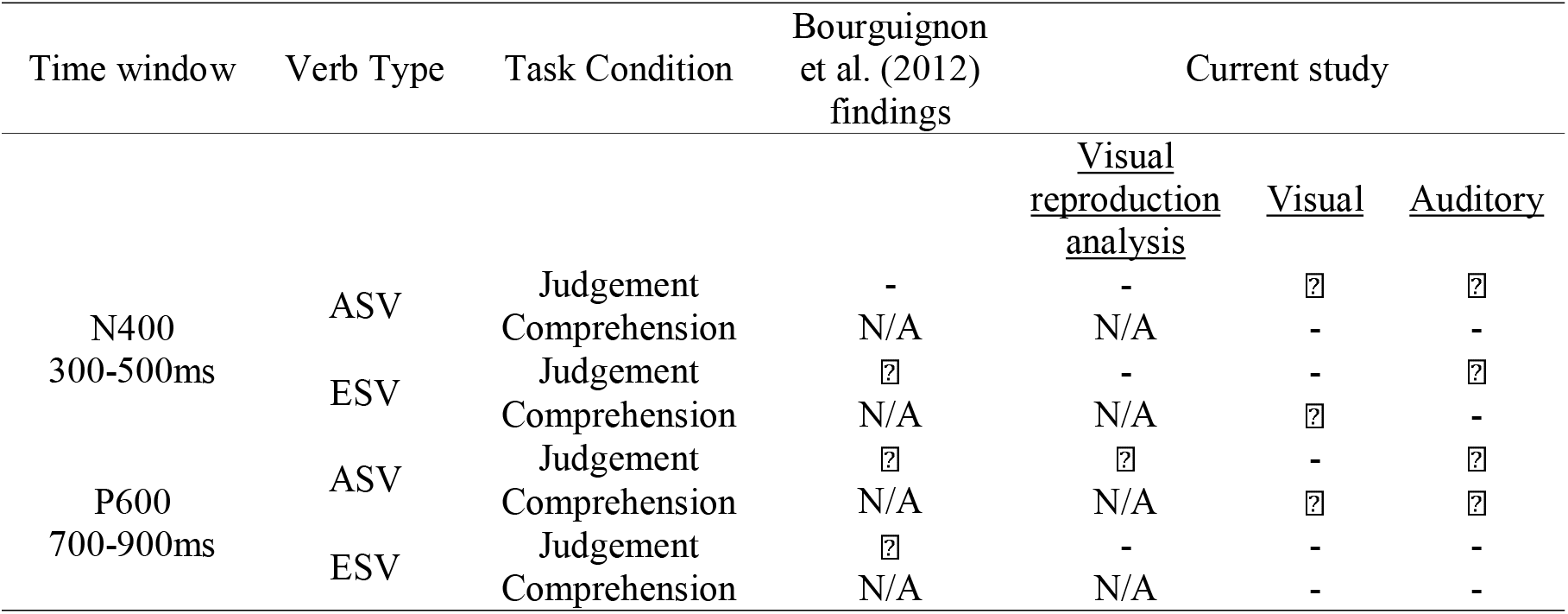
Overview of the presence/absence of significant ERP effects in the verb N400 and P600 time-windows for each verb type, task, modality, and analysis method.

| Time window | Verb Type | Task Condition | Bourguignon et al. (2012) findings | Current study |  |  |
| --- | --- | --- | --- | --- | --- | --- |
|  |  |  |  | <u>Visual reproduction analysis</u> | <u>Visual</u> | <u>Auditory</u> |
| N400<br>300-500ms | ASV | Judgement | - | - | ☐ | ☐ |
|  |  | Comprehension | N/A | N/A | - | - |
|  | ESV | Judgement | ☐ | - | - | ☐ |
|  |  | Comprehension | N/A | N/A | ☐ | - |
| P600<br>700-900ms | ASV | Judgement | ☐ | ☐ | - | ☐ |
|  |  | Comprehension | N/A | N/A | ☐ | ☐ |
|  | ESV | Judgement | ☐ | - | - | - |
|  |  | Comprehension | N/A | N/A | - | - |

We also observed an N400 effect to visual and auditory ASV violations in the Judgement condition, although inspection of the ERP plots suggests a smaller amplitude than the N400 to ESV violations. The presence of an N400 effect to an ASV violation was unexpected, as current theories were built upon findings of a monophasic P600 to similar English and Dutch SRAs (Hoeks et al., 2004; Kim & Osterhout, 2005; Kolk et al., 2003; Kuperberg et al., 2003; van Herten et al., 2006; Van Herten et al., 2005). While our findings of a “verb type-independent N400” to SRAs support Bornkessel-Schlesewsky et al.’s (2011) claim that monophasic P600 effects are not the only index of SRA processing, the presence of an N400 effect to typical SRA constructions in English falsifies all current theoretical approaches, including our own (Bornkessel-Schlesewsky et al., 2011). Bornkessel-Schlesewsky et al. (2011) propose a cross-linguistic perspective where cues to sentence interpretation vary in their dominance depending on the language in question. Our ASV N400 finding invalidates the hypothesis that the sequence-dependency of English, a language that relies primarily on word order for argument role assignments, necessarily leads to the absence of an N400 to SRAs.

A source of variance between the present study and Bourguignon et al. (2012) is the change from present perfect to future tense. While the difference in aspect resulting from this change may influence the “naturalness” of the sentences (which was the purpose of the change), it is not clear how this could explain the observed pattern of N400 effects. We observed N400 effects for ESV conditions - though not consistently across all modalities and tasks - even though the ESV sentences in the present study were more natural-sounding in tense. In regard to the more commonly examined ASV sentences, there is a high degree of diversity in the tense and aspect used in previous semantic reversal anomalies studies. In addition to the present perfect tense used by Bourguignon et al. (2012), this includes past progressive aspect (e.g. *The hearty meal was devouring…*, Kim & Osterhout, 2005) and conditional mood (e.g. *…the eggs would only eat.*, Kuperberg et al., 2003). These studies also did not find an N400 to agent subject verb violations, consistent with the findings of Bourguignon et al. (2012). While the use of future tense in the current study thus differs from previous research, it appears unlikely it would lead to these qualitative differences in the presence versus absence of N400 effects.

Another possibility is that some dialectal aspect of Australian English contributed to the differential ERP effects observed. To our knowledge, this is the first SRA investigation in Australian English, with all other English studies examining speakers of North American English (Bourguignon et al., 2012; Kim & Osterhout, 2005). Two observations support the idea that Australian English behaves differently to North American English: behavioural response differences, and recent findings of dialect-based ERP differences in German (Dröge et al., 2019). Compared to Bourguignon et al. (2012), our participants were 5.53% more likely to reject grammatical constructions, and 4.31% more likely to accept ungrammatical structures than North American participants. This suggests some meaningful difference in the way that Australian English speakers perceive the acceptability of SRA constructions. Dröge et al. (2019) found qualitative differences in the ERPs elicited by ambiguous object-first sentences across Standard German, Zurich German, and Fering (a North Frisian dialect). They found that the presence or absence of ERP effects was influenced by the most prominent cue to sentence interpretation in each dialect - either case marking or animacy (Dröge et al., 2019). With this recent finding of within-language differences depending on dialect, it is possible that dialectal differences in Australian English play a role in our results. However, we acknowledge that this is a post-hoc interpretation of our results that will require further investigation in future research.

#### 6.2.3. The “verb type-dependentP600”

A significant P600 effect was elicited by violation sentences for the ASV condition only. For the Judgement condition, a P600 was observed in the visual (reproduction analysis only) and auditory modalities. For the Comprehension condition, a P600 was observed in both modalities. These results fit with previous findings of a P600 to SRAs (although previous research reports only a monophasic P600 (Hoeks et al., 2004; Kim & Osterhout, 2005; Kolk et al., 2003; Kuperberg et al., 2003; van Herten et al., 2006; Van Herten et al., 2005).

The complete absence of a P600 to ESV violations was unexpected. While Bourguignon et al. (2012) found a P600 elicited by this verb type, their use of an Experiencer verb was novel. It is possible that ESV violations were less salient than violations in the filler sentences, leading to a diminished ERP response, as suggested by Bourguignon et al. (2012) and Steinhauer & Drury (2012).

#### 6.2.4. The quantitative and qualitative influence of task

We hypothesised (H2) a P600 at the Verb (with an Animacy effect). We predicted that the P600 effect would be larger for the Judgement task, due to the task-salience of the violation verb for grammaticality judgements (Schacht et al., 2014). Violations or perceived errors are still salient aspects for comprehension, so we expected a P600 for the Comprehension condition also, albeit with a smaller amplitude. While we predicted quantitative task-related differences, the observed qualitative differences were unexpected. In Experiment 1, there was a P600 effect in the Comprehension condition but no significant P600 for the Judgement condition, thus not supporting our hypothesis. However, we did see a significant P600 for the Judgement condition in the replication ANOVA analysis. In Experiment 2, we observed a P600 effect to auditory ASV violations in both task conditions, larger for the Judgement compared to Comprehension group, supporting our hypothesis.

Further, we found no evidence of a P600 to ESV violations for any modality or task, where we had predicted a P600 as reported by Bourguignon et al. (2012). In the 700-900ms time-window we observed a right-anterior negativity for ESV violations in the auditory Comprehension group. This is in line with the findings of Schacht et al. (2014) who found that by replacing a judgement task with a probe task, the P600 effect was replaced with a significant frontal negativity elicited by syntactic violations. They interpreted this negativity as representing semantic reinterpretation, which could fit with our results, although it does not explain the absence of a P600 in the auditory Judgement condition (Schacht et al., 2014).

#### 6.2.5. Modality influences comprehension and EEG topography

We hypothesised a difference in N400 topography based on modality, where stimuli presented auditorily would produce more broadly distributed ERP patterns. We had no hypotheses regarding behavioural responses, but Judgement and Comprehension accuracy were higher in the auditory experiment. This behavioural difference may be due to a facilitation of comprehension from the additional information in a natural speech stream. Speakers use disambiguating prosody in natural speech, and this information contributes to sentence comprehension (Marslen-Wilson, Tyler, Warren, Grenier, & Lee, 1992; Price, Ostendorf, Shattuck-Hufnagel, & Fong, 1998; Schafer, Speer, Warren, & White, 2000; Steinhauer, Alter, & Friederici, 1999). Prosody can influence sentence interpretation prior to the onset of an ambiguous phrase, aiding prediction of upcoming information (Snedeker & Trueswell, 2003). This auditory advantage was most pronounced for comprehension questions regarding ungrammatical (violation) sentences, suggesting prosodic information may have aided comprehension of semantically implausible constructions. While by-item variance in response accuracy was similar across modalities for the judgement task, the comprehension task responses showed a larger variance for the visual modality compared to less variance in the auditory modality (see Appendix E for plots). This may indicate that additional cues present in the auditory modality (e.g. prosody) facilitated the comprehension of specific items (sentences) that may have been more difficult to interpret in the visual modality.

At the Subject Noun, we observed a topographical difference in the animacy-based N400 effect, where the visual ERP was right-lateralised, spanning the middle and right columns. In the auditory modality, the N400 was spread across all columns in central and posterior sagitality. The results at the Subject Noun support our hypothesis that the ERP patterns would be more distributed in the auditory condition, and match with previous N400 topography findings (Holcomb & Neville, 1990).

At the verb it is difficult to compare N400 topography across modalities, as task and verb type influenced topography. Overall, for the auditory modality, the N400 was spread across more topographical regions than in the visual modality. At the Subject Noun, where differences are clearest, the N400 had a broader distribution across sagitality when stimuli were presented auditorily. At the Verb, the N400 to auditory ASV and ESV violations appears to be more broadly spread, encompassing a larger number of topographical regions.

#### 6.2.6. Statistical analysis decisions qualitatively affect results

There are two instances in our results where the analysis method used on the same data affects the presence/absence of a statistically significant ERP effect. Where the reproduction analysis using ANOVAs does not show an N400 elicited by ASV or ESV violations, the MEM analysis shows an N400 to ASV violations. Further, the reproduction analysis shows a significant P600 effect to ASV violations while the same visual data analysed with an MEM does not show these effects.

There are several potential explanations for these differences. Firstly, some have argued that high-pass filters over 0.1 Hz distort the data (Tanner, Norton, Morgan-Short, & Luck, 2016). The 0.4 Hz filter used in the reproduction may be more likely to introduce spurious components into the ERP waveform, compared to the relatively conservative 0.1 Hz filter used in the MEM analysis (Tanner et al., 2016). The influence of filter settings on these ERPs is unlikely however, as the MEM results of Experiment 2 displayed the two “missing” ERPS using a 0.1 Hz filter (presence of an N400 to ESV violations and P600 to ASV violations for judgement task). Secondly, the reproduction analysis was response-contingent, including only trials followed by a correct response. Though we did not expect this to have a significant effect on the presence/absence of the visual ESV N400 and ASV P600 in the Judgement condition, we conducted a response-contingent MEM for Experiment 1. The resultant MEMs for the time-windows of interest (Verb N400 and P600) showed no meaningful change from the original results.

The advantages of MEM analyses (and disadvantages of ANOVA) have recently been examined in detail by Kretzschmar and Alday (2020), who propose that MEMs be used as the default statistical analysis approach for electrophysiological and psycholinguistic data. The modelling of item-based variability is a major difference between ANOVA and MEM analyses. ANOVA analyses aggregate by participant and remove any variance caused by item variability. This is less than ideal, as each item in linguistic material (e.g. a word or sentence) is unique, even if the stimuli have been matched on parameters such as length or frequency (Janssen, 2012). On the other hand, MEM analyses do not aggregate, instead analysing trial-by-trial data while taking participant and item variability into account. Another key advantage of MEMs is that it is possible to include participants no matter the number of trials available due to the possibility of comparing unbalanced groups (Kretzschmar & Alday, 2020). For example, if after preprocessing EEG data, a small number of trials remain for a participant, this participant’s data can be included in the model rather than being excluded wholly as would be the case for a traditional ANOVA analysis. See Kretzschmar and Alday (2020) for an overview of MEM analyses and suggestions for application to EEG data.

#### 6.2.7. Item-based variability may drive differences in results

The sensitivity of MEMs to item variability may explain the absence of two ERP effects for the Judgement task when analysed using an MEM compared to an ANOVA. Regarding the ESV N400, the nature of ESV sentences, which lack the “concreteness” of ASV sentences, mean that the ERP response is likely to significantly vary across items. Item-specific effects such as the concreteness of word meaning or number and frequency of lexical associates have been shown to influence the amplitude of the N400 (Laszlo & Federmeier, 2011; Van Petten, 2014). Concrete, imageable words elicit a greater N400 amplitude than abstract words (Barber, Otten, Kousta, & Vigliocco, 2013; Gullick, Mitra, & Coch, 2013), thus the N400 amplitude in our study may have fluctuated depending on verb imageability (e.g. ESV “enjoy” may be more imageable than “regret”). Regarding the visual ASV P600, it is also likely that item-based variance is driving the lack of a significant P600 effect. Visual inspection of the ERP plots shows a positivity in the 700-900ms time-window, suggesting that the significant findings in the reproduction ANOVA are meaningful. Our data show larger item-based variance than subject-based variance on the intercept of the MEM (see Appendix F for an example). It is possible that the replication analysis is reflecting an N400 elicited by ESV violations and P600 elicited by ASV violations which do not occur consistently across items, and this inter-item variance is being considered by the MEM analysis.

In line with the current view that linguistic P600 effects may be at least partly driven by task-salience (Sassenhagen et al., 2014), we recommend that linguistic experiments integrate alternate tasks, such as probe or comprehension questions. Further, as it is not clear which methodological parameters most closely index the neural responses of interest, we suggest that future research should systematically compare the behavioural and ERP differences that arise across stimulus modalities, task types, and other experimental demands or contexts. Additionally, it appears that naturalistic auditory sentences increase the level of comprehension, so future research could investigate the extent to which this behavioural advantage continues outside of semantic anomalies. Finally, as shown by our direct comparison of ANOVAs and MEMs, the statistical analyses selected and their potential influence on ERP results, should be considered when developing future research.

## 7. Author contributions

LK, MS, and IB-S designed the research. LK conducted the research. LK and IB-S analysed the data. LK, MS and IB-S wrote the paper.

## 8. Acknowledgements

We would like to thank Zachariah Cross and Andrew Corcoran for helpful discussions, and Erica Wilkinson, Angela Osborn, and Nicole Vass for their assistance in data acquisition. We would like to thank Madeleine D’Angelo for recording the auditory stimuli. LK acknowledges the support of an Australian Government Research Training Program Scholarship (212791). IB-S acknowledges the support of an Australian Research Council Future Fellowship (FT160100437). We thank four anonymous reviewers for their helpful feedback.

## 9. List of Abbreviations

ASV: Agent subject verb
EEG: Electroencephalography
EOG: Electrooculography
EOV: Experiencer object verb
ERP: Event-Related Potential
ESV: Experiencer subject verb
FIR: Finite impulse response
MEM: Mixed effects (regression) model
PSV: Phrase structure violation
RSVP: Rapid serial visual presentation
SRA: Semantic Reversal Anomaly

## Footnotes

1 Two studies of Mandarin Chinese (Chow, Lau, Wang & Phillips, 2018; Chow & Phillips, 2013) report a monophasic P600 to SRAs in Mandarin Chinese, which appear to oppose the assertion that sequence-independent languages show an N400 to SRAs. These studies employ b□ constructions, where the coverb b□ denotes the NP1 as the actor and NP2 as the undergoer. However, these findings correspond to those reported by Bornkessel-Schlesewsky et al., (2011), who found a P600 for b□ but an N400 for bèi-constructions (which denote the NP1 as the undergoer and NP2 as the actor). Bornkessel-Schlesewsky et al., (2011) hypothesise that the absence of an N400 response to b□ SRAs may be due to the availability of an alternative relative clause analysis which is not possible for bèi SRAs.

2 Wassenaar & Hagoort (2007) employed a picture-matching task, with a picture presented prior to auditory sentence presentation in a control and clinical population. They observed an N400 - late positivity in control subjects to violations of semantically irreversible active and semantically reversible passive sentences. As the image was presented prior to the sentence, it is possible that the positivity observed was a task-related response when mapping the conceptualisation of the image onto the presented sentence.

3 For List 1: 19 Subject probes, 6 Object probes, 35 Yes/No probes; for List 2: 16 Subject probes, 6 Object probes, 38 Yes/No probes.

4 However, given that ASV and ESV are not directly compared at the Subject NP or auxiliary (instead, animate and inanimate Subject NPs are compared collapsed across conditions) the reported differences in duration are acceptable.

## References

Adams, R. A., Stephan, K. E., Brown, H. R., Frith, C. D., & Friston, K. J. (2013). The Computational Anatomy of Psychosis. Frontiers in Psychiatry, 4. https://doi.org/10.3389/fpsyt.2013.00047

Alday, P. M., Schlesewsky, M., & Bornkessel-Schlesewsky, I. (2017). Electrophysiology reveals the neural dynamics of naturalistic auditory language processing: event-related potentials reflect continuous model updates. Eneuro, 4(6).

Alday, P. M. (2018). lmerOut: LaTeX Output for Mixed Effects Models with lme4. R package version 0.5. https://bitbucket.org/palday/lmerout

Alday, P. M. (2019). How much baseline correction do we need in ERP research? Extended GLM model can replace baseline correction while lifting its limits. Psychophysiology, 56(12). https://doi.org/10.1111/psyp.13451

Allen, M., Poggiali, D., Whitaker, K., Marshall, T. R., & Kievit, R. A. (2019). Raincloud plots: A multi-platform tool for robust data visualization. Wellcome Open Research, 4, 63. https://doi.org/10.12688/wellcomeopenres.15191.1

Balconi, M., & Pozzoli, U. (2004). N400 and P600 or the role of the ERP correlates in sentence comprehension: Some applications to the Italian language. The Journal of General Psychology, 131(3), 268–303. https://doi.org/10.3200/genp.131.3.268-303

Barber, H. A., Otten, L. J., Kousta, S.-T., & Vigliocco, G. (2013). Concreteness in word processing: ERP and behavioral effects in a lexical decision task. Brain and Language, 125(1), 47–53. https://doi.org/10.1016/j.bandl.2013.01.005

Bates, D., Kliegl, R., Vasishth, S., & Baayen, H. (2015). Parsimonious mixed models. arXiv preprint arXiv:1506.04967.

Bates, D., Mächler, M., Bolker, B., & Walker, S. (2014). Fitting linear mixed-effects models using lme4. ArXiv Preprint ArXiv:1406.5823.

Bates, D., Maechler, M., Bolker, B., & Walker, S. (2015). Fitting Linear Mixed-Effects Models Using lme4. Journal of Statistical Software, 67(1), 1–48. doi:10.18637/jss.v067.i01.

Boersma, P. (2006). Praat: Doing phonetics by computer. Http://Www.Praat.Org/. Retrieved from http://ci.nii.ac.jp/naid/10017594077/

Bohan, J. (2008). Depth of processing and semantic anomalies (Doctoral Thesis, Department of Psychology, University of Glasgow, Glasgow, Scotland). Retrieved from http://theses.gla.ac.uk/1127/

Bornkessel-Schlesewsky, I., & Schlesewsky, M. (2013). Reconciling time, space and function: A new dorsal-ventral stream model of sentence comprehension. Brain and Language, 125(1), 60–76. https://doi.org/10.1016/j.bandl.2013.01.010

Bornkessel-Schlesewsky, I., & Schlesewsky, M. (2019). Toward a Neurobiologically Plausible Model of Language-Related, Negative Event-Related Potentials. Frontiers in Psychology, 10. https://doi.org/10.3389/fpsyg.2019.00298

Bornkessel-Schlesewsky, I., Kretzschmar, F., Tune, S., Wang, L., Genç, S., Philipp, M., … Schlesewsky, M. (2011). Think globally: Cross-linguistic variation in electrophysiological activity during sentence comprehension. Brain and Language, 117(3), 133–152. https://doi.org/10.1016/j.bandl.2010.09.010

Bornkessel-Schlesewsky, I., Schlesewsky, M., Small, S. L., & Rauschecker, J. P. (2015). Neurobiological roots of language in primate audition: Common computational properties. Trends in Cognitive Sciences, 19(3), 142–150. https://doi.org/10.1016/j.tics.2014.12.008

Bourguignon, N., Drury, J. E., Valois, D., & Steinhauer, K. (2012). Decomposing animacy reversals between agents and experiencers: An ERP study. Brain and Language, 122(3), 179–189. https://doi.org/10.1016/j.bandl.2012.05.001

Brennan, J., & Pylkkänen, L. (2010). Processing psych verbs: Behavioural and MEG measures of two different types of semantic complexity. Language and Cognitive Processes, 25(6), 777–807. https://doi.org/10.1080/01690961003616840

Brouwer, H., Fitz, H., & Hoeks, J. (2012). Getting real about semantic illusions: Rethinking the functional role of the P600 in language comprehension. Brain Research, 1446, 127–143. https://doi.org/10.1016/j.brainres.2012.01.055

Choudhary, K. K., Schlesewsky, M., Roehm, D., & Bornkessel-Schlesewsky, I. (2009). The N400 as a correlate of interpretively relevant linguistic rules: Evidence from Hindi. Neuropsychologia, 47(13), 3012–3022.

Chow, W. Y., & Phillips, C. (2013). No semantic illusions in the “Semantic P600” phenomenon: ERP evidence from Mandarin Chinese. Brain Research, 1506, 76–93.

Chow, W. Y., Lau, E., Wang, S., & Phillips, C. (2018). Wait a second! Delayed impact of argument roles on on-line verb prediction. Language, Cognition and Neuroscience, 33(7), 803–828.

Coulson, S., King, J. W., & Kutas, M. (1998). Expect the Unexpected: Event-related Brain Response to Morphosyntactic Violations. Language and Cognitive Processes, 13(1), 21–58. https://doi.org/10.1080/016909698386582

Domalski, P., Smith, M. E., & Halgren, E. (1991). Cross-Modal Repetition Effects on the N4. Psychological Science, 2(3), 173–178. 10.1111/j.1467-9280.1991.tb00126.x

Dröge, A., Rabs, E., Jürg, F., Billion, S. K. H., Meyer, M., Schmid, S., … Bornkessel-Schlesewsky, I. (2019). Case syncretism, animacy, and word order in Continental West Germanic: Neurolinguistic evidence from a comparative study on Standard German, Zurich German, and Fering (North Frisian). in press.

Feldman, H., & Friston, K. J. (2010). Attention, Uncertainty, and Free-Energy. Frontiers in Human Neuroscience, 4(215).

Fodor, J. A. (1983). The modularity of mind. MIT press.

Fox, J. & Weisberg, S. (2019). An {R} Companion to Applied Regression, Third Edition. Thousand Oaks CA: Sage. URL: https://socialsciences.mcmaster.ca/jfox/Books/Companion/

Friederici, A. D. (2011). The brain basis of language processing: From structure to function. Physiological Reviews, 91(4), 1357–1392. https://doi.org/10.1152/physrev.00006.2011

Friederici, A. D. (2012). The cortical language circuit: From auditory perception to sentence comprehension. Trends in Cognitive Sciences, 16(5), 262–268. https://doi.org/10.1016/j.tics.2012.04.001

Gramfort, A., Luessi, M., Larson, E., Engemann, D. A., Strohmeier, D., Brodbeck, C., … & Hämäläinen, M. S. (2014). MNE software for processing MEG and EEG data. Neuroimage, 86, 446–460.

Gullick, M. M., Mitra, P., & Coch, D. (2013). Imagining the truth and the moon: An electrophysiological study of abstract and concrete word processing. Psychophysiology, 50(5), 431–440. https://doi.org/10.1111/psyp.12033

Gunter, T. C., & Friederici, A. D. (1999). Concerning the automaticity of syntactic processing. Psychophysiology, 36(1), 126–137. https://doi.org/10.1017/s004857729997155x

Hagiwara, H., Soshi, T., Ishihara, M., & Imanaka, K. (2007). A topographical study on the event-related potential correlates of scrambled word order in Japanese complex sentences. Journal of Cognitive Neuroscience, 19(2), 175–193. 10.1162/jocn.2007.19.2.175

Hagoort, P., & Brown, C. M. (2000). ERP effects of listening to speech compared to reading: The P600/SPS to syntactic violations in spoken sentences and rapid serial visual presentation. Neuropsychologia, 38(11), 1531–1549. https://doi.org/10.1016/S0028-3932(00)00053-1

Hagoort, P., Brown, C., & Groothusen, J. (1993). The syntactic positive shift (SPS) as an ERP measure of syntactic processing. Language and Cognitive Processes, 8(4), 439–483. https://doi.org/10.1080/01690969308407585

Hoeks, J. C., Stowe, L. A., & Doedens, G. (2004). Seeing words in context: The interaction of lexical and sentence level information during reading. Cognitive Brain Research, 19(1), 59–73. https://doi.org/10.1016/j.cogbrainres.2003.10.022

Holcomb, P. J., & Neville, H. J. (1990). Auditory and visual semantic priming in lexical decision: A comparison using event-related brain potentials. Language and Cognitive Processes, 5(4), 281–312. https://doi.org/10.1080/01690969008407065

Holcomb, P. J., Coffey, S. A., & Neville, H. J. (1992). Visual and auditory sentence processing: A developmental analysis using event-related brain potentials. Developmental Neuropsychology, 8(2), 203–241. https://doi.org/10.1080/87565649209540525

Hosemann, J., Herrmann, A., Steinbach, M., Bornkessel-Schlesewsky, I., & Schlesewsky, M. (2013). Lexical prediction via forward models: N400 evidence from German Sign Language. Neuropsychologia, 51(11), 2224–2237. https://doi.org/10.1016/j.neuropsychologia.2013.07.013

Janssen, D. P. (2012). Twice random, once mixed: Applying mixed models to simultaneously analyze random effects of language and participants. Behavior Research Methods, 44(1), 232–247. https://doi.org/10.3758/s13428-011-0145-1

Kassambara, A. (2020). ggpubr: “ggplot2” Based Publication Ready Plots (Version 0.3.0) [Computer software]. https://CRAN.R-project.org/package=ggpubr

Kim, A., & Osterhout, L. (2005). The independence of combinatory semantic processing: Evidence from event-related potentials. Journal of Memory and Language, 52(2), 205–225. https://doi.org/10.1016/j.jml.2004.10.002

Kolk, H. H., Chwilla, D. J., Van Herten, M., & Oor, P. J. (2003). Structure and limited capacity in verbal working memory: A study with event-related potentials. Brain and Language, 85(1), 1–36. https://doi.org/10.1016/s0093-934x(02)00548-5

Kotz, S. A., Frisch, S., Von Cramon, D. Y., & Friederici, A. D. (2003). Syntactic language processing: ERP lesion data on the role of the basal ganglia. Journal of the International Neuropsychological Society, 9(7), 1053–1060. https://doi.org/10.1017/s1355617703970093

Kretzschmar, F., & Alday, P. M. (2020). Principles of statistical analyses: Old and new tools [Preprint]. PsyArXiv. https://doi.org/10.31234/osf.io/nyj3k

Kuperberg, G. R., Sitnikova, T., Caplan, D., & Holcomb, P. J. (2003). Electrophysiological distinctions in processing conceptual relationships within simple sentences. Cognitive Brain Research, 17(1), 117–129. https://doi.org/10.1016/s0926-6410(03)00086-7

Kutas, M., & Hillyard, S. A. (1980). Reading senseless sentences: Brain potentials reflect semantic incongruity. Science, 207(4427), 203–205. https://doi.org/10.1126/science.7350657

Kutas, M., Neville, H. J., & Holcomb, P. J. (1987). A preliminary comparison of the N400 response to semantic anomalies during reading, listening and signing. Electroencephalography and Clinical Neurophysiology Supplement, 39, 325–330.

Kutas, M., Van Petten, C. K., & Kluender, R. (2006). Psycholinguistics electrified II (1994-2005). In Handbook of psycholinguistics (pp. 659–724). Elsevier.

Laszlo, S., & Federmeier, K. D. (2011). The N400 as a snapshot of interactive processing: Evidence from regression analyses of orthographic neighbor and lexical associate effects. Psychophysiology, 48(2), 176–186. https://doi.org/10.1111/j.1469-8986.2010.01058.x

Lau, E. F., Phillips, C., & Poeppel, D. (2008). A cortical network for semantics: (De)constructing the N400. Nature Reviews Neuroscience, 9(12), 920–933. https://doi.org/10.1038/nrn2532

Lawrence, M. A. (2016). ez: Easy Analysis and Visualization of Factorial Experiments. R package version 4.4-0. https://CRAN.R-project.org/package=ez

MacWhinney, B., Bates, E., & Kliegl, R. (1984). Cue validity and sentence interpretation in English, German, and Italian. Journal of Verbal Learning and Verbal Behavior, 23(2), 127–150. https://doi.org/10.1016/s0022-5371(84)90093-8

Marslen-Wilson, W. D. (1975). Sentence perception as an interactive parallel process. Science, 189(4198), 226–228.

Marslen-Wilson, W. D., Tyler, L. K., Warren, P., Grenier, P., & Lee, C. S. (1992). Prosodic effects in minimal attachment. The Quarterly Journal of Experimental Psychology, 45(1), 73–87. https://doi.org/10.1080/14640749208401316

Neville, H. J., Coffey, S. A., Lawson, D. S., Fischer, A., Emmorey, K., & Bellugi, U. (1997). Neural systems mediating American Sign Language: Effects of sensory experience and age of acquisition. Brain and Language, 57(3), 285–308. https://doi.org/10.1006/brln.1997.1739

Neville, H. J., Mills, D. L., & Lawson, D. S. (1992). Fractionating language: Different neural subsystems with different sensitive periods. Cerebral Cortex, 2(3), 244–258. https://doi.org/10.1093/cercor/2.3.244

Osterhout, L., & Holcomb, P. J. (1992). Event-related brain potentials elicited by syntactic anomaly. Journal of Memory and Language, 31(6), 785–806. https://doi.org/10.1080/01690969308407584

Osterhout, L., & Holcomb, P. J. (1993). Event-related potentials and syntactic anomaly: Evidence of anomaly detection during the perception of continuous speech. Language and Cognitive Processes, 8(4), 413–437. https://doi.org/10.1016/0749-596x(92)90039-z

Osterhout, L., & McKinnon, R. (1996). On the language specificity of the brain response to syntactic anomalies: Is the syntactic… Journal of Cognitive Neuroscience, 8(6), 507. https://doi.org/10.1162/jocn.1996.8.6.507

Osterhout, L., Allen, M. D., Mclaughlin, J., & Inoue, K. (2002). Brain potentials elicited by prose-embedded linguistic anomalies. Mem Cogn, 30(8), 1304–1312. https://doi.org/10.3758/BF03213412

Payne, B. R., Lee, C. L., & Federmeier, K. D. (2015). Revisiting the incremental effects of context on word processing: Evidence from single-word event-related brain potentials. Psychophysiology, 52(11), 1456–1469.

Price, P. J., Ostendorf, M., Shattuck-Hufnagel, S., & Fong, C. (1998). The use of prosody in syntactic disambiguation. The Journal of the Acoustical Society of America, 90(6), 2956. https://doi.org/10.1121/L401770

R Core Team (2018). R: A language and environment for statistical computing. R Foundation for Statistical Computing, Vienna, Austria. URL https://www.R-project.org/

Roehm, D., Bornkessel-Schlesewsky, I., Rösler, F., & Schlesewsky, M. (2007). To predict or not to predict: Influences of task and strategy on the processing of semantic relations. Journal of Cognitive Neuroscience, 19(8), 1259–1274. https://doi.org/10.1162/jocn.2007.19.8.1259

Sassenhagen, J., Schlesewsky, M., & Bornkessel-Schlesewsky, I. (2014). The P600-as-P3 hypothesis revisited: Single-trial analyses reveal that the late EEG positivity following linguistically deviant material is reaction time aligned. Brain and Language, 137, 29–39. https://doi.org/10.1016/j.bandl.2014.07.010

Schacht, A., Sommer, W., Shmuilovich, O., Martíenz, P. C., & Martín-Loeches, M. (2014). Differential Task Effects on N400 and P600 Elicited by Semantic and Syntactic Violations. PLOS ONE, 9(3), e91226. https://doi.org/10.1371/journal.pone.0091226

Schafer, A. J., Speer, S. R., Warren, P., & White, S. D. (2000). Intonational Disambiguation in Sentence Production and Comprehension. Journal of Psycholinguistic Research, 29(2), 169–182. https://doi.org/10.1023/A:1005192911512

Snedeker, J., & Trueswell, J. (2003). Using prosody to avoid ambiguity: Effects of speaker awareness and referential context. Journal of Memory and Language, 48(1), 103–130. https://doi.org/10.1016/S0749-596X(02)00519-3

Steinhauer, K., & Drury, J. E. (2012). On the early left-anterior negativity (ELAN) in syntax studies. Brain and Language, 120(2), 135–162. https://doi.org/10.1016/j.bandl.2011.07.001

Steinhauer, K., Alter, K., & Friederici, A. D. (1999). Brain potentials indicate immediate use of prosodic cues in natural speech processing. Nature Neuroscience, 2(2), 191–196. https://doi.org/10.1038/5757

Stroud, C., & Phillips, C. (2012). Examining the evidence for an independent semantic analyzer: An ERP study in Spanish. Brain and Language, 120(2), 108–126. https://doi.org/10.1016/j.bandl.2011.02.001

Tanner, D., Norton, J. J. S., Morgan-Short, K., & Luck, S. J. (2016). On high-pass filter artifacts (they’re real) and baseline correction (it’s a good idea) in ERP/ERMF analysis. Journal of Neuroscience Methods, 266, 166–170. https://doi.org/10.1016/j.jneumeth.2016.01.002

The British National Corpus, version 3 (BNC XML Edition). (2007). Retrieved from http://www.natcorp.ox.ac.uk/

Van de Meerendonk, N., Kolk, H. H., Chwilla, D. J., & Vissers, C. T. W. (2009). Monitoring in language perception. Language and Linguistics Compass, 3(5), 1211–1224.

Van Herten, M., Chwilla, D. J., & Kolk, H. H. J. (2006). When Heuristics Clash with Parsing Routines: ERP Evidence for Conflict Monitoring in Sentence Perception. Journal of Cognitive Neuroscience, 18(7), 1181–1197. https://doi.org/10.1162/jocn.2006.18.7.1181

Van Herten, M., Kolk, H. H., & Chwilla, D. J. (2005). An ERP study of P600 effects elicited by semantic anomalies. Cognitive Brain Research, 22(2), 241–255. https://doi.org/10.1016/j.cogbrainres.2004.09.002

Van Petten, C. (2014). Examining the N400 semantic context effect item-by-item: Relationship to corpus-based measures of word co-occurrence. International Journal of Psychophysiology, 94(3), 407–419. https://doi.org/10.1016/j.ijpsycho.2014.10.012

Wickham, H. (2007). Reshaping Data with the reshape Package. Journal of Statistical Software, 21(12), 1–20. URL http://www.jstatsoft.org/v21/i12/.

Wickham, H. (2017). tidyverse: Easily Install and Load the ‘Tidyverse’. R package version 1.2.1. https://CRAN.R-project.org/package=tidyverse

Wickham, H., Chang, W., Henry, L., Pedersen, T. L., Takahashi, K., Wilke, C., Woo, K., Yutani, H., Dunnington, D., & RStudio. (2020). ggplot2: Create Elegant Data Visualisations Using the Grammar of Graphics (Version 3.3.0) [Computer software]. https://CRAN.R-project.org/package=ggplot2

Wolff, S., Schlesewsky, M., Hirotani, M., & Bornkessel-Schlesewsky, I. (2008). The neural mechanisms of word order processing revisited: Electrophysiological evidence from Japanese. Brain and Language, 107(2), 133–157. https://doi.org/10.1016/j.bandl.2008.06.003

