## Appendices for "Semantic Reversal Anomalies under the Microscope: Task and Modality Influences on Language-Associated Event-Related Potentials"

### Appendix A: Experimental Stimuli

**ASV Correct**
The parties will ratify the agreement at last.
The man will smoke the pipe at the camp.
The student will write the answer on the exam.
The teacher will explain the assessment very convincingly.
The children will hear the serenades during the night.
The civilians will barricade the shelters during the war.
The athletes will practice the workouts every day.
The medic will treat the wound for a long time.
The tourists will visit the pyramids for many years.
The gardener will plough the soil with his tools.
The ministers will tackle the difficulties in Parliament.
The enemies will sign the treaties in peace.
The hikers will use the compass in the forest.
The labourer will complete the work in the mine.
The workmen will inhale the fumes in the mine.
The customers will drink the beers in the pub.
The men will devour the meals in the restaurant.
The writers will compose the poems in the salon.
The hikers will consult the maps in the wilderness.
The campers will spend the holidays in the woods.
The villagers will flee the attacks near their homes.
The agents will record the whispers next door.
The tenants will overhear the argument next door.
The child will swallow the pill with hot tea.
The culprits will commit the crimes on the street.
The managers will incur the debts of the factory.
The nurse will bandage the scar on his back.
The committees will coordinate the tasks since the morning.
The postmen will send the letters to the border.

**ASV Violation**
The boys will eat the fries too quickly.
The agreement will ratify the parties at last.
The pipe will smoke the man at the camp.
The answer will write the student on the exam.
The assessment will explain the teacher very convincingly.
The serenades will hear the children during the night.
The shelters will barricade the civilians during the war.
The workouts will practice the athletes every day.
The wound will treat the medic for a long time.
The pyramids will visit the tourists for many years.
The soil will plough the gardener with his tools.
The difficulties will tackle the ministers in Parliament.
The treaties will sign the enemies in peace.
The compass will use the hikers in the forest.
The fumes will inhale the workmen in the mine.
The work will complete the labourer in the mine.
The beers will drink the customers in the pub.
The meals will devour the men in the restaurant.
The poems will compose the writers in the salon.
The maps will consult the hikers in the wilderness.
The holidays will spend the campers in the woods.
The attacks will flee the villagers near their homes.
The argument will overhear the tenants next door.
The whispers will record the agents next door.
The pill will swallow the child with hot tea.
The crimes will commit the culprits on the street.
The debts will incur the managers of the factory.
The scar will bandage the nurse on his back.
The tasks will coordinate the committees since the morning.
The letters will send the postmen to the border.
The fries will eat the boys too quickly.

**ESV Correct**
Her parents will appreciate my achievements a great deal.
His professors will support his efforts until the end.
The boys will relish the pastries all week long.
The citizens will hate the crimes and the politicians.
The kids will love the carousels at the fair.
The judges will despise the movies at the festival.
The jury will distrust the charges at the hearing.
Her husband will dislike her behaviour at the party.
The students will heed the reforms at the university.
The public will adore the music at those concerts.
The spectators will enjoy the jokes every evening.
The people will admire the inventions for a long time.
The patient will dread the operation for its risks.
The readers will cherish the poems for many centuries.
Her boss will value her work for many years.
The artist will resent the questions for some reason.
The girls will fear the storms for weeks.
The children will like the gifts of the orphanage.
The laureate will appreciate the prize at the ceremony
The customer will praise the service in that store.
The soldiers will mourn the losses in the army.
The students will lament the results in the competition.
The children will enjoy the holidays in the village.
The journalists will disdain the remarks of that newspaper.
The competitors will envy the successes of the company.
The managers will deplore the failures of the company.
The opponents will trust the proposals of the party.
The volunteers will regret the incidents on the ground.
Her boyfriend will grieve her death very much.
The pupils will dread the exams without reason.
My teachers will value my efforts a great deal.
The travellers will enjoy the journeys very much.
The victims will mourn the catastrophes in the area.
The boy will fear the wind at night.
The minister will dread the outbreak for long.
The detectives will suspect the clues at the investigation.
The believers will heed the sermons at the mass.
The brides will appreciate the compliments at the weddings.
My mother will dislike my comments during the meal.
The scientists will relish the challenge for its prestige.
The conductors will dread the rehearsals for many months.
The kids will enjoy the toys for many weeks.
Her neighbours will envy her achievements for no reason.
My folks will deplore my grades for the first time.
His auditors will cherish his songs for their beauty.
Her friends will disdain her handicaps for years.
The guests will praise the presents of the palace.
The tourists will like the drinks in the bar.
The students will admire the books in the class.
The doctors will regret the catastrophes in the country.
The researchers will trust the experiments in the other labs.
The fans will love the concerts in the park.
The veterans will grieve the losses of past wars.
The lawyers will resent the policies of the company.
The members will support the decisions of the party.
His supporters will hate his views once again.
Her brothers will appreciate my encouragements so much.
My children will adore my tales since birth.
The critics will despise the book very badly.
The team will lament the score very deeply.

**ESV Violation**
My achievements will appreciate her parents a great deal.
His efforts will support his professors until the end.
The pastries will relish the boys all week long.
The crimes will hate the citizens and the politicians.
The carousels will love the kids at the fair.
The movies will despise the judges at the festival.
The charges will distrust the jury at the hearing.
Her behaviour will dislike her husband at the party.
The reforms will heed the students at the university.
The music will adore the public at those concerts.
The jokes will enjoy the spectators every evening.
The inventions will admire the people for a long time.
The operation will dread the patient for its risks.
The poems will cherish the readers for many centuries.
Her work will value her boss for many years.
The questions will resent the artist for some reason.
The storms will fear the girls for weeks.
The gifts will like the children of the orphanage.
The prize will appreciate the laureate at the ceremony
The service will praise the customer in that store.
The losses will mourn the soldiers in the army.
The results will lament the students in the competition.
The holidays will enjoy the children in the village.
The remarks will disdain the journalists of that newspaper.

The failures will deplore the managers of the company.
The successes will envy the competitors of the company.
The proposals will trust the opponents of the party.
The incidents will regret the volunteers on the ground.
Her death will grieve her boyfriend very much.
The exams will dread the pupils without reason.
My efforts will value my teachers a great deal.
The journeys will enjoy the travellers very much.
The catastrophes will mourn the victims in the area.
The wind will fear the boy at night.
The outbreak will dread the minister for long.
The clues will suspect the detectives at the investigation.
The sermons will heed the believers at the mass.
The compliments will appreciate the brides at the weddings.
My comments will dislike my mother during the meal.
The challenge will relish the scientists for its prestige.
The rehearsals will dread the conductors for many months.
The toys will enjoy the kids for many weeks.
Her achievements will envy her neighbours for no reason.
My grades will deplore my folks for the first time.
His songs will cherish his auditors for their beauty.
Her handicaps will disdain her friends for years.
The presents will praise the guests of the palace.
The drinks will like the tourists in the bar.
The books will admire the students in the class.
The catastrophes will regret the doctors in the country.
The experiments will trust the researchers in the other labs.
The concerts will love the fans in the park.
The losses will grieve the veterans of past wars.
The policies will resent the lawyers of the company.
The decisions will support the members of the party.
His views will hate his supporters once again.
My encouragements will appreciate her brothers so much.
My tales will adore my children since birth.
The book will despise the critics very badly.
The score will lament the team very deeply.

**EOV Correct**

My achievements will gladden her parents a great deal.
His efforts will content his professors until the end.
The pastries will tempt the boys all week long.
The crimes will appal the citizens and the politicians.
The carousels will thrill the kids at the fair.
The movies will displease the judges at the festival.
The charges will bother the jury at the hearing.
Her behaviour will embarrass her husband at the party.
The reforms will impress the students at the university.
The music will enchant the public at those concerts.
The jokes will amuse the spectators every evening.
The inventions will fascinate the people for a long time.
The operation will disquiet the patient for its risks.
The poems will seduce the readers for many centuries.
Her work will satisfy her boss for many years.
The questions will upset the artist for some reason.
The storms will frighten the girls for weeks.
The gifts will please the children of the orphanage.
The prize will cheer the laureate at the ceremony
The service will delight the customer in that store.
The losses will afflict the soldiers in the army.
The results will demoralise the students in the competition.
The holidays will excite the children in the village.
The remarks will repulse the journalists of that newspaper.
The failures will disappoint the managers of the company.
The successes will frustrate the competitors of the company.
The proposals will convince the opponents of the party.
The incidents will depress the volunteers on the ground.
Her death will affect her boyfriend very much.
The exams will torment the pupils without reason.
My efforts will satisfy my teachers a great deal.
The journeys will excite the travellers very much.
The catastrophes will afflict the victims in the area.
The wind will frighten the boy at night.
The outbreak will disquiet the minister for long.
The clues will bother the detectives at the investigation.
The sermons will impress the believers at the mass.
The compliments will cheer the brides at the weddings.
My comments will embarrass my mother during the meal.
The challenge will tempt the scientists for its prestige.
The rehearsals will torment the conductors for many months.
The toys will amuse the kids for many weeks.
Her achievements will frustrate her neighbours for no reason.
My grades will disappoint my folks for the first time.
His songs will seduce his auditors for their beauty.
Her handicaps will repulse her friends for years.
The presents will delight the guests of the palace.
The drinks will please the tourists in the bar.
The books will fascinate the students in the class.
The catastrophes will depress the doctors in the country.
The experiments will convince the researchers in the other labs.
The concerts will thrill the fans in the park.
The losses will affect the veterans of past wars.
The policies will upset the lawyers of the company.
The decisions will content the members of the party.
His views will appal his supporters once again.
My encouragements will gladden her brothers so much.
My tales will enchant my children since birth.
The book will displease the critics very badly.
The score will demoralise the team very deeply.

**EOV Violation**

Her parents will gladden my achievements a great deal.
His professors will content his efforts until the end.
The boys will tempt the pastries all week long.
The citizens will appal the crimes and the politicians.
The kids will thrill the carousels at the fair.
The judges will displease the movies at the festival.
The jury will bother the charges at the hearing.
Her husband will embarrass her behaviour at the party.
The students will impress the reforms at the university.
The public will enchant the music at those concerts.
The spectators will amuse the jokes every evening.
The people will fascinate the inventions for a long time.
The patient will disquiet the operation for its risks.
The readers will seduce the poems for many centuries.
Her boss will satisfy her work for many years.
The artist will upset the questions for some reason.
The girls will frighten the storms for weeks.
The children will please the gifts of the orphanage.
The laureate will cheer the prize at the ceremony
The customer will delight the service in that store.
The soldiers will afflict the losses in the army.
The students will demoralise the results in the competition.
The children will excite the holidays in the village.
The journalists will repulse the remarks of that newspaper.
The competitors will frustrate the successes of the company.
The managers will disappoint the failures of the company.
The opponents will convince the proposals of the party.
The volunteers will depress the incidents on the ground.
Her boyfriend will affect her death very much.
The pupils will torment the exams without reason.
My teachers will satisfy my efforts a great deal.
The travellers will excite the journeys very much.
The victims will afflict the catastrophes in the area.
The wind will frighten the wind at night.
The minister will disquiet the outbreak for long.
The detectives will bother the clues at the investigation.
The believers will impress the sermons at the mass.
The brides will cheer the compliments at the weddings.
My mother will embarrass my comments during the meal.
The scientists will tempt the challenge for its prestige.
The conductors will torment the rehearsals for many months.
The kids will amuse the toys for many weeks.
Her neighbours will frustrate her achievements for no reason.
My folks will disappoint my grades for the first time.
His auditors will seduce his songs for their beauty.
Her friends will repulse her handicaps for years.
The guests will delight the presents of the palace.
The tourists will please the drinks in the bar.
The students will fascinate the books in the class.
The doctors will depress the catastrophes in the country.
The researchers will convince the experiments in the other labs.
The fans will thrill the concerts in the park.

The veterans will affect the losses of past wars.
The lawyers will upset the policies of the company.
The members will content the decisions of the party.
His supporters will appal his views once again.
Her brothers will gladden my encouragements so much.
My children will enchant my tales since birth.
The critics will displease the book very badly.
The team will demoralise the score very deeply.
**PSV Correct**
It will be hard to survive this August without water.
It will be hard this August to survive without water.
The university wants to publish the student this semester.
The university wants the student to publish this semester.
The general tried to hide the army from the enemy.
The general asked the army to hide from the enemy.
The actor hoped to improve the music in his scene.
The actor chose his music to improve his scene.
The priest arranged to marry the teachers at the school.
The priest wanted the teachers to marry at the school.
The people began to prepare the city for the election.
The people decorated the city to prepare for the election.
He started to thank his wife at the ceremony.
He asked his wife to thank the driver.
The man chose to adopt the rabbit for his kids.
The man chose the rabbit to adopt for his kids.
The officer tried to commit the lady for the crime.
The man forced the lady to commit the crime.
The girl wanted to collect a bird from the woods.
The girl helped the bird to collect a pile of sticks.
This letter was written to replace the letter he wrote yesterday.
He wrote this letter to replace the one from yesterday.
The teacher hated to fail the children in math.
The teacher hated for the children to fail in math.
He wanted to begin this story with a poem.
He wanted this story to begin in the morning.
The fence worked to refuse the horse entry to the field.
The farmer expected the horse to refuse to drink.
The boss tried to explain his method to the workers.
Their boss found one method to explain the huge loss.
We were asked to join the lieutenant up with a partner.
Everyone wanted the lieutenant to join their department.
The manager wants to reflect their product in the company name.
The company expects the product to reflect their values.
My father hopes to grow a tree in his yard.
My father hopes for a tree to grow in his yard.
The director works to inspect the restaurant in the mornings.
The director entered the restaurant to inspect the kitchen.
The writer expected to vary the ending of his film.
The audience expects the ending to vary every week.
She waited to open her business until after Christmas.
She waited for the business to open the new store.
The players were known to select a leader who was strong.
The players wanted their leader to select a practice time.
I would never try to threaten a friendship between two people.
I expect their friendship to threaten our time together.
I have to admit my strength in sport is weak.
I have the strength to admit when I am wrong.
I want to ignore the news while I eat.
I want the news to ignore the scandal.
She wanted to compare this trial with an earlier one.
She ran this trial to compare our performances.
She wants me to prefer her cat to mine.
She wants her cat to prefer her company to mine.
My parents want to discuss my choice of university.
I made the choice to discuss my career change.
I came to manage the department and its courses.
I came to the department to manage these courses.
We came to greet the owner of the park.
We asked the owner to greet our employees.
**PSV Violation**
It will be hard to August this survive without water.
It will be hard this survive to August without water.
The university wants to student the publish this semester.
The university wants the publish to student this semester.
The general tried to army the hide from the enemy.
The general asked the hide to army from the enemy.
The actor hoped to music the improve in his scene.
The actor chose his improve to music his scene.
The priest arranged to teachers the marry at the school.
The priest wanted the marry to teachers at the school.
The people began to city the prepare for the election.
The people decorated the prepare to city for the election.
He started to wife his thank at the ceremony.
He asked his thank to wife the driver.
The man chose to rabbit the adopt for his kids.
The man chose the adopt to rabbit for his kids.
The officer tried to lady the commit for the crime.
The man forced the commit to lady the crime.
The girl wanted to bird a collect from the woods.
The girl helped the collect to bird a pile of sticks.
This letter was written to letter the replace he wrote yesterday.
He wrote this replace to letter the one from yesterday.
The teacher hated to children the fail in math.
The teacher hated for the fail to children in math.
He wanted to story this begin with a poem.
He wanted this begin to story in the morning.
The fence worked the refuse to horse entry to the field.
The farmer expected the refuse to horse to drink.
The boss tried to method his explain to the workers.
Their boss found one explain to method the huge loss.
We were asked to lieutenant the join up with a partner.
Everyone wanted the join to lieutenant their department.
The manager wants to product their reflect in the company name.
The company expects the reflect to product their values.
My father hopes to tree a grow in his yard.
My father hopes for a grow to tree in his yard.
The director works to restaurant the inspect in the mornings.
The director entered the inspect to restaurant the kitchen.
The writer expected to ending the vary of his film.
The audience expects the vary to ending every week.
She waited to business her open until after Christmas.
She waited for the open to business the new store.
The players were known to leader a select who was strong.
The players wanted their select to leader a practice time.
I would never try to friendship a threaten between two people.
I expect their threaten to friendship our time together.
I have to strength my admit in sport is weak.
I have the admit to strength when I am wrong.
I want to news the ignore while I eat.
I want the ignore to news the scandal.
She wanted to trial this compare with an earlier one.
She ran this compare to trial our performances.
She wants me to cat her prefer to mine.
She wants her prefer to cat her company to mine.
My parents want to choice my discuss of university.
I made the discuss to choice my career change.
I came to department the manage and its courses.
I came to the manage to department these courses.
We came to owner the greet of the park.
We asked the greet to owner our employees.
**SEA Correct**
The senate will abolish the laws on taxes.
The cook will mince the apples with a knife.
The company will achieve the budget in no time.
The gardener will plant the tomatoes in no time.
The committee will cancel the meeting of next week.
The staff will mend the cracks in the wall.
The bank will invest the money into technology.
The farmer will break the eggs in the henhouse.
The crew will compute the trajectories of the aircraft.
The peasants will peel the carrots in the kitchen.
The crowd will witness the incidents on the street.
The apprentice will grind the seeds to make flour.
The class will ask the questions before the exam.
The housemaid will wash the laundry in the sink.
The orchestra will compose the sonatas for a soprano.
The lady will sweep the floor before the party.
The operators will dial the numbers on the list.

The librarians will shelve the books on the list.
The seller will endorse the expenses of the purchase.
The mechanic will unpack the instruments in the workshop.
The troupe will recite the prose at the show.
The roadie will pack the spotlight for the show.
The ministry will spare the expenses quite drastically.
The technician will empty the battery for the winter.
The Parliament will endorse the policy on traffic.
The chef will season the salads with vinegar.
The press will condemn the murders on television.
The florist will sell the roses on the market.
The secretary will schedule the appointments in the afternoon.
The barman will fill the glasses with water.
The jury will explain the exercises to the class.
The caretaker will dust the sofas in the hall.
The team will solve the equations very skilfully.
The mason will cement the bricks very skilfully.
The society will finance the costs in the long run.
The artist will photograph the stars in the sky.
The police will investigate the murders for a week.
The maid will clean the kettles with a sponge.
The government will violate the agreements on purpose.
The musician will play the piano on stage.
The laboratory will calculate the measurements with a computer.
The professor will consult the dictionaries in the library.
The factory will cancel the process too early.
The mother will salt the beans too early.
The troops will plan the attacks on the enemy.
The women will boil the potatoes with garlic.
The party will supervise the negotiations of the ministers.
The shoemaker will polish the boots with a cloth.
The convent will dedicate the prayers to the poor.
The electrician will repair the telephones in the house.
The prison will acquit the crimes of the man.
The sleeper will stuff the feathers in the pillows.
The studio will record the songs in a few days.
The barmaid will squeeze the oranges to make punch.
The prompter will mouth the texts to the actors.
The baker will cook the cakes in the oven.
The philosopher will interpret the ideas very badly.
The labourer will glue the wallpaper very badly.
The senate will correct the mistakes in the document.
The walker will pick the strawberries in the woods.

**SEA Violation**

The senate will abolish the apples on taxes.
The cook will mince the laws with a knife.
The company will achieve the tomatoes in no time.
The gardener will plant the budget in no time.
The committee will cancel the cracks of next week.
The staff will mend the meeting in the wall.
The bank will invest the eggs into technology.
The farmer will break the money in the henhouse.
The crew will compute the carrots of the aircraft.
The peasants will peel the trajectories in the kitchen.
The crowd will witness the seeds on the street.
The apprentice will grind the incidents to make flour.
The class will ask the clothes before the exam.
The housemaid will wash the questions in the sink.
The orchestra will compose the floor for a soprano.
The lady will sweep the sonatas before the party.
The operators will dial the books on the list.
The librarians will shelve the numbers on the list.
The seller will endorse the instruments of the purchase.
The mechanic will unpack the expenses in the workshop.
The troupe will recite the spotlight at the show.
The roadie will pack the prose for the show.
The ministry will spare the battery quite drastically.
The technician will empty the expenses for the winter.
The Parliament will endorse the salads on traffic.
The chef will season the policy with vinegar.
The press will condemn the roses on television.
The florist will sell the murders on the market.
The secretary will schedule the glasses in the afternoon.
The barman will fill the appointments with water.
The jury will explain the sofas to the class.
The caretaker will dust the exercises in the hall.
The team will solve the bricks very skilfully.
The mason will cement the equations very skilfully.
The society will finance the stars in the long run.
The artist will photograph the costs in the sky.
The police will investigate the kettles for a week.
The maid will clean the murders with a sponge.
The government will violate the piano on purpose.
The musician will play the agreements on stage.
The laboratory will calculate the dictionaries with a computer.
The professor will consult the measurements in the library.
The factory will cancel the beans too early.
The mother will salt the process too early.
The troops will plan the potatoes on the enemy.
The women will boil the attacks with garlic.
The party will supervise the boots of the ministers.
The shoemaker will polish the negotiations with a cloth.
The convent will dedicate the telephones to the poor.
The electrician will repair the prayers in the house.
The prison will acquit the feathers of the man.
The sleeper will stuff the crimes in the pillows.
The studio will record the oranges in a few days.
The barmaid will squeeze the songs to make punch.
The prompter will mouth the cakes to the actors.
The baker will cook the texts in the oven.
The philosopher will interpret the wallpaper very badly.
The labourer will glue the ideas very badly.
The senate will correct the strawberries in the document.
The walker will pick the mistakes in the woods.

#### Appendix B: Model Summaries for the Mixed Effects Models conducted in Experiments 1 and 2.

Note. Categorical variables were encoded with sum encoding (ANOVA-style coding) where the model coefficient represents the size of the contrast from a given predictor level to the grand mean. For a two-level predictor, this is half the distance between the two levels, as the mean is an equal distance from both points.

Table S1: Visual study: Judgement behavioural model.

Generalized linear mixed model fit by maximum likelihood (Laplace Approximation)
Family: binomial Link:logit

| | AIC | BIC | logLik | deviance | | |
|---|---|---|---|---|---|---|
| | 401269 | 401325 | -200630 | 401259 | | |
| Scaled residuals: | | | | | | |
| | Min | 1Q | Median | 3Q | Max | |
| | -4.63 | 0.25 | 0.36 | 0.45 | 0.68 | |
| Random effects: | | | | | | |
| | Groups | Name | Std.Dev. | | | |
| | subj | (Intercept) | 0.43793 | | | |
| Number of obs: 509112, groups: subj, 22. | | | | | | |
| Fixed effects: | | | | | | |
| | | Estimate | Std. Error | z value | Pr(>\|z\|) | |
| (Intercept) | | 1.9 | 0.099 | 19 | 1.8e−81 | *** |
| animacy[Grammatical] | | −0.071 | 0.0042 | −17 | 1.6e−65 | *** |
| verbtype[ASV] | | 0.33 | 0.0042 | 78 | 0 | *** |
| animacy[Grammatical]:verbtype[ASV] | | 0.0093 | 0.0042 | 2.2 | 0.026 | * |

Table S2: Visual study: Comprehension behavioural model.

Generalized linear mixed model fit by maximum likelihood (Laplace Approximation)
Family: binomial Link:logit

| | | | | | | |
|---|---|---|---|---|---|---|
| | AIC | BIC | logLik | deviance | | |
| | 403323 | 403420 | -201652 | 403305 | | |
| Scaled residuals: | | | | | | |
| | Min | 1Q | Median | 3Q | Max | |
| | -4.99 | -0.95 | 0.44 | 0.71 | 1.76 | |
| Random effects: | | | | | | |
| | Groups | Name | Std.Dev. | | | |
| | subj | (Intercept) | 0.77078 | | | |
| Number of obs: 355320, groups: subj, 23. | | | | | | |
| Fixed effects: | | | | | | |
| | | Estimate | Std. Error | z value | Pr(>\|z\|) | |
| | (Intercept) | 0.85 | 0.14 | 6 | 2.2e−09 | *** |
| | animacy[Grammatical] | 0.37 | 0.004 | 94 | 0 | *** |
| | verbtype[ASV] | 0.1 | 0.0039 | 26 | 3.7e−148 | *** |
| | qtype[Actor Probe] | 0.0016 | 0.0039 | 0.4 | 0.69 | |
| | animacy[Grammatical]:verbtype[ASV] | −0.027 | 0.0039 | −6.9 | 5.9e−12 | *** |
| | animacy[Grammatical]:qtype[Actor Probe] | −0.31 | 0.004 | −78 | 0 | *** |
| | verbtype[ASV]:qtype[Actor Probe] | −0.089 | 0.0043 | −21 | 7.1e−96 | *** |
| | animacy[Grammatical]:verbtype[ASV]:qtype[Actor Probe] | −0.082 | 0.0039 | −21 | 1.6e−96 | *** |

Table S3: Visual study: Noun 1 N400 model.

Linear mixed model fit by REML

REML criterion at convergence: 682415

Scaled residuals:

| Min | 1Q | Median | 3Q | Max |
|---|---|---|---|---|
| -11.85 | -0.58 | 0.02 | 0.6 | 14.41 |

Random effects:

| Groups | Name | Std.Dev. |
|---|---|---|
| item | (Intercept) | 0.088337 |
| subj | (Intercept) | 0.198157 |
| Residual | | 0.960517 |

Number of obs: 247212, groups: item, 60; subj, 45.

Fixed effects:

| | Estimate | Std. Error | t value |
|---|---|---|---|
| (Intercept) | 0.0028 | 0.032 | 0.087 |
| animacy[Animate] | 0.023 | 0.0019 | 12 |
| task[C] | −0.011 | 0.03 | −0.36 |
| lat.[L] | 0.1 | 0.0027 | 38 |
| lat.[R] | 0.025 | 0.0027 | 9.1 |
| sag.[ant] | −0.044 | 0.0027 | −16 |
| sag.[post] | 0.07 | 0.0027 | 25 |
| scale(prestim) | −0.16 | 0.002 | −78 |
| animacy[Animate]:task[C] | −0.021 | 0.0019 | −11 |
| animacy[Animate]:lat.[L] | −0.012 | 0.0027 | −4.4 |
| animacy[Animate]:lat.[R] | 0.0023 | 0.0027 | 0.84 |
| task[C]:lat.[L] | 0.0012 | 0.0027 | 0.43 |
| task[C]:lat.[R] | −0.0083 | 0.0027 | −3 |
| animacy[Animate]:sag.[ant] | 0.0013 | 0.0027 | 0.48 |
| animacy[Animate]:sag.[post] | −0.0036 | 0.0027 | −1.3 |
| task[C]:sag.[ant] | 0.0013 | 0.0027 | 0.48 |
| task[C]:sag.[post] | 0.0022 | 0.0027 | 0.81 |
| lat.[L]:sag.[ant] | −0.01 | 0.0039 | −2.7 |
| lat.[R]:sag.[ant] | −0.019 | 0.0039 | −4.8 |
| lat.[L]:sag.[post] | −0.01 | 0.0039 | −2.6 |
| lat.[R]:sag.[post] | 0.0082 | 0.0039 | 2.1 |
| animacy[Animate]:task[C]:lat.[L] | 0.0045 | 0.0027 | 1.6 |
| animacy[Animate]:task[C]:lat.[R] | 0.0022 | 0.0027 | 0.79 |
| animacy[Animate]:task[C]:sag.[ant] | 0.0039 | 0.0027 | 1.4 |
| animacy[Animate]:task[C]:sag.[post] | −0.0012 | 0.0027 | −0.42 |
| animacy[Animate]:lat.[L]:sag.[ant] | −0.0041 | 0.0039 | −1.1 |
| animacy[Animate]:lat.[R]:sag.[ant] | 0.0063 | 0.0039 | 1.6 |
| animacy[Animate]:lat.[L]:sag.[post] | 0.0023 | 0.0039 | 0.59 |
| animacy[Animate]:lat.[R]:sag.[post] | −0.0022 | 0.0039 | −0.57 |
| task[C]:lat.[L]:sag.[ant] | 0.004 | 0.0039 | 1 |
| task[C]:lat.[R]:sag.[ant] | 0.0049 | 0.0039 | 1.3 |
| task[C]:lat.[L]:sag.[post] | −0.01 | 0.0039 | −2.6 |
| task[C]:lat.[R]:sag.[post] | 0.0074 | 0.0039 | 1.9 |
| animacy[Animate]:task[C]:lat.[L]:sag.[ant] | −0.0052 | 0.0039 | −1.4 |
| animacy[Animate]:task[C]:lat.[R]:sag.[ant] | 0.0016 | 0.0039 | 0.42 |
| animacy[Animate]:task[C]:lat.[L]:sag.[post] | 0.0012 | 0.0039 | 0.31 |
| animacy[Animate]:task[C]:lat.[R]:sag.[post] | −0.00099 | 0.0039 | −0.26 |

Table S4: Visual study: Noun 1 P600 model.

Linear mixed model fit by REML

REML criterion at convergence: 627543

Scaled residuals:

| Min | 1Q | Median | 3Q | Max |
|---|---|---|---|---|
| -11.4 | -0.59 | -0.01 | 0.59 | 8.72 |

Random effects:

| Groups | Name | Std.Dev. |
|---|---|---|
| item | (Intercept) | 0.069131 |
| subj | (Intercept) | 0.162508 |
| Residual | | 0.859645 |

Number of obs: 247212, groups: item, 60; subj, 45.

Fixed effects:

| | Estimate | Std. Error | t value |
|---|---|---|---|
| (Intercept) | −0.0034 | 0.026 | −0.13 |
| animacy[Animate] | −0.00078 | 0.0017 | −0.45 |
| task[C] | −0.035 | 0.024 | −1.5 |
| lat.[L] | −0.042 | 0.0024 | −17 |
| lat.[R] | −0.023 | 0.0024 | −9.5 |
| sag.[ant] | 0.025 | 0.0025 | 10 |
| sag.[post] | −0.042 | 0.0025 | −17 |
| scale(prestim) | −0.48 | 0.0018 | −2.7e+02 |
| animacy[Animate]:task[C] | 0.012 | 0.0017 | 6.7 |
| animacy[Animate]:lat.[L] | 0.0022 | 0.0024 | 0.89 |
| animacy[Animate]:lat.[R] | −0.0006 | 0.0024 | −0.25 |
| task[C]:lat.[L] | 0.0066 | 0.0024 | 2.7 |
| task[C]:lat.[R] | 0.021 | 0.0024 | 8.7 |
| animacy[Animate]:sag.[ant] | −0.0035 | 0.0024 | −1.4 |
| animacy[Animate]:sag.[post] | 0.0044 | 0.0024 | 1.8 |
| task[C]:sag.[ant] | −0.0044 | 0.0024 | −1.8 |
| task[C]:sag.[post] | 0.01 | 0.0024 | 4.2 |
| lat.[L]:sag.[ant] | 0.0027 | 0.0035 | 0.78 |
| lat.[R]:sag.[ant] | 0.00079 | 0.0035 | 0.23 |
| lat.[L]:sag.[post] | 0.014 | 0.0035 | 3.9 |
| lat.[R]:sag.[post] | 0.0069 | 0.0035 | 2 |
| animacy[Animate]:task[C]:lat.[L] | −0.0067 | 0.0024 | −2.7 |
| animacy[Animate]:task[C]:lat.[R] | 0.0052 | 0.0024 | 2.1 |
| animacy[Animate]:task[C]:sag.[ant] | 0.0046 | 0.0024 | 1.9 |
| animacy[Animate]:task[C]:sag.[post] | −0.0064 | 0.0024 | −2.6 |
| animacy[Animate]:lat.[L]:sag.[ant] | −0.003 | 0.0035 | −0.87 |
| animacy[Animate]:lat.[R]:sag.[ant] | 0.0027 | 0.0035 | 0.79 |
| animacy[Animate]:lat.[L]:sag.[post] | 0.0023 | 0.0035 | 0.66 |
| animacy[Animate]:lat.[R]:sag.[post] | −0.0017 | 0.0035 | −0.49 |
| task[C]:lat.[L]:sag.[ant] | 0.01 | 0.0035 | 3 |
| task[C]:lat.[R]:sag.[ant] | −0.013 | 0.0035 | −3.8 |
| task[C]:lat.[L]:sag.[post] | −0.016 | 0.0035 | −4.6 |
| task[C]:lat.[R]:sag.[post] | 0.013 | 0.0035 | 3.8 |
| animacy[Animate]:task[C]:lat.[L]:sag.[ant] | −0.0021 | 0.0035 | −0.6 |
| animacy[Animate]:task[C]:lat.[R]:sag.[ant] | 0.002 | 0.0035 | 0.57 |
| animacy[Animate]:task[C]:lat.[L]:sag.[post] | 0.0019 | 0.0035 | 0.54 |
| animacy[Animate]:task[C]:lat.[R]:sag.[post] | −0.0017 | 0.0035 | −0.49 |

Table S5: Visual study: Noun 1 P600 (later window) model.

Linear mixed model fit by REML

REML criterion at convergence: 632826

Scaled residuals:

| Min | 1Q | Median | 3Q | Max |
|---|---|---|---|---|
| -10 | -0.58 | 0 | 0.57 | 13.23 |

Random effects:

| Groups | Name | Std.Dev. |
|---|---|---|
| item | (Intercept) | 0.077876 |
| subj | (Intercept) | 0.114157 |
| Residual | | 0.868914 |

Number of obs: 247212, groups: item, 60; subj, 45.

Fixed effects:

| | Estimate | Std. Error | t value |
|---|---|---|---|
| (Intercept) | 0.004 | 0.02 | 0.2 |
| animacy[Animate] | −0.014 | 0.0018 | −8 |
| task[C] | −0.0027 | 0.017 | −0.16 |
| lat.[L] | −0.0031 | 0.0025 | −1.3 |
| lat.[R] | 0.0055 | 0.0025 | 2.2 |
| sag.[ant] | 0.03 | 0.0025 | 12 |
| sag.[post] | −0.031 | 0.0025 | −13 |
| scale(prestim) | −0.47 | 0.0018 | −2.6e+02 |
| animacy[Animate]:task[C] | −0.0023 | 0.0018 | −1.3 |
| animacy[Animate]:lat.[L] | 0.0017 | 0.0025 | 0.7 |
| animacy[Animate]:lat.[R] | 0.0014 | 0.0025 | 0.58 |
| task[C]:lat.[L] | −0.0025 | 0.0025 | −1 |
| task[C]:lat.[R] | 0.01 | 0.0025 | 4.1 |
| animacy[Animate]:sag.[ant] | −0.0025 | 0.0025 | −1 |
| animacy[Animate]:sag.[post] | 0.0044 | 0.0025 | 1.8 |
| task[C]:sag.[ant] | 0.01 | 0.0025 | 4.1 |
| task[C]:sag.[post] | −0.0047 | 0.0025 | −1.9 |
| lat.[L]:sag.[ant] | −0.025 | 0.0035 | −7.1 |
| lat.[R]:sag.[ant] | 0.014 | 0.0035 | 4 |
| lat.[L]:sag.[post] | 0.032 | 0.0035 | 9 |
| lat.[R]:sag.[post] | −0.02 | 0.0035 | −5.7 |
| animacy[Animate]:task[C]:lat.[L] | −0.0013 | 0.0025 | −0.53 |
| animacy[Animate]:task[C]:lat.[R] | 0.0035 | 0.0025 | 1.4 |
| animacy[Animate]:task[C]:sag.[ant] | 0.0018 | 0.0025 | 0.72 |
| animacy[Animate]:task[C]:sag.[post] | −0.002 | 0.0025 | −0.8 |
| animacy[Animate]:lat.[L]:sag.[ant] | −0.0026 | 0.0035 | −0.76 |
| animacy[Animate]:lat.[R]:sag.[ant] | −0.0038 | 0.0035 | −1.1 |
| animacy[Animate]:lat.[L]:sag.[post] | 0.0021 | 0.0035 | 0.59 |
| animacy[Animate]:lat.[R]:sag.[post] | 0.0012 | 0.0035 | 0.33 |
| task[C]:lat.[L]:sag.[ant] | 0.0067 | 0.0035 | 1.9 |
| task[C]:lat.[R]:sag.[ant] | −0.0084 | 0.0035 | −2.4 |
| task[C]:lat.[L]:sag.[post] | −0.0096 | 0.0035 | −2.7 |
| task[C]:lat.[R]:sag.[post] | 0.0087 | 0.0035 | 2.5 |
| animacy[Animate]:task[C]:lat.[L]:sag.[ant] | 0.0028 | 0.0035 | 0.82 |
| animacy[Animate]:task[C]:lat.[R]:sag.[ant] | 0.00012 | 0.0035 | 0.035 |
| animacy[Animate]:task[C]:lat.[L]:sag.[post] | −0.0017 | 0.0035 | −0.5 |
| animacy[Animate]:task[C]:lat.[R]:sag.[post] | −0.00036 | 0.0035 | −0.1 |

Table S6: Visual study: Verb N400.
Summary of residuals and random effects of model.

| Linear mixed model fit by REML | | | | |
|---|---|---|---|---|
| REML criterion at convergence: 1442764 | | | | |
| Scaled residuals: | | | | |
| Min | 1Q | Median | 3Q | Max |
| -7.99 | -0.59 | 0.02 | 0.6 | 9.43 |
| Random effects: | | | | |
| Groups | Name | Std.Dev. | | |
| item | (Intercept) | 0.44031 | | |
| subj | (Intercept) | 0.81661 | | |
| Residual | | 4.14379 | | |
| Number of obs: 253800, groups: item, 60; subj, 45. | | | | |

Table S7: Visual study: Verb N400. Summary of fixed effects of model.

Linear mixed model fit by REML

REML criterion at convergence: 1442764

Fixed effects:

| | Estimate | Std. Error | t value |
|---|---|---|---|
| (Intercept) | −1 | 0.13 | −7.4 |
| animacy[Grammatical] | 0.055 | 0.0082 | 6.7 |
| verbtype[ASV] | −0.048 | 0.0082 | −5.8 |
| task[C] | 0.046 | 0.12 | 0.38 |
| lat.[L] | 0.5 | 0.012 | 43 |
| lat.[R] | 0.016 | 0.012 | 1.4 |
| sag.[ant] | 0.33 | 0.012 | 28 |
| sag.[post] | −0.21 | 0.012 | −18 |
| scale(prestim) | −0.53 | 0.0084 | −63 |
| animacy[Grammatical]:verbtype[ASV] | 0.024 | 0.0082 | 3 |
| animacy[Grammatical]:task[C] | −0.0012 | 0.0082 | −0.14 |
| verbtype[ASV]:task[C] | 0.018 | 0.0082 | 2.1 |
| animacy[Grammatical]:lat.[L] | 0.0031 | 0.012 | 0.27 |
| animacy[Grammatical]:lat.[R] | −0.0071 | 0.012 | −0.61 |
| verbtype[ASV]:lat.[L] | 0.046 | 0.012 | 4 |
| verbtype[ASV]:lat.[R] | −0.031 | 0.012 | −2.7 |
| task[C]:lat.[L] | −0.077 | 0.012 | −6.6 |
| task[C]:lat.[R] | 0.027 | 0.012 | 2.3 |
| animacy[Grammatical]:sag.[ant] | −0.033 | 0.012 | −2.8 |
| animacy[Grammatical]:sag.[post] | 0.044 | 0.012 | 3.8 |
| verbtype[ASV]:sag.[ant] | −0.069 | 0.012 | −5.9 |
| verbtype[ASV]:sag.[post] | 0.057 | 0.012 | 4.9 |
| task[C]:sag.[ant] | −0.05 | 0.012 | −4.3 |
| task[C]:sag.[post] | 0.056 | 0.012 | 4.8 |
| lat.[L]:sag.[ant] | −0.075 | 0.016 | −4.5 |
| lat.[R]:sag.[ant] | −0.054 | 0.016 | −3.3 |
| lat.[L]:sag.[post] | 0.014 | 0.016 | 0.86 |
| lat.[R]:sag.[post] | 0.014 | 0.016 | 0.83 |
| animacy[Grammatical]:verbtype[ASV]:task[C] | −0.089 | 0.0082 | −11 |
| animacy[Grammatical]:verbtype[ASV]:lat.[L] | −0.021 | 0.012 | −1.8 |
| animacy[Grammatical]:verbtype[ASV]:lat.[R] | 0.0088 | 0.012 | 0.76 |
| animacy[Grammatical]:task[C]:lat.[L] | 0.062 | 0.012 | 5.3 |
| animacy[Grammatical]:task[C]:lat.[R] | −0.029 | 0.012 | −2.5 |
| verbtype[ASV]:task[C]:lat.[L] | −0.03 | 0.012 | −2.6 |
| verbtype[ASV]:task[C]:lat.[R] | 0.0044 | 0.012 | 0.38 |
| animacy[Grammatical]:verbtype[ASV]:sag.[ant] | −0.007 | 0.012 | −0.6 |
| animacy[Grammatical]:verbtype[ASV]:sag.[post] | 0.011 | 0.012 | 0.93 |
| animacy[Grammatical]:task[C]:sag.[ant] | 0.089 | 0.012 | 7.6 |
| animacy[Grammatical]:task[C]:sag.[post] | −0.09 | 0.012 | −7.8 |
| verbtype[ASV]:task[C]:sag.[ant] | −0.021 | 0.012 | −1.8 |
| verbtype[ASV]:task[C]:sag.[post] | 0.0031 | 0.012 | 0.27 |
| animacy[Grammatical]:lat.[L]:sag.[ant] | −0.0036 | 0.016 | −0.22 |
| animacy[Grammatical]:lat.[R]:sag.[ant] | 0.0064 | 0.016 | 0.39 |
| animacy[Grammatical]:lat.[L]:sag.[post] | 0.013 | 0.016 | 0.78 |
| animacy[Grammatical]:lat.[R]:sag.[post] | −0.011 | 0.016 | −0.66 |
| verbtype[ASV]:lat.[L]:sag.[ant] | 0.0038 | 0.016 | 0.23 |
| verbtype[ASV]:lat.[R]:sag.[ant] | −0.012 | 0.016 | −0.75 |
| verbtype[ASV]:lat.[L]:sag.[post] | −0.0084 | 0.016 | −0.51 |
| verbtype[ASV]:lat.[R]:sag.[post] | 0.013 | 0.016 | 0.76 |
| task[C]:lat.[L]:sag.[ant] | 0.015 | 0.016 | 0.89 |
| task[C]:lat.[R]:sag.[ant] | −0.0037 | 0.016 | −0.22 |
| task[C]:lat.[L]:sag.[post] | −0.015 | 0.016 | −0.92 |
| task[C]:lat.[R]:sag.[post] | 0.023 | 0.016 | 1.4 |
| animacy[Grammatical]:verbtype[ASV]:task[C]:lat.[L] | 0.057 | 0.012 | 4.9 |
| animacy[Grammatical]:verbtype[ASV]:task[C]:lat.[R] | −0.02 | 0.012 | −1.7 |
| animacy[Grammatical]:verbtype[ASV]:task[C]:sag.[ant] | −0.022 | 0.012 | −1.9 |
| animacy[Grammatical]:verbtype[ASV]:task[C]:sag.[post] | 0.026 | 0.012 | 2.3 |
| animacy[Grammatical]:verbtype[ASV]:lat.[L]:sag.[ant] | 0.0076 | 0.016 | 0.46 |
| animacy[Grammatical]:verbtype[ASV]:lat.[R]:sag.[ant] | −0.0014 | 0.016 | −0.085 |
| animacy[Grammatical]:verbtype[ASV]:lat.[L]:sag.[post] | −0.006 | 0.016 | −0.37 |
| animacy[Grammatical]:verbtype[ASV]:lat.[R]:sag.[post] | 0.0021 | 0.016 | 0.13 |
| animacy[Grammatical]:task[C]:lat.[L]:sag.[ant] | −0.00092 | 0.016 | −0.056 |
| animacy[Grammatical]:task[C]:lat.[R]:sag.[ant] | −0.00088 | 0.016 | −0.053 |
| animacy[Grammatical]:task[C]:lat.[L]:sag.[post] | −0.015 | 0.016 | −0.93 |
| animacy[Grammatical]:task[C]:lat.[R]:sag.[post] | 0.0087 | 0.016 | 0.53 |
| verbtype[ASV]:task[C]:lat.[L]:sag.[ant] | 0.01 | 0.016 | 0.61 |
| verbtype[ASV]:task[C]:lat.[R]:sag.[ant] | −0.0098 | 0.016 | −0.6 |
| verbtype[ASV]:task[C]:lat.[L]:sag.[post] | −0.0044 | 0.016 | −0.27 |
| verbtype[ASV]:task[C]:lat.[R]:sag.[post] | 0.011 | 0.016 | 0.69 |
| animacy[Grammatical]:verbtype[ASV]:task[C]:lat.[L]:sag.[ant] | 0.01 | 0.016 | 0.61 |
| animacy[Grammatical]:verbtype[ASV]:task[C]:lat.[R]:sag.[ant] | −0.0026 | 0.016 | −0.16 |
| animacy[Grammatical]:verbtype[ASV]:task[C]:lat.[L]:sag.[post] | −0.023 | 0.016 | −1.4 |
| animacy[Grammatical]:verbtype[ASV]:task[C]:lat.[R]:sag.[post] | 0.0087 | 0.016 | 0.53 |

Table S8: Visual study: Verb P600.
Summary of residuals and random effects of model.

Linear mixed model fit by REML

REML criterion at convergence: 1426043

Scaled residuals:

| Min | 1Q | Median | 3Q | Max |
|---|---|---|---|---|
| -6.9 | -0.59 | -0.01 | 0.59 | 9.64 |

Random effects:

| Groups | Name | Std.Dev. |
|---|---|---|
| item | (Intercept) | 0.36410 |
| subj | (Intercept) | 0.63175 |
| Residual | | 4.00977 |

Number of obs: 253800, groups: item, 60; subj, 45.

Table S9: Visual study: Verb P600.Summary of fixed effects of model.

Linear mixed model fit by REML

REML criterion at convergence: 1426043

Fixed effects:

| | Estimate | Std. Error | t value |
|---|---|---|---|
| (Intercept) | 0.69 | 0.11 | 6.5 |
| animacy[Grammatical] | −0.046 | 0.008 | −5.7 |
| verbtype[ASV] | 0.06 | 0.008 | 7.5 |
| task[C] | −0.24 | 0.095 | −2.6 |
| lat.[L] | −0.2 | 0.011 | −18 |
| lat.[R] | 0.029 | 0.011 | 2.6 |
| sag.[ant] | −0.25 | 0.011 | −22 |
| sag.[post] | 0.18 | 0.011 | 16 |
| scale(prestim) | −2.2 | 0.0081 | −2.7e+02 |
| animacy[Grammatical]:verbtype[ASV] | −0.06 | 0.008 | −7.5 |
| animacy[Grammatical]:task[C] | −0.041 | 0.008 | −5.2 |
| verbtype[ASV]:task[C] | 0.051 | 0.008 | 6.4 |
| animacy[Grammatical]:lat.[L] | −0.019 | 0.011 | −1.7 |
| animacy[Grammatical]:lat.[R] | 0.043 | 0.011 | 3.8 |
| verbtype[ASV]:lat.[L] | −0.0012 | 0.011 | −0.1 |
| verbtype[ASV]:lat.[R] | −0.02 | 0.011 | −1.7 |
| task[C]:lat.[L] | 0.09 | 0.011 | 8 |
| task[C]:lat.[R] | 0.045 | 0.011 | 4 |
| animacy[Grammatical]:sag.[ant] | 0.02 | 0.011 | 1.8 |
| animacy[Grammatical]:sag.[post] | −0.0055 | 0.011 | −0.49 |
| verbtype[ASV]:sag.[ant] | 0.027 | 0.011 | 2.4 |
| verbtype[ASV]:sag.[post] | −0.023 | 0.011 | −2 |
| task[C]:sag.[ant] | 0.017 | 0.011 | 1.5 |
| task[C]:sag.[post] | 0.027 | 0.011 | 2.4 |
| lat.[L]:sag.[ant] | 0.012 | 0.016 | 0.74 |
| lat.[R]:sag.[ant] | 0.016 | 0.016 | 1 |
| lat.[L]:sag.[post] | 0.043 | 0.016 | 2.7 |
| lat.[R]:sag.[post] | −0.0097 | 0.016 | −0.61 |
| animacy[Grammatical]:verbtype[ASV]:task[C] | −0.038 | 0.008 | −4.7 |
| animacy[Grammatical]:verbtype[ASV]:lat.[L] | 0.0055 | 0.011 | 0.48 |
| animacy[Grammatical]:verbtype[ASV]:lat.[R] | 0.021 | 0.011 | 1.8 |
| animacy[Grammatical]:task[C]:lat.[L] | −0.044 | 0.011 | −3.9 |
| animacy[Grammatical]:task[C]:lat.[R] | 0.025 | 0.011 | 2.2 |
| verbtype[ASV]:task[C]:lat.[L] | −0.006 | 0.011 | −0.53 |
| verbtype[ASV]:task[C]:lat.[R] | −0.0043 | 0.011 | −0.38 |
| animacy[Grammatical]:verbtype[ASV]:sag.[ant] | −0.061 | 0.011 | −5.4 |
| animacy[Grammatical]:verbtype[ASV]:sag.[post] | 0.054 | 0.011 | 4.8 |
| animacy[Grammatical]:task[C]:sag.[ant] | −0.058 | 0.011 | −5.2 |
| animacy[Grammatical]:task[C]:sag.[post] | 0.048 | 0.011 | 4.2 |
| verbtype[ASV]:task[C]:sag.[ant] | 0.021 | 0.011 | 1.8 |
| verbtype[ASV]:task[C]:sag.[post] | −0.018 | 0.011 | −1.6 |
| animacy[Grammatical]:lat.[L]:sag.[ant] | 0.00043 | 0.016 | 0.027 |
| animacy[Grammatical]:lat.[R]:sag.[ant] | 0.0092 | 0.016 | 0.58 |
| animacy[Grammatical]:lat.[L]:sag.[post] | −0.004 | 0.016 | −0.25 |
| animacy[Grammatical]:lat.[R]:sag.[post] | −0.012 | 0.016 | −0.73 |
| verbtype[ASV]:lat.[L]:sag.[ant] | −0.0025 | 0.016 | −0.16 |
| verbtype[ASV]:lat.[R]:sag.[ant] | 0.003 | 0.016 | 0.19 |
| verbtype[ASV]:lat.[L]:sag.[post] | −0.002 | 0.016 | −0.13 |
| verbtype[ASV]:lat.[R]:sag.[post] | 0.00062 | 0.016 | 0.039 |
| task[C]:lat.[L]:sag.[ant] | 0.052 | 0.016 | 3.3 |
| task[C]:lat.[R]:sag.[ant] | −0.085 | 0.016 | −5.4 |
| task[C]:lat.[L]:sag.[post] | −0.088 | 0.016 | −5.5 |
| task[C]:lat.[R]:sag.[post] | 0.097 | 0.016 | 6.1 |
| animacy[Grammatical]:verbtype[ASV]:task[C]:lat.[L] | −0.024 | 0.011 | −2.1 |
| animacy[Grammatical]:verbtype[ASV]:task[C]:lat.[R] | 0.04 | 0.011 | 3.6 |
| animacy[Grammatical]:verbtype[ASV]:task[C]:sag.[ant] | −0.0062 | 0.011 | −0.55 |
| animacy[Grammatical]:verbtype[ASV]:task[C]:sag.[post] | 0.0052 | 0.011 | 0.46 |
| animacy[Grammatical]:verbtype[ASV]:lat.[L]:sag.[ant] | −0.0002 | 0.016 | −0.013 |
| animacy[Grammatical]:verbtype[ASV]:lat.[R]:sag.[ant] | 0.0042 | 0.016 | 0.26 |
| animacy[Grammatical]:verbtype[ASV]:lat.[L]:sag.[post] | 0.0012 | 0.016 | 0.076 |
| animacy[Grammatical]:verbtype[ASV]:lat.[R]:sag.[post] | −0.0038 | 0.016 | −0.24 |
| animacy[Grammatical]:task[C]:lat.[L]:sag.[ant] | −0.023 | 0.016 | −1.5 |
| animacy[Grammatical]:task[C]:lat.[R]:sag.[ant] | 0.0079 | 0.016 | 0.49 |
| animacy[Grammatical]:task[C]:lat.[L]:sag.[post] | 0.034 | 0.016 | 2.1 |
| animacy[Grammatical]:task[C]:lat.[R]:sag.[post] | −0.014 | 0.016 | −0.86 |
| verbtype[ASV]:task[C]:lat.[L]:sag.[ant] | −0.0015 | 0.016 | −0.097 |
| verbtype[ASV]:task[C]:lat.[R]:sag.[ant] | 0.0021 | 0.016 | 0.13 |
| verbtype[ASV]:task[C]:lat.[L]:sag.[post] | 0.00052 | 0.016 | 0.033 |
| verbtype[ASV]:task[C]:lat.[R]:sag.[post] | 0.0015 | 0.016 | 0.092 |
| animacy[Grammatical]:verbtype[ASV]:task[C]:lat.[L]:sag.[ant] | −0.0052 | 0.016 | −0.32 |
| animacy[Grammatical]:verbtype[ASV]:task[C]:lat.[R]:sag.[ant] | 0.0081 | 0.016 | 0.51 |
| animacy[Grammatical]:verbtype[ASV]:task[C]:lat.[L]:sag.[post] | 0.0045 | 0.016 | 0.28 |
| animacy[Grammatical]:verbtype[ASV]:task[C]:lat.[R]:sag.[post] | −0.013 | 0.016 | −0.79 |

Table S10: Auditory study: Judgement behavioural model.

Generalized linear mixed model fit by maximum likelihood (Laplace Approximation)
Family: binomial Link:logit

| | AIC | BIC | logLik | deviance | | |
|---|---|---|---|---|---|---|
| | 408746 | 408803 | -204368 | 408736 | | |
| Scaled residuals: | | | | | | |
| | Min | 1Q | Median | 3Q | Max | |
| | -5.49 | 0.25 | 0.31 | 0.38 | 0.6 | |
| Random effects: | | | | | | |
| | Groups | Name | Std.Dev. | | | |
| | subj | (Intercept) | 0.36248 | | | |
| Number of obs: 637470, groups: subj, 23. | | | | | | |
| Fixed effects: | | | | | | |
| | | Estimate | Std. Error | z value | Pr(>\|z\|) | |
| (Intercept) | | 2.3 | 0.071 | 32 | 7.4e−223 | *** |
| animacy[Grammatical] | | −0.016 | 0.0043 | −3.7 | 0.00019 | *** |
| verbtype[ASV] | | 0.32 | 0.0043 | 74 | 0 | *** |
| animacy[Grammatical]:verbtype[ASV] | | −0.0033 | 0.0043 | −0.76 | 0.45 | |

Table S11: Auditory study: Comprehension behavioural model.

Generalized linear mixed model fit by maximum likelihood (Laplace Approximation)
Family: binomial Link:logit

| AIC | BIC | logLik | deviance |
|---|---|---|---|
| 356876 | 356974 | -178429 | 356858 |

Scaled residuals:

| Min | 1Q | Median | 3Q | Max |
|---|---|---|---|---|
| -9.36 | 0.12 | 0.36 | 0.54 | 1.41 |

Random effects:

| Groups | Name | Std.Dev. |
|---|---|---|
| subj | (Intercept) | 1.0621 |

Number of obs: 403650, groups: subj, 24.

Fixed effects:

| | Estimate | Std. Error | z value | Pr(>\|z\|) | |
|---|---|---|---|---|---|
| (Intercept) | 1.6 | 0.13 | 13 | 5.7e−38 | *[illegible] |
| animacy[Grammatical] | 0.42 | 0.0043 | 98 | 0 | *[illegible] |
| verbtype[ASV] | 0.18 | 0.0043 | 41 | 0 | *[illegible] |
| qtype[Actor Probe] | 0.11 | 0.0043 | 25 | 4.4e−142 | *[illegible] |
| animacy[Grammatical]:verbtype[ASV] | 0.11 | 0.0043 | 25 | 2.3e−134 | *[illegible] |
| animacy[Grammatical]:qtype[Actor Probe] | −0.17 | 0.0043 | −38 | 0 | *[illegible] |
| verbtype[ASV]:qtype[Actor Probe] | −0.15 | 0.0043 | −36 | 1.5e−279 | *[illegible] |
| animacy[Grammatical]:verbtype[ASV]:qtype[Actor Probe] | −0.0065 | 0.0043 | −1.5 | 0.13 | |

Table S12: Auditory study: Noun 1 N400 model.

Linear mixed model fit by REML

REML criterion at convergence: 669486

Scaled residuals:

| Min | 1Q | Median | 3Q | Max |
|---|---|---|---|---|
| -11.87 | -0.58 | 0.02 | 0.59 | 9.13 |

Random effects:

| Groups | Name | Std.Dev. |
|---|---|---|
| item | (Intercept) | 0.09304 |
| subj | (Intercept) | 0.13721 |
| Residual | | 0.98339 |

Number of obs: 238464, groups: item, 60; subj, 47.

Fixed effects:

| | Estimate | Std. Error | t value |
|---|---|---|---|
| (Intercept) | −0.014 | 0.023 | −0.59 |
| animacy[Animate] | 0.018 | 0.002 | 8.7 |
| task[C] | −0.023 | 0.02 | −1.1 |
| lat.[L] | 0.022 | 0.0029 | 7.8 |
| lat.[R] | 0.046 | 0.0028 | 16 |
| sag.[ant] | 0.086 | 0.0028 | 30 |
| sag.[post] | −0.079 | 0.0028 | −28 |
| scale(prestim) | 0.012 | 0.0021 | 6 |
| animacy[Animate]:task[C] | −0.011 | 0.002 | −5.6 |
| animacy[Animate]:lat.[L] | −0.012 | 0.0028 | −4.1 |
| animacy[Animate]:lat.[R] | −0.0033 | 0.0028 | −1.1 |
| task[C]:lat.[L] | −0.0076 | 0.0028 | −2.7 |
| task[C]:lat.[R] | 0.016 | 0.0028 | 5.7 |
| animacy[Animate]:sag.[ant] | −0.028 | 0.0028 | −9.9 |
| animacy[Animate]:sag.[post] | 0.024 | 0.0028 | 8.5 |
| task[C]:sag.[ant] | −0.013 | 0.0028 | −4.6 |
| task[C]:sag.[post] | 0.018 | 0.0028 | 6.3 |
| lat.[L]:sag.[ant] | 0.0045 | 0.004 | 1.1 |
| lat.[R]:sag.[ant] | 0.0043 | 0.004 | 1.1 |
| lat.[L]:sag.[post] | −0.012 | 0.004 | −3 |
| lat.[R]:sag.[post] | −0.014 | 0.004 | −3.4 |
| animacy[Animate]:task[C]:lat.[L] | −0.0051 | 0.0028 | −1.8 |
| animacy[Animate]:task[C]:lat.[R] | 0.0047 | 0.0028 | 1.7 |
| animacy[Animate]:task[C]:sag.[ant] | −0.0043 | 0.0028 | −1.5 |
| animacy[Animate]:task[C]:sag.[post] | 0.0069 | 0.0028 | 2.4 |
| animacy[Animate]:lat.[L]:sag.[ant] | −0.0069 | 0.004 | −1.7 |
| animacy[Animate]:lat.[R]:sag.[ant] | 0.0053 | 0.004 | 1.3 |
| animacy[Animate]:lat.[L]:sag.[post] | 0.011 | 0.004 | 2.6 |
| animacy[Animate]:lat.[R]:sag.[post] | −0.006 | 0.004 | −1.5 |
| task[C]:lat.[L]:sag.[ant] | −0.0029 | 0.004 | −0.73 |
| task[C]:lat.[R]:sag.[ant] | 0.0075 | 0.004 | 1.9 |
| task[C]:lat.[L]:sag.[post] | 0.0042 | 0.004 | 1 |
| task[C]:lat.[R]:sag.[post] | −0.01 | 0.004 | −2.5 |
| animacy[Animate]:task[C]:lat.[L]:sag.[ant] | 0.0032 | 0.004 | 0.8 |
| animacy[Animate]:task[C]:lat.[R]:sag.[ant] | −0.0039 | 0.004 | −0.96 |
| animacy[Animate]:task[C]:lat.[L]:sag.[post] | −0.0031 | 0.004 | −0.77 |
| animacy[Animate]:task[C]:lat.[R]:sag.[post] | 0.0045 | 0.004 | 1.1 |

Table S13: Auditory study: Noun 1 P600 model.

| Linear mixed model fit by REML | | | | | |
|---|---|---|---|---|---|
| REML criterion at convergence: 641525 | | | | | |
| Scaled residuals: | | | | | |
| | Min | 1Q | Median | 3Q | Max |
| | -13.83 | -0.57 | 0 | 0.57 | 11.55 |
| Random effects: | | | | | |
| | Groups | Name | Std.Dev. | | |
| | item | (Intercept) | 0.084541 | | |
| | subj | (Intercept) | 0.131605 | | |
| | Residual | | 0.927394 | | |
| Number of obs: 238464, groups: item, 60; subj, 47. | | | | | |
| Fixed effects: | | | | | |
| | | Estimate | Std. Error | t value | |
| | (Intercept) | 0.013 | 0.022 | 0.58 | |
| | animacy[Animate] | 0.0053 | 0.0019 | 2.8 | |
| | task[C] | 0.0099 | 0.019 | 0.51 | |
| | lat.[L] | −0.018 | 0.0027 | −6.7 | |
| | lat.[R] | −0.0042 | 0.0027 | −1.5 | |
| | sag.[ant] | −0.018 | 0.0027 | −6.6 | |
| | sag.[post] | 0.034 | 0.0027 | 13 | |
| | scale(prestim) | −0.35 | 0.002 | −1.8e+02 | |
| | animacy[Animate]:task[C] | −0.00016 | 0.0019 | −0.084 | |
| | animacy[Animate]:lat.[L] | −0.012 | 0.0027 | −4.3 | |
| | animacy[Animate]:lat.[R] | 0.0027 | 0.0027 | 1 | |
| | task[C]:lat.[L] | 0.0008 | 0.0027 | 0.3 | |
| | task[C]:lat.[R] | −0.0054 | 0.0027 | −2 | |
| | animacy[Animate]:sag.[ant] | −0.019 | 0.0027 | −7.2 | |
| | animacy[Animate]:sag.[post] | 0.018 | 0.0027 | 6.7 | |
| | task[C]:sag.[ant] | −0.01 | 0.0027 | −3.8 | |
| | task[C]:sag.[post] | 0.0033 | 0.0027 | 1.2 | |
| | lat.[L]:sag.[ant] | 0.0066 | 0.0038 | 1.7 | |
| | lat.[R]:sag.[ant] | −0.001 | 0.0038 | −0.27 | |
| | lat.[L]:sag.[post] | 0.0022 | 0.0038 | 0.57 | |
| | lat.[R]:sag.[post] | 0.0058 | 0.0038 | 1.5 | |
| | animacy[Animate]:task[C]:lat.[L] | 0.014 | 0.0027 | 5.3 | |
| | animacy[Animate]:task[C]:lat.[R] | −0.014 | 0.0027 | −5.2 | |
| | animacy[Animate]:task[C]:sag.[ant] | −0.0015 | 0.0027 | −0.55 | |
| | animacy[Animate]:task[C]:sag.[post] | −0.0016 | 0.0027 | −0.59 | |
| | animacy[Animate]:lat.[L]:sag.[ant] | −0.0024 | 0.0038 | −0.63 | |
| | animacy[Animate]:lat.[R]:sag.[ant] | 0.001 | 0.0038 | 0.27 | |
| | animacy[Animate]:lat.[L]:sag.[post] | 0.0019 | 0.0038 | 0.51 | |
| | animacy[Animate]:lat.[R]:sag.[post] | 0.00092 | 0.0038 | 0.24 | |
| | task[C]:lat.[L]:sag.[ant] | 0.0012 | 0.0038 | 0.31 | |
| | task[C]:lat.[R]:sag.[ant] | −0.0082 | 0.0038 | −2.2 | |
| | task[C]:lat.[L]:sag.[post] | −0.0038 | 0.0038 | −1 | |
| | task[C]:lat.[R]:sag.[post] | 0.0079 | 0.0038 | 2.1 | |
| | animacy[Animate]:task[C]:lat.[L]:sag.[ant] | 0.002 | 0.0038 | 0.52 | |
| | animacy[Animate]:task[C]:lat.[R]:sag.[ant] | −0.0019 | 0.0038 | −0.51 | |
| | animacy[Animate]:task[C]:lat.[L]:sag.[post] | −0.0026 | 0.0038 | −0.69 | |
| | animacy[Animate]:task[C]:lat.[R]:sag.[post] | 0.0031 | 0.0038 | 0.81 | |

Table S14: Auditory study: Noun 1 P600 (later window) model.

Linear mixed model fit by REML

REML criterion at convergence: 621096

Scaled residuals:

| Min | 1Q | Median | 3Q | Max |
|---|---|---|---|---|
| -9.68 | -0.58 | 0 | 0.58 | 10.5 |

Random effects:

| Groups | Name | Std.Dev. |
|---|---|---|
| item | (Intercept) | 0.093683 |
| subj | (Intercept) | 0.123056 |
| Residual | | 0.888474 |

Number of obs: 238464, groups: item, 60; subj, 47.

Fixed effects:

| | Estimate | Std. Error | t value |
|---|---|---|---|
| (Intercept) | 0.01 | 0.022 | 0.48 |
| animacy[Animate] | 0.01 | 0.0018 | 5.7 |
| task[C] | 0.0054 | 0.018 | 0.3 |
| lat.[L] | −0.0014 | 0.0026 | −0.55 |
| lat.[R] | −0.021 | 0.0026 | −8.1 |
| sag.[ant] | −0.022 | 0.0026 | −8.5 |
| sag.[post] | 0.038 | 0.0026 | 15 |
| scale(prestim) | −0.44 | 0.0019 | −2.3e+02 |
| animacy[Animate]:task[C] | 0.0097 | 0.0018 | 5.3 |
| animacy[Animate]:lat.[L] | −0.0079 | 0.0026 | −3.1 |
| animacy[Animate]:lat.[R] | 0.012 | 0.0026 | 4.6 |
| task[C]:lat.[L] | 0.00025 | 0.0026 | 0.096 |
| task[C]:lat.[R] | −0.0081 | 0.0026 | −3.2 |
| animacy[Animate]:sag.[ant] | 0.021 | 0.0026 | 8.1 |
| animacy[Animate]:sag.[post] | −0.019 | 0.0026 | −7.5 |
| task[C]:sag.[ant] | −0.0094 | 0.0026 | −3.7 |
| task[C]:sag.[post] | 0.0028 | 0.0026 | 1.1 |
| lat.[L]:sag.[ant] | 0.014 | 0.0036 | 3.9 |
| lat.[R]:sag.[ant] | −0.01 | 0.0036 | −2.8 |
| lat.[L]:sag.[post] | −0.0059 | 0.0036 | −1.6 |
| lat.[R]:sag.[post] | 0.015 | 0.0036 | 4.2 |
| animacy[Animate]:task[C]:lat.[L] | 0.0067 | 0.0026 | 2.6 |
| animacy[Animate]:task[C]:lat.[R] | −0.0075 | 0.0026 | −2.9 |
| animacy[Animate]:task[C]:sag.[ant] | 0.0099 | 0.0026 | 3.8 |
| animacy[Animate]:task[C]:sag.[post] | −0.012 | 0.0026 | −4.5 |
| animacy[Animate]:lat.[L]:sag.[ant] | −0.0017 | 0.0036 | −0.46 |
| animacy[Animate]:lat.[R]:sag.[ant] | 0.0053 | 0.0036 | 1.5 |
| animacy[Animate]:lat.[L]:sag.[post] | 0.0025 | 0.0036 | 0.68 |
| animacy[Animate]:lat.[R]:sag.[post] | −0.006 | 0.0036 | −1.6 |
| task[C]:lat.[L]:sag.[ant] | −0.0028 | 0.0036 | −0.77 |
| task[C]:lat.[R]:sag.[ant] | −0.0065 | 0.0036 | −1.8 |
| task[C]:lat.[L]:sag.[post] | 0.002 | 0.0036 | 0.56 |
| task[C]:lat.[R]:sag.[post] | 0.0054 | 0.0036 | 1.5 |
| animacy[Animate]:task[C]:lat.[L]:sag.[ant] | −0.00025 | 0.0036 | −0.068 |
| animacy[Animate]:task[C]:lat.[R]:sag.[ant] | −0.00053 | 0.0036 | −0.15 |
| animacy[Animate]:task[C]:lat.[L]:sag.[post] | −0.0017 | 0.0036 | −0.47 |
| animacy[Animate]:task[C]:lat.[R]:sag.[post] | 0.0011 | 0.0036 | 0.3 |

Table S15: Auditory study: Verb N400.
Summary of residuals and random effects of model.

| Linear mixed model fit by REML | | | | |
|---|---|---|---|---|
| REML criterion at convergence: 1482684 | | | | |
| | | | Scaled residuals | |
| Min | 1Q | Median | 3Q | Max |
| -12.47 | -0.58 | 0 | 0.59 | 9.87 |
| Random effects: | | | | |
| Groups | Name | Std.Dev. | | |
| item | (Intercept) | 0.55729 | | |
| subj | (Intercept) | 0.52248 | | |
| Residual | | 4.70972 | | |
| Number of obs: 249588, groups: item, 60; subj, 47. | | | | |

Table S16: Auditory study: Verb N400.Summary of fixed effects of model.

Linear mixed model fit by REML
Fixed effects:

| | Estimate | Std. Error | t value |
|---|---|---|---|
| (Intercept) | −0.059 | 0.11 | −0.56 |
| animacy[Grammatical] | 0.047 | 0.0094 | 5 |
| verbtype[ASV] | −0.02 | 0.0095 | −2.1 |
| task[C] | −0.0082 | 0.077 | −0.11 |
| lat.[L] | 0.037 | 0.013 | 2.8 |
| lat.[R] | 0.11 | 0.013 | 8.1 |
| sag.[ant] | 0.47 | 0.013 | 35 |
| sag.[post] | −0.38 | 0.013 | −28 |
| scale(prestim) | −0.062 | 0.0097 | −6.4 |
| animacy[Grammatical]:verbtype[ASV] | 0.039 | 0.0095 | 4.1 |
| animacy[Grammatical]:task[C] | −0.039 | 0.0094 | −4.2 |
| verbtype[ASV]:task[C] | 0.098 | 0.0094 | 10 |
| animacy[Grammatical]:lat.[L] | −0.062 | 0.013 | −4.7 |
| animacy[Grammatical]:lat.[R] | 0.068 | 0.013 | 5.1 |
| verbtype[ASV]:lat.[L] | −0.023 | 0.013 | −1.7 |
| verbtype[ASV]:lat.[R] | 0.01 | 0.013 | 0.76 |
| task[C]:lat.[L] | −0.053 | 0.013 | −3.9 |
| task[C]:lat.[R] | 0.057 | 0.013 | 4.3 |
| animacy[Grammatical]:sag.[ant] | −0.061 | 0.013 | −4.6 |
| animacy[Grammatical]:sag.[post] | 0.054 | 0.013 | 4 |
| verbtype[ASV]:sag.[ant] | −0.059 | 0.013 | −4.4 |
| verbtype[ASV]:sag.[post] | 0.072 | 0.013 | 5.4 |
| task[C]:sag.[ant] | −0.094 | 0.013 | −7 |
| task[C]:sag.[post] | 0.085 | 0.013 | 6.3 |
| lat.[L]:sag.[ant] | 0.14 | 0.019 | 7.3 |
| lat.[R]:sag.[ant] | −0.026 | 0.019 | −1.4 |
| lat.[L]:sag.[post] | −0.11 | 0.019 | −5.7 |
| lat.[R]:sag.[post] | 0.016 | 0.019 | 0.82 |
| animacy[Grammatical]:verbtype[ASV]:task[C] | 0.031 | 0.0095 | 3.3 |
| animacy[Grammatical]:verbtype[ASV]:lat.[L] | 0.026 | 0.013 | 2 |
| animacy[Grammatical]:verbtype[ASV]:lat.[R] | −0.023 | 0.013 | −1.7 |
| animacy[Grammatical]:task[C]:lat.[L] | 0.069 | 0.013 | 5.2 |
| animacy[Grammatical]:task[C]:lat.[R] | −0.047 | 0.013 | −3.5 |
| verbtype[ASV]:task[C]:lat.[L] | 0.0077 | 0.013 | 0.58 |
| verbtype[ASV]:task[C]:lat.[R] | −0.047 | 0.013 | −3.5 |
| animacy[Grammatical]:verbtype[ASV]:sag.[ant] | −0.019 | 0.013 | −1.4 |
| animacy[Grammatical]:verbtype[ASV]:sag.[post] | 0.013 | 0.013 | 1 |
| animacy[Grammatical]:task[C]:sag.[ant] | 0.062 | 0.013 | 4.6 |
| animacy[Grammatical]:task[C]:sag.[post] | −0.046 | 0.013 | −3.5 |
| verbtype[ASV]:task[C]:sag.[ant] | 0.025 | 0.013 | 1.9 |
| verbtype[ASV]:task[C]:sag.[post] | −0.04 | 0.013 | −3 |
| animacy[Grammatical]:lat.[L]:sag.[ant] | −0.034 | 0.019 | −1.8 |
| animacy[Grammatical]:lat.[R]:sag.[ant] | 0.042 | 0.019 | 2.2 |
| animacy[Grammatical]:lat.[L]:sag.[post] | 0.049 | 0.019 | 2.6 |
| animacy[Grammatical]:lat.[R]:sag.[post] | −0.059 | 0.019 | −3.2 |
| verbtype[ASV]:lat.[L]:sag.[ant] | 0.00035 | 0.019 | 0.019 |
| verbtype[ASV]:lat.[R]:sag.[ant] | 0.013 | 0.019 | 0.69 |
| verbtype[ASV]:lat.[L]:sag.[post] | −0.0015 | 0.019 | −0.082 |
| verbtype[ASV]:lat.[R]:sag.[post] | −0.0066 | 0.019 | −0.35 |
| task[C]:lat.[L]:sag.[ant] | −0.05 | 0.019 | −2.6 |
| task[C]:lat.[R]:sag.[ant] | −0.0098 | 0.019 | −0.52 |
| task[C]:lat.[L]:sag.[post] | 0.036 | 0.019 | 1.9 |
| task[C]:lat.[R]:sag.[post] | −0.011 | 0.019 | −0.56 |
| animacy[Grammatical]:verbtype[ASV]:task[C]:lat.[L] | −0.023 | 0.013 | −1.8 |
| animacy[Grammatical]:verbtype[ASV]:task[C]:lat.[R] | 0.01 | 0.013 | 0.77 |
| animacy[Grammatical]:verbtype[ASV]:task[C]:sag.[ant] | 0.022 | 0.013 | 1.6 |
| animacy[Grammatical]:verbtype[ASV]:task[C]:sag.[post] | −0.018 | 0.013 | −1.3 |
| animacy[Grammatical]:verbtype[ASV]:lat.[L]:sag.[ant] | 0.0036 | 0.019 | 0.19 |
| animacy[Grammatical]:verbtype[ASV]:lat.[R]:sag.[ant] | 0.0053 | 0.019 | 0.28 |
| animacy[Grammatical]:verbtype[ASV]:lat.[L]:sag.[post] | −0.002 | 0.019 | −0.11 |
| animacy[Grammatical]:verbtype[ASV]:lat.[R]:sag.[post] | −0.019 | 0.019 | −1 |
| animacy[Grammatical]:task[C]:lat.[L]:sag.[ant] | 0.03 | 0.019 | 1.6 |
| animacy[Grammatical]:task[C]:lat.[R]:sag.[ant] | −0.041 | 0.019 | −2.2 |
| animacy[Grammatical]:task[C]:lat.[L]:sag.[post] | −0.043 | 0.019 | −2.3 |
| animacy[Grammatical]:task[C]:lat.[R]:sag.[post] | 0.041 | 0.019 | 2.2 |
| verbtype[ASV]:task[C]:lat.[L]:sag.[ant] | −0.033 | 0.019 | −1.7 |
| verbtype[ASV]:task[C]:lat.[R]:sag.[ant] | 0.0064 | 0.019 | 0.34 |
| verbtype[ASV]:task[C]:lat.[L]:sag.[post] | 0.04 | 0.019 | 2.1 |
| verbtype[ASV]:task[C]:lat.[R]:sag.[post] | −0.003 | 0.019 | −0.16 |
| animacy[Grammatical]:verbtype[ASV]:task[C]:lat.[L]:sag.[ant] | 0.02 | 0.019 | 1.1 |
| animacy[Grammatical]:verbtype[ASV]:task[C]:lat.[R]:sag.[ant] | −0.021 | 0.019 | −1.1 |
| animacy[Grammatical]:verbtype[ASV]:task[C]:lat.[L]:sag.[post] | −0.018 | 0.019 | −0.95 |
| animacy[Grammatical]:verbtype[ASV]:task[C]:lat.[R]:sag.[post] | 0.026 | 0.019 | 1.4 |

Table S17: Auditory study: Verb P600.
Summary of residuals and random effects of model.

| Linear mixed model fit by REML | | | | |
|---|---|---|---|---|
| REML criterion at convergence: 1457113 | | | | |
| Scaled residuals: | | | | |
| Min | 1Q | Median | 3Q | Max |
| -9.85 | -0.58 | 0 | 0.59 | 10.26 |
| Random effects: | | | | |
| Groups | Name | Std.Dev. | | |
| item | (Intercept) | 0.37870 | | |
| subj | (Intercept) | 0.38294 | | |
| Residual | | 4.47502 | | |
| Number of obs: 249588, groups: item, 60; subj, 47. | | | | |

Table S18: Auditory study: Verb P600.Summary of fixed effects of model.

Linear mixed model fit by REML

REML criterion at convergence: 1457113

Fixed effects:

| | Estimate | Std. Error | t value |
|---|---|---|---|
| (Intercept) | −0.28 | 0.075 | −3.7 |
| animacy[Grammatical] | −0.012 | 0.009 | −1.3 |
| verbtype[ASV] | 0.013 | 0.009 | 1.5 |
| task[C] | 0.056 | 0.057 | 0.98 |
| lat.[L] | 0.072 | 0.013 | 5.7 |
| lat.[R] | −0.051 | 0.013 | −4 |
| sag.[ant] | −0.15 | 0.013 | −12 |
| sag.[post] | 0.18 | 0.013 | 14 |
| scale(prestim) | −1.7 | 0.0092 | −1.8e+02 |
| animacy[Grammatical]:verbtype[ASV] | −0.054 | 0.009 | −6.1 |
| animacy[Grammatical]:task[C] | 0.035 | 0.009 | 3.9 |
| verbtype[ASV]:task[C] | −0.047 | 0.009 | −5.3 |
| animacy[Grammatical]:lat.[L] | 0.0054 | 0.013 | 0.42 |
| animacy[Grammatical]:lat.[R] | −0.022 | 0.013 | −1.8 |
| verbtype[ASV]:lat.[L] | 0.015 | 0.013 | 1.2 |
| verbtype[ASV]:lat.[R] | −0.012 | 0.013 | −0.95 |
| task[C]:lat.[L] | −0.038 | 0.013 | −3 |
| task[C]:lat.[R] | 0.0084 | 0.013 | 0.66 |
| animacy[Grammatical]:sag.[ant] | −0.0015 | 0.013 | −0.12 |
| animacy[Grammatical]:sag.[post] | 0.0074 | 0.013 | 0.58 |
| verbtype[ASV]:sag.[ant] | −0.034 | 0.013 | −2.7 |
| verbtype[ASV]:sag.[post] | 0.017 | 0.013 | 1.3 |
| task[C]:sag.[ant] | 0.073 | 0.013 | 5.8 |
| task[C]:sag.[post] | −0.076 | 0.013 | −6 |
| lat.[L]:sag.[ant] | −0.015 | 0.018 | −0.86 |
| lat.[R]:sag.[ant] | 0.0084 | 0.018 | 0.47 |
| lat.[L]:sag.[post] | 0.011 | 0.018 | 0.63 |
| lat.[R]:sag.[post] | −0.0066 | 0.018 | −0.37 |
| animacy[Grammatical]:verbtype[ASV]:task[C] | −0.041 | 0.009 | −4.6 |
| animacy[Grammatical]:verbtype[ASV]:lat.[L] | 0.0012 | 0.013 | 0.095 |
| animacy[Grammatical]:verbtype[ASV]:lat.[R] | 0.00023 | 0.013 | 0.018 |
| animacy[Grammatical]:task[C]:lat.[L] | −0.023 | 0.013 | −1.8 |
| animacy[Grammatical]:task[C]:lat.[R] | 0.011 | 0.013 | 0.86 |
| verbtype[ASV]:task[C]:lat.[L] | −0.042 | 0.013 | −3.3 |
| verbtype[ASV]:task[C]:lat.[R] | 0.051 | 0.013 | 4 |
| animacy[Grammatical]:verbtype[ASV]:sag.[ant] | −0.0028 | 0.013 | −0.22 |
| animacy[Grammatical]:verbtype[ASV]:sag.[post] | 0.00049 | 0.013 | 0.039 |
| animacy[Grammatical]:task[C]:sag.[ant] | −0.00024 | 0.013 | −0.019 |
| animacy[Grammatical]:task[C]:sag.[post] | 0.004 | 0.013 | 0.31 |
| verbtype[ASV]:task[C]:sag.[ant] | −0.026 | 0.013 | −2.1 |
| verbtype[ASV]:task[C]:sag.[post] | 0.039 | 0.013 | 3.1 |
| animacy[Grammatical]:lat.[L]:sag.[ant] | 0.017 | 0.018 | 0.94 |
| animacy[Grammatical]:lat.[R]:sag.[ant] | −0.021 | 0.018 | −1.2 |
| animacy[Grammatical]:lat.[L]:sag.[post] | −0.0053 | 0.018 | −0.3 |
| animacy[Grammatical]:lat.[R]:sag.[post] | 0.02 | 0.018 | 1.1 |
| verbtype[ASV]:lat.[L]:sag.[ant] | 0.027 | 0.018 | 1.5 |
| verbtype[ASV]:lat.[R]:sag.[ant] | −0.021 | 0.018 | −1.2 |
| verbtype[ASV]:lat.[L]:sag.[post] | −0.031 | 0.018 | −1.7 |
| verbtype[ASV]:lat.[R]:sag.[post] | 0.013 | 0.018 | 0.71 |
| task[C]:lat.[L]:sag.[ant] | −0.032 | 0.018 | −1.8 |
| task[C]:lat.[R]:sag.[ant] | 0.015 | 0.018 | 0.84 |
| task[C]:lat.[L]:sag.[post] | 0.042 | 0.018 | 2.4 |
| task[C]:lat.[R]:sag.[post] | −0.017 | 0.018 | −0.93 |
| animacy[Grammatical]:verbtype[ASV]:task[C]:lat.[L] | 0.015 | 0.013 | 1.2 |
| animacy[Grammatical]:verbtype[ASV]:task[C]:lat.[R] | 0.017 | 0.013 | 1.3 |
| animacy[Grammatical]:verbtype[ASV]:task[C]:sag.[ant] | −0.0033 | 0.013 | −0.26 |
| animacy[Grammatical]:verbtype[ASV]:task[C]:sag.[post] | 0.0066 | 0.013 | 0.52 |
| animacy[Grammatical]:verbtype[ASV]:lat.[L]:sag.[ant] | −0.037 | 0.018 | −2.1 |
| animacy[Grammatical]:verbtype[ASV]:lat.[R]:sag.[ant] | 0.015 | 0.018 | 0.85 |
| animacy[Grammatical]:verbtype[ASV]:lat.[L]:sag.[post] | 0.034 | 0.018 | 1.9 |
| animacy[Grammatical]:verbtype[ASV]:lat.[R]:sag.[post] | −0.015 | 0.018 | −0.86 |
| animacy[Grammatical]:task[C]:lat.[L]:sag.[ant] | −0.029 | 0.018 | −1.6 |
| animacy[Grammatical]:task[C]:lat.[R]:sag.[ant] | 0.0059 | 0.018 | 0.33 |
| animacy[Grammatical]:task[C]:lat.[L]:sag.[post] | 0.026 | 0.018 | 1.5 |
| animacy[Grammatical]:task[C]:lat.[R]:sag.[post] | −0.0065 | 0.018 | −0.36 |
| verbtype[ASV]:task[C]:lat.[L]:sag.[ant] | 0.00095 | 0.018 | 0.053 |
| verbtype[ASV]:task[C]:lat.[R]:sag.[ant] | 0.0086 | 0.018 | 0.48 |
| verbtype[ASV]:task[C]:lat.[L]:sag.[post] | 0.014 | 0.018 | 0.8 |
| verbtype[ASV]:task[C]:lat.[R]:sag.[post] | −0.026 | 0.018 | −1.5 |
| animacy[Grammatical]:verbtype[ASV]:task[C]:lat.[L]:sag.[ant] | −0.012 | 0.018 | −0.64 |
| animacy[Grammatical]:verbtype[ASV]:task[C]:lat.[R]:sag.[ant] | 0.035 | 0.018 | 1.9 |
| animacy[Grammatical]:verbtype[ASV]:task[C]:lat.[L]:sag.[post] | 0.0033 | 0.018 | 0.18 |
| animacy[Grammatical]:verbtype[ASV]:task[C]:lat.[R]:sag.[post] | −0.023 | 0.018 | −1.3 |

Appendix C: ERP plots including confidence intervals by subject and by item

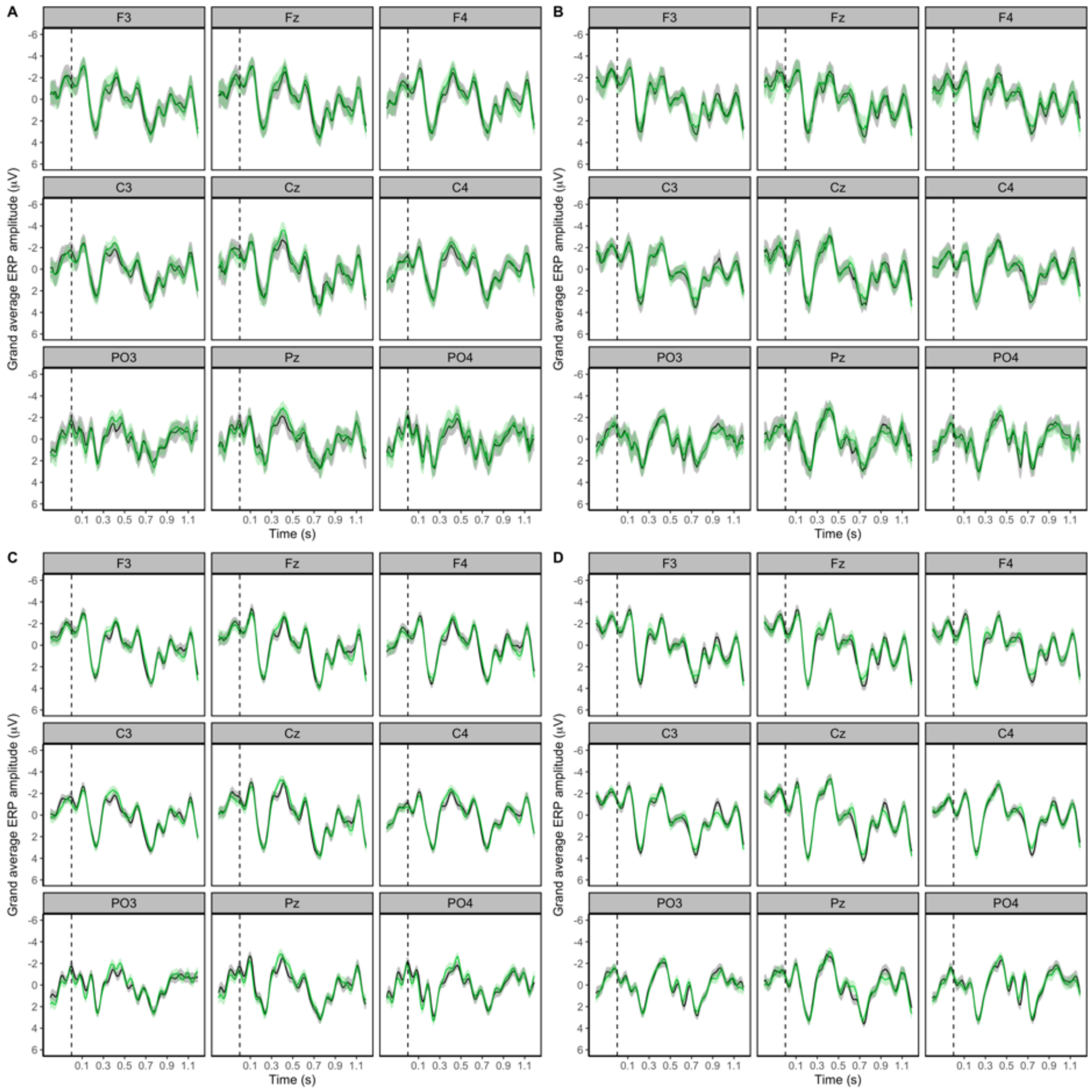


Figure 1. Grand average waveform of the visual ERPs elicited at the Subject Noun of Agent- and Experiencer- Subject Verb sentences (conditions collapsed) for the Judgement and Comprehension task conditions. Black lines represent animate Subject Nouns while green lines represent inanimate Subject Nouns. Negativity is plotted upwards and waveforms are time-locked to the onset of the noun with a -200ms to 0ms prestimulus interval. Shaded regions represent 95% confidence intervals (CIs) by subject or item. **A.** Judgement task group, by-subject CIs, **B.** Comprehension task group, by-subject CIs, **C.** Judgement task group, by-item CIs, **D.** Comprehension task group, by-item CIs. This plot corresponds to Figure 2 of the main text.

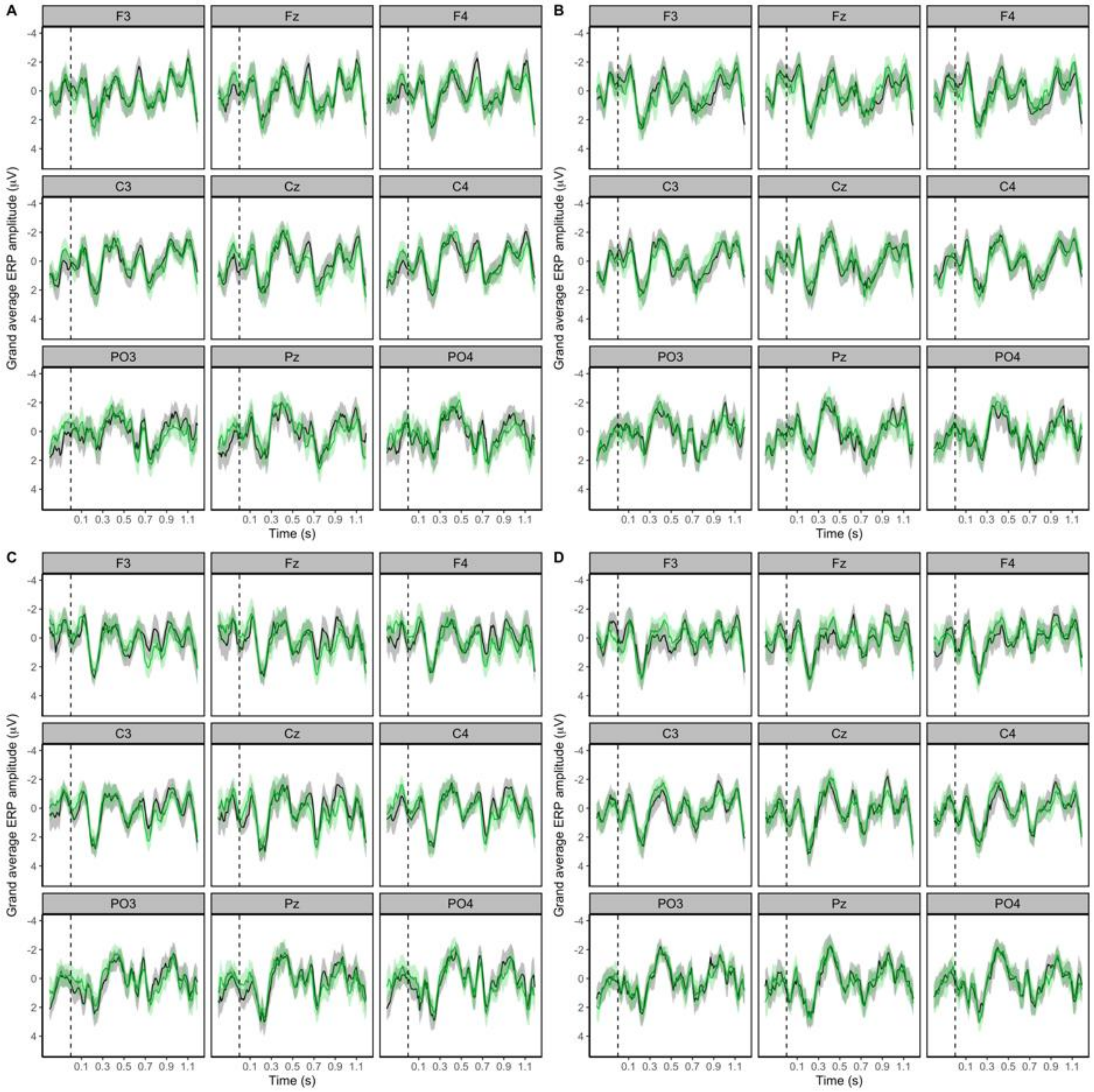


Figure 2. Grand average waveform of the visual ERPs elicited at the verb of Agent Subject Verb (ASV) and Experiencer Subject Verb (ESV) sentences for the Judgement and Comprehension task conditions. Black lines represent correct/grammatical sentences while green lines represent incorrect/ungrammatical sentences. Negativity is plotted upwards and waveforms are time-locked to the onset of the verb with a -200ms to 0ms prestimulus interval. Shaded regions represent 95% confidence intervals (CIs) by subject. **A.** Judgement task group, ASV, **B.** Judgement task group, ESV, **C.** Comprehension task group, ASV, **D.** Comprehension task group, ESV. This plot corresponds to Figure 4 of the main text.

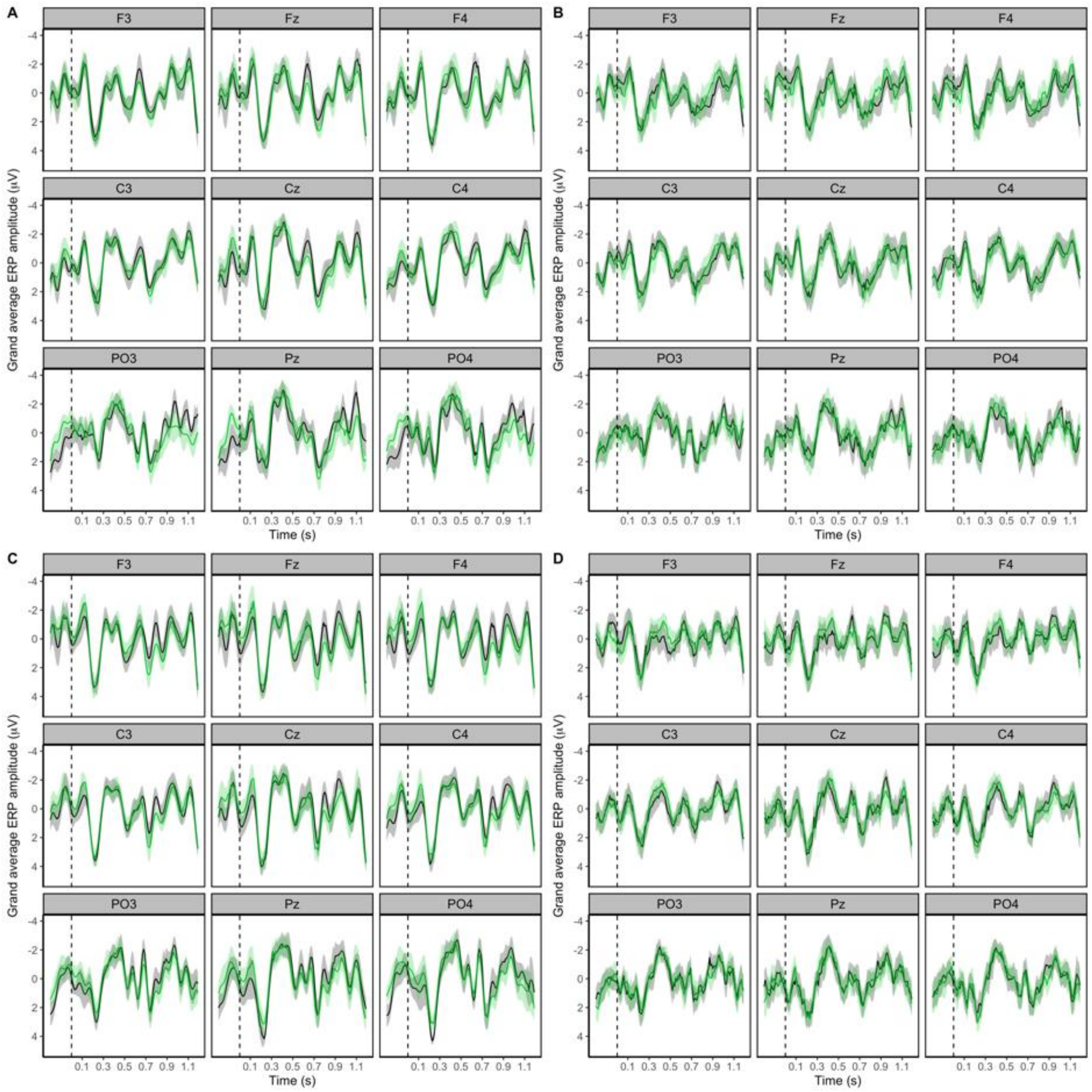


Figure 3. Grand average waveform of the visual ERPs elicited at the verb of Agent Subject Verb (ASV) and Experiencer Subject Verb (ESV) sentences for the Judgement and Comprehension task conditions. Black lines represent correct/grammatical sentences while green lines represent incorrect/ungrammatical sentences. Negativity is plotted upwards and waveforms are time-locked to the onset of the verb with a -200ms to 0ms prestimulus interval. Shaded regions represent 95% confidence intervals (CIs) by item. **A.** Judgement task group, ASV, **B.** Judgement task group, ESV, **C.** Comprehension task group, ASV, **D.** Comprehension task group, ESV. This plot corresponds to Figure 4 of the main text.

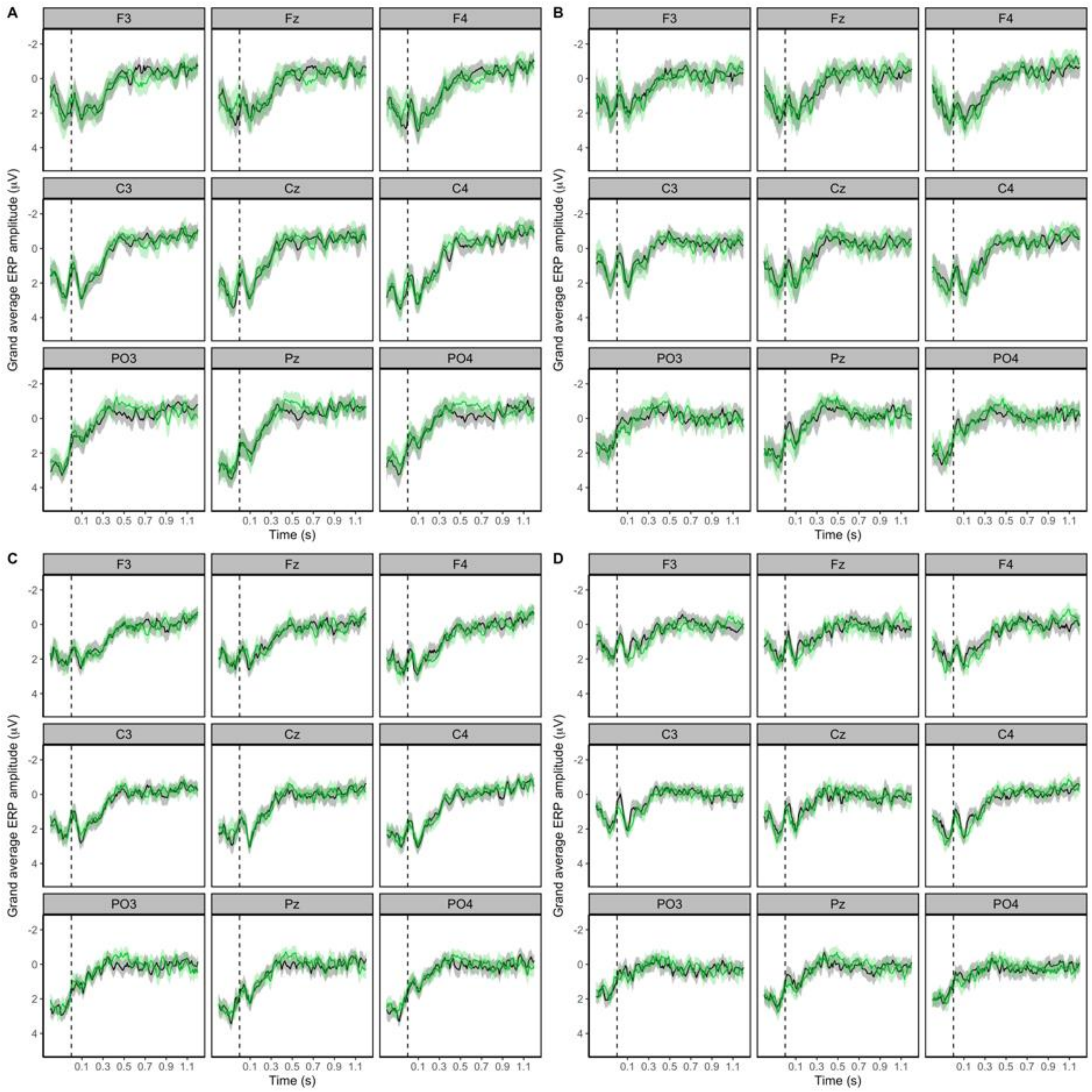


Figure 4. Grand average waveform of the auditory ERPs elicited at the Subject Noun of Agent- and Experiencer- Subject Verb sentences (conditions collapsed) for the Judgement and Comprehension task conditions. Black lines represent animate Subject Nouns while green lines represent inanimate Subject Nouns. Negativity is plotted upwards and waveforms are time-locked to the onset of the noun with a -200ms to 0ms prestimulus interval. Shaded regions represent 95% confidence intervals (CIs) by subject or item. **A.** Judgement task group, by-subject CIs, **B.** Comprehension task group, by-subject CIs, **C.** Judgement task group, by-item CIs, **D.** Comprehension task group, by-item CIs. This plot corresponds to Figure 6 of the main text.

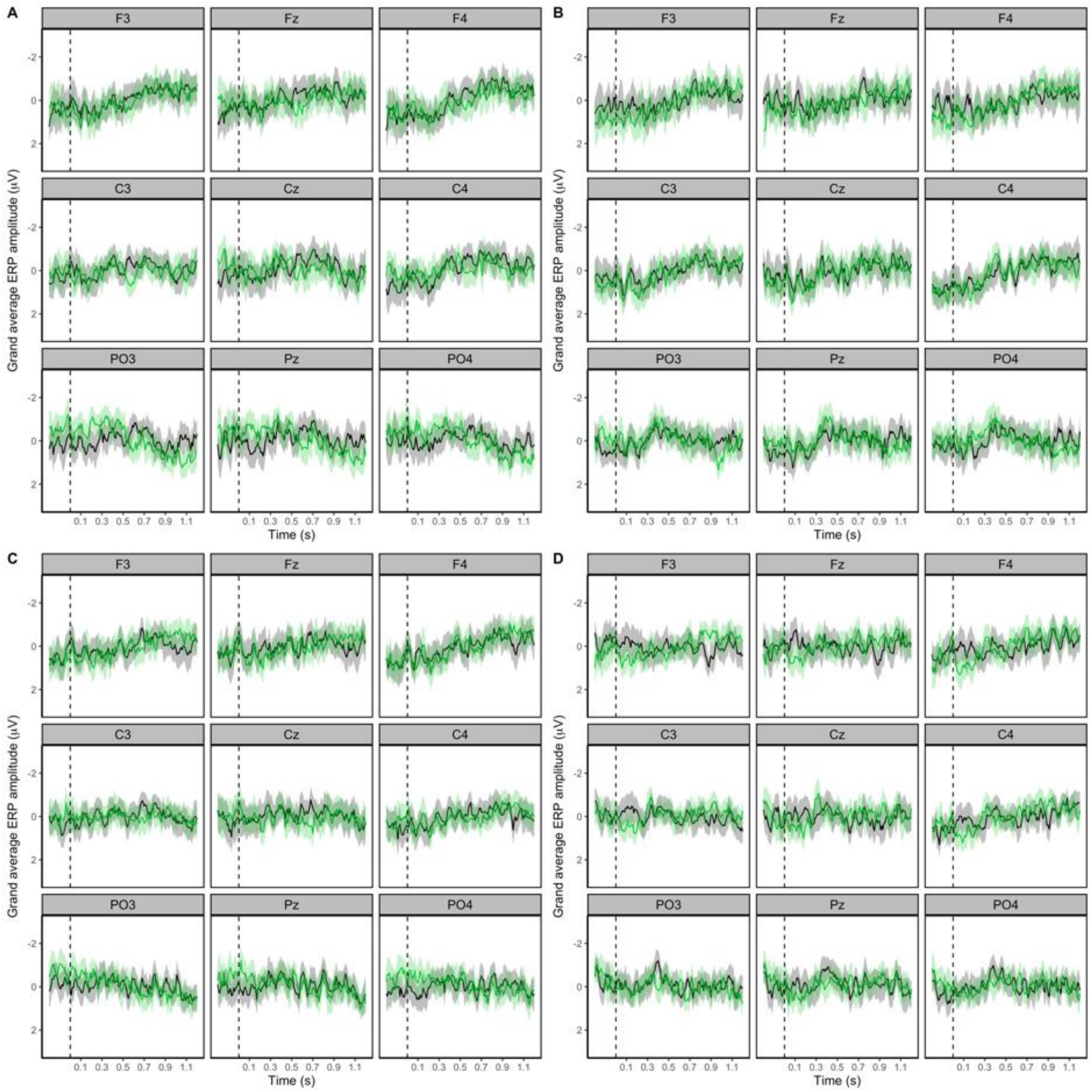


Figure 5. Grand average waveform of the auditory ERPs elicited at the verb of Agent Subject Verb (ASV) and Experiencer Subject Verb (ESV) sentences for the Judgement and Comprehension task conditions. Black lines represent correct/grammatical sentences while green lines represent incorrect/ungrammatical sentences. Negativity is plotted upwards and waveforms are time-locked to the onset of the verb with a -200ms to 0ms prestimulus interval. Shaded regions represent 95% confidence intervals (CIs) by subject. **A.** Judgement task group, ASV, **B.** Judgement task group, ESV, **C.** Comprehension task group, ASV, **D.** Comprehension task group, ESV. This plot corresponds to Figure 8 of the main text.

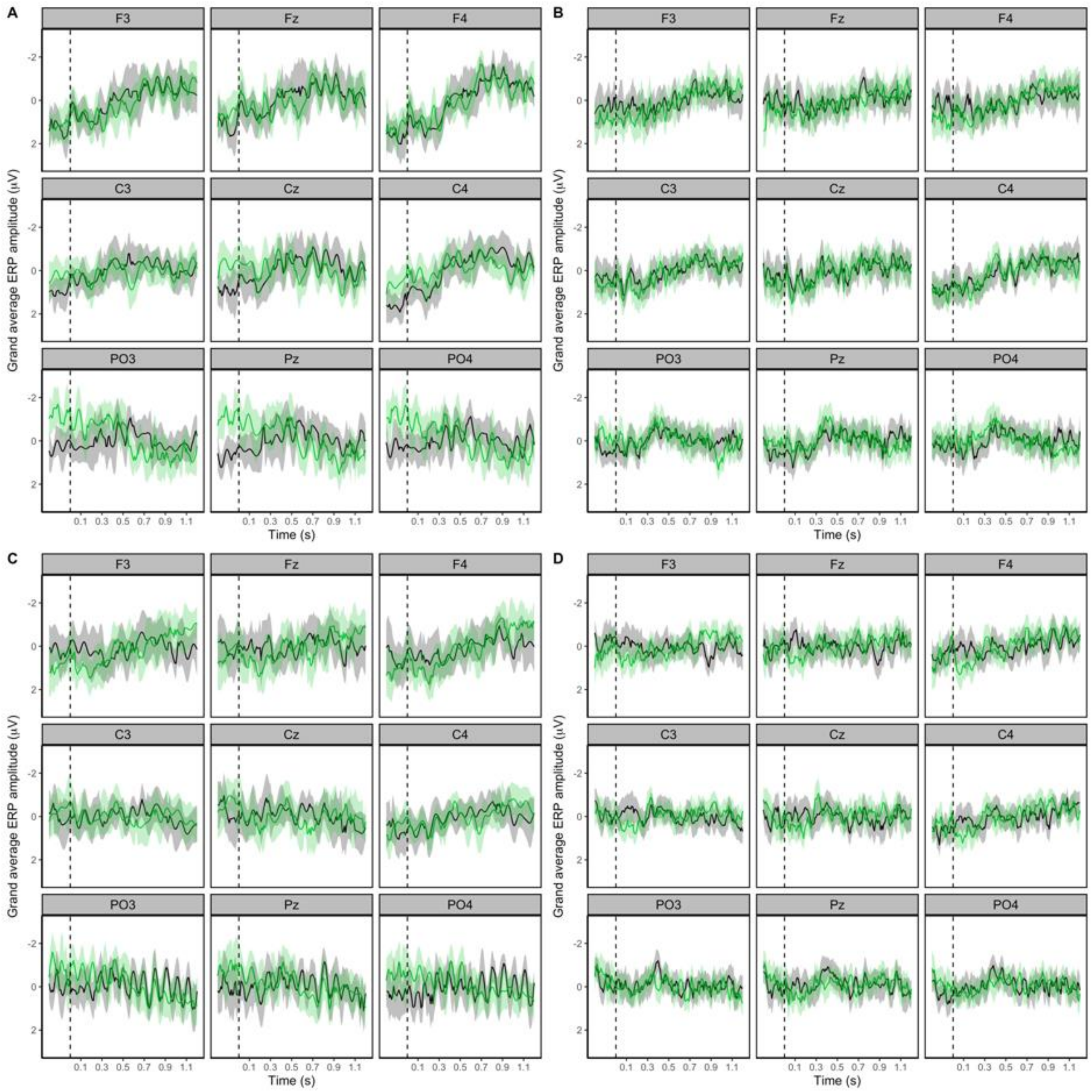


Figure 6. Grand average waveform of the auditory ERPs elicited at the verb of Agent Subject Verb (ASV) and Experiencer Subject Verb (ESV) sentences for the Judgement and Comprehension task conditions. Black lines represent correct/grammatical sentences while green lines represent incorrect/ungrammatical sentences. Negativity is plotted upwards and waveforms are time-locked to the onset of the verb with a -200ms to 0ms prestimulus interval. Shaded regions represent 95% confidence intervals (CIs) by item. **A.** Judgement task group, ASV, **B.** Judgement task group, ESV, **C.** Comprehension task group, ASV, **D.** Comprehension task group, ESV. This plot corresponds to Figure 8 of the main text.

#### Appendix D: Comparing behavioural accuracy variance by subject and item across modalities

To further investigate the variance in behavioural accuracy across the visual and auditory experiments, raincloud plots were developed (Allen et al., 2019). Figure 1 shows the by-participant and by-item variance for Experiment 1 (visual) while Figure 2 shows this variance for Experiment 2 (auditory). Response accuracy variance is relatively similar across modalities when aggregated by-participant. When aggregated by-item, response accuracy variance for the judgement condition is also similar across modalities. Most prominent is the by-item differences by modality for the comprehension task. Here, the by-item variance is larger for the visual modality, and smaller for the auditory modality with more of the data clustered around the mean and a smaller dispersion. These plots suggest that for some items, comprehension was consistently low in Experiment 1 (visual) as indexed by low response accuracy. In the auditory modality, these items did not have a similarly low accuracy score, indicating that some aspect of the auditory stimuli may have assisted in the comprehension of these "difficult-to-comprehend" items.

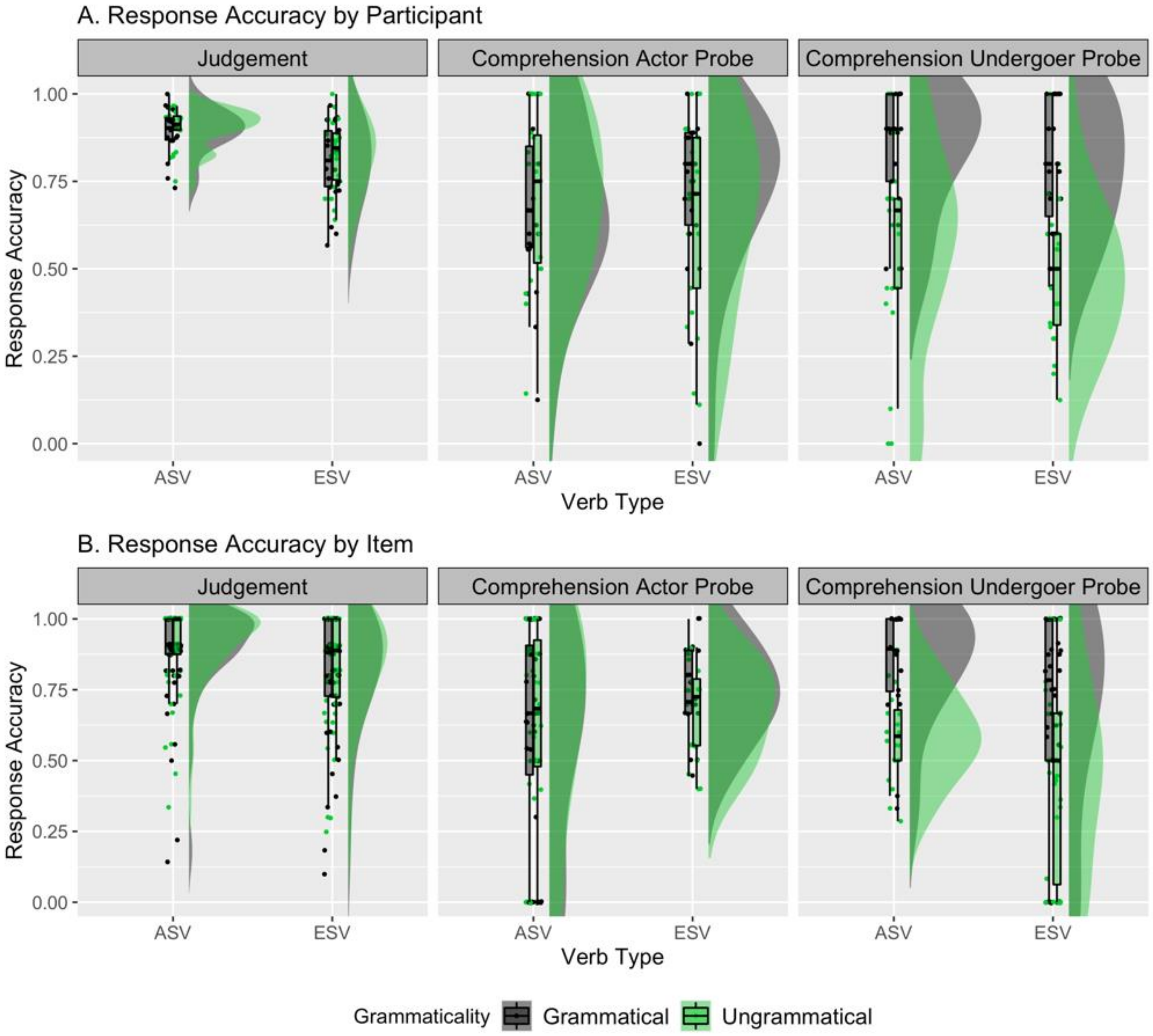


Figure 1. Behavioural response accuracy in Experiment 1 (visual) for Agent Subject Verbs (ASV) and Experiencer Subject Verbs (ESV) in the Judgement task condition and the Comprehension task condition (split into Actor probe questions and Undergoer Probe questions). Black dots represent grammatical sentences, while green dots represent ungrammatical sentences. Row A shows variability by participant, where individual data points represent the mean by-participant accuracy of ASV and ESV. Row B shows variability by item, where individual data points represent the mean by-item accuracy of ASV and ESV. Shaded areas represent the distribution of the data, where the width of the shaded area represents frequency.

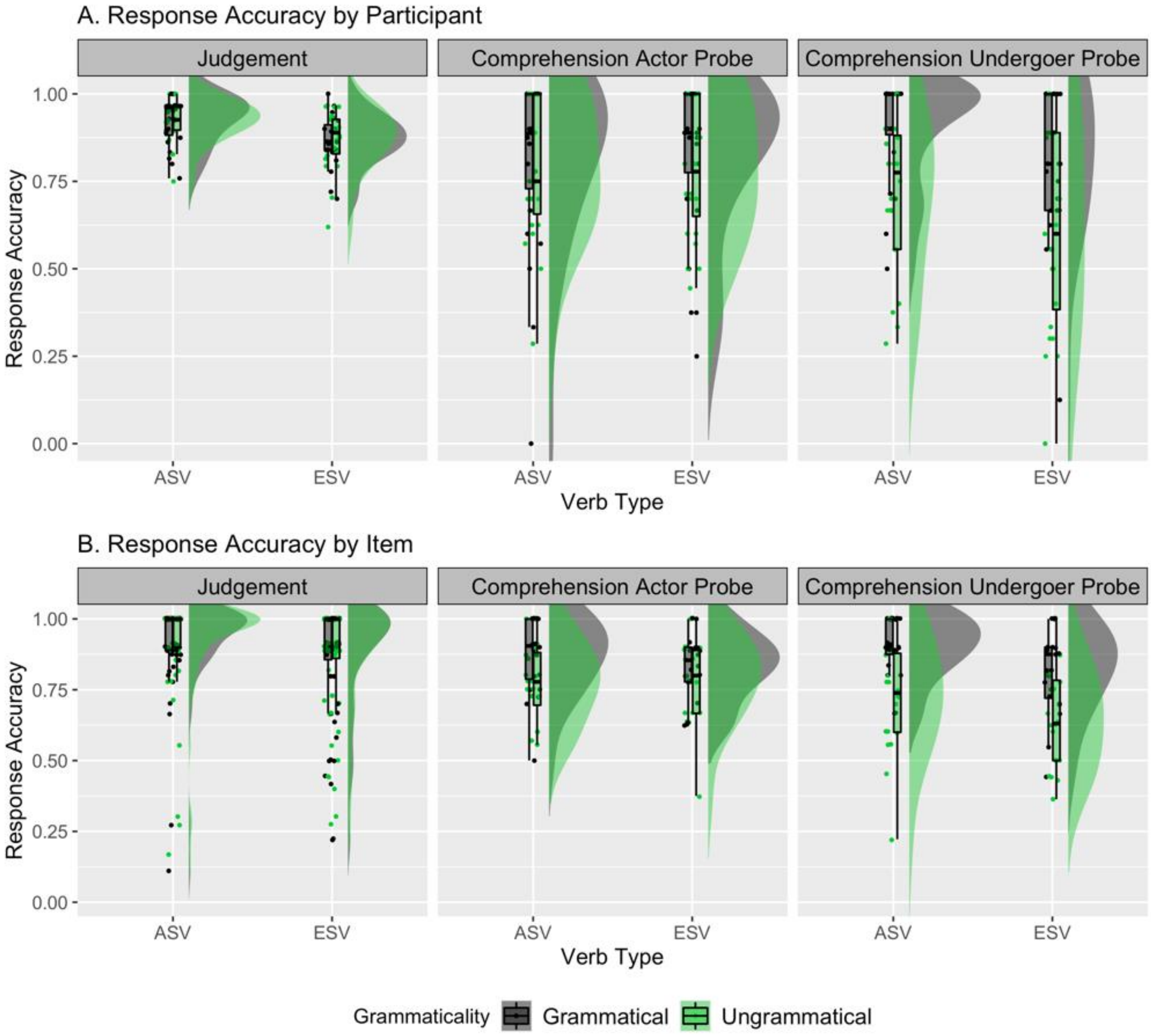


Figure 2. Behavioural response accuracy in Experiment 2 (auditory) for Agent Subject Verbs (ASV) and Experiencer Subject Verbs (ESV) in the Judgement task condition and the Comprehension task condition (split into Actor probe questions and Undergoer Probe questions). Black dots represent grammatical sentences, while green dots represent ungrammatical sentences. Row A shows variability by participant, where individual data points represent the mean by-participant accuracy of ASV and ESV. Row B shows variability by item, where individual data points represent the mean by-item accuracy of ASV and ESV. Shaded areas represent the distribution of the data, where the width of the shaded area represents frequency.

#### Appendix E: Phrase-length ERP plots for Experiment 2 (Auditory)

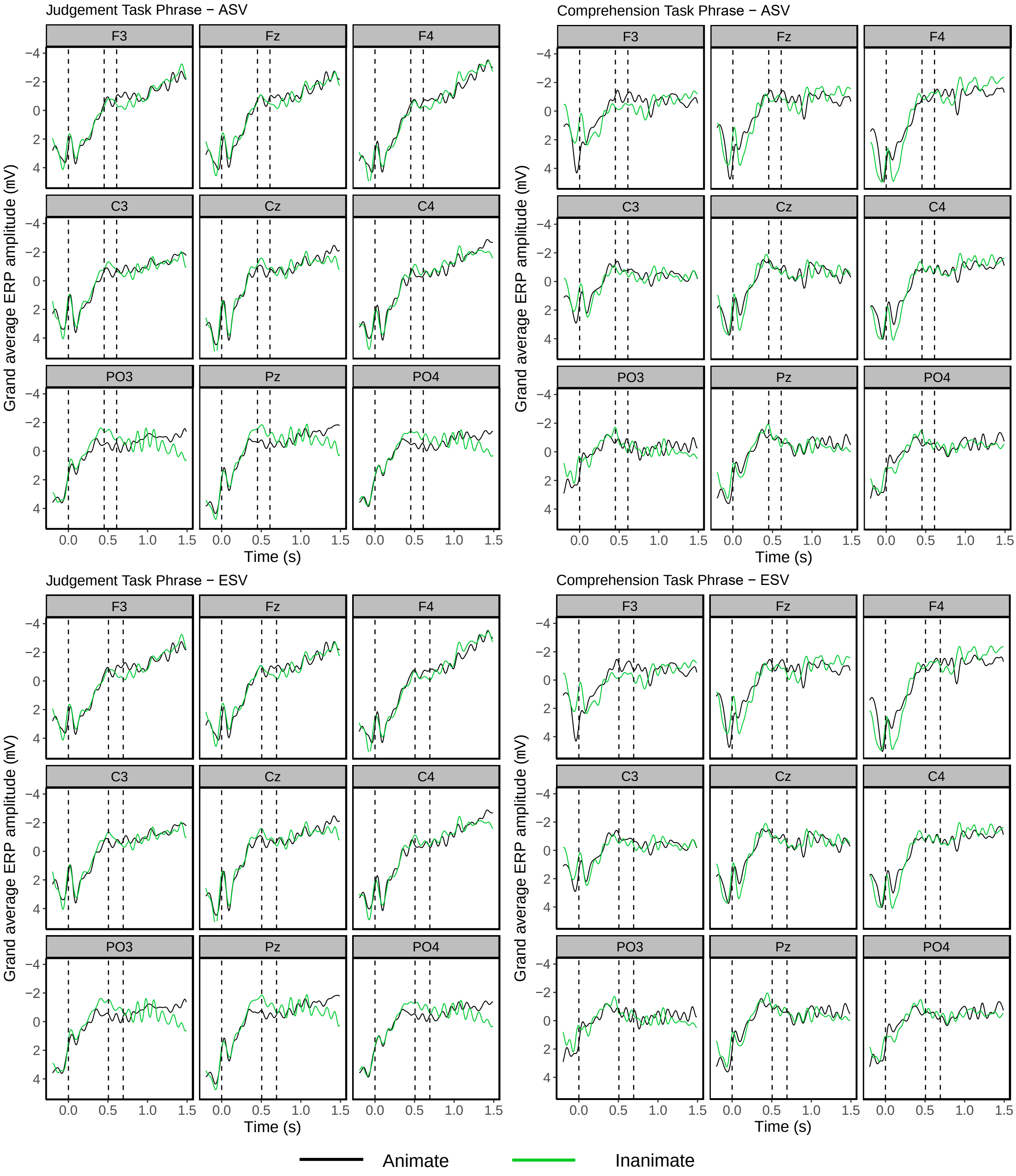


Figure 1. Grand average waveform of the auditory ERPs elicited at the Subject NP of Agent Subject Verb and Experiencer Subject Verb sentences (conditions collapsed) for both Judgement and Comprehension task groups. Black lines represent animate subject nouns while green lines represent inanimate subject nouns. Negativity is plotted upwards and waveforms are time-locked to the onset of the noun with a -200ms to 0ms prestimulus interval. The first dashed line indicates the onset of the subject noun, while the second dashed line indicates the approximate (average) onset of the auxiliary ("will"), and the third dashed line indicates the approximate (average) onset of the verb. These phrase-length ERP plots show that the prestimulus negativity seen for ungrammatical sentences at the verb (Figure 7 of main text) reflects the continuation of a negativity elicited by inanimate subjects.

#### Appendix F: Comparing random effects variance by subject and item

To investigate the relative variability of the random effects included in our Mixed Effects Models, we examined the random effects of Participant ID (Figure 1) and Item (Figure 2) using *sjPlot* (Lüdecke, 2019) in *R* (R Core Team, 2018) using the data from Experiment 2 (auditory). We plotted the random effects structures for the N400 time-window at the verb, using a subset of electrodes (middle laterality, central sagitality). Model structures included fixed effects of Animacy, Task, Verb Type with Scaled Prestimulus Amplitude included as a main effect. This matches the main analyses included in this paper. Here, random slopes of Animacy and Verb Type were included on the random effects of Subject and Item.

When visually comparing item-based variance (Figure 1) to subject-based variance (Figure 2) a wider range of variance from the intercept is evident for the fixed effects of Animacy, Verb Type, and the interaction of Animacy x Verb Type when plotted against the item intercept compared to the subject intercept. This represents a larger variability in the random effect of item compared to the variability in the random effect of subject. It is therefore possible that the larger amount of item-based variance may be driving the absence of a significant N400 to Experiencer Subject Verbs and absence of a P600 to Agent Subject Verbs, as further discussed in the main text.

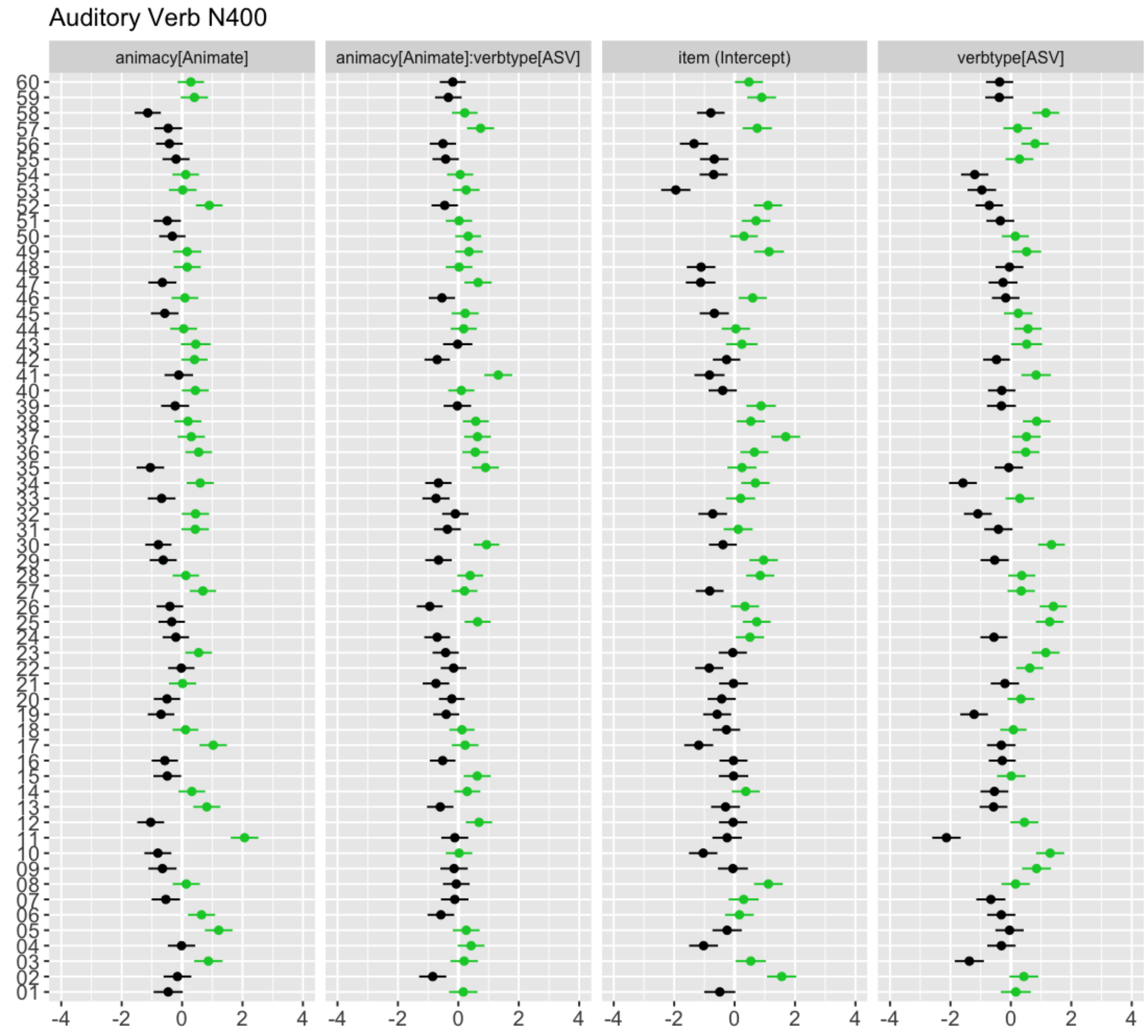


Figure 1. Mixed Effects Model fixed effects plotted against the intercept, grouped by item. These plots represent the intercept of the model examining the N400 ERP effect at the verb in Experiment 2 (auditory). Input data was from a subset of the electrophysiological data (middle laterality, central sagitality). Bars indicate 95% confidence intervals, black dots indicate a value below the intercept of the model, and green dots indicate a value above the intercept.

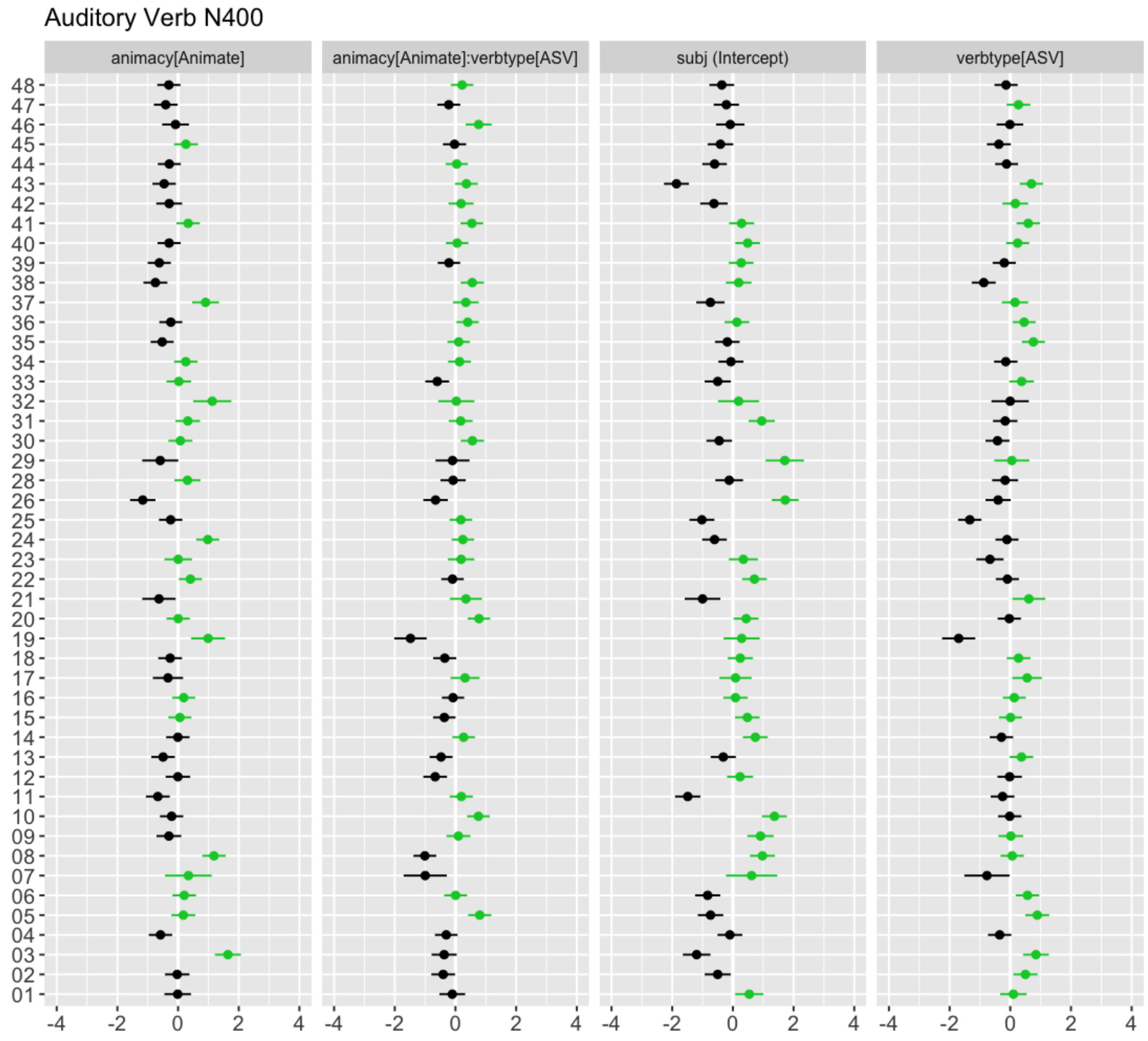


Figure 2. Mixed Effects Model fixed effects plotted against the intercept, grouped by subject. These plots represent the intercept of the model examining the N400 ERP effect at the verb in Experiment 2 (auditory). Input data was from a subset of the electrophysiological data (middle laterality, central sagitality). Bars indicate 95% confidence intervals, black dots indicate a value below the intercept of the model, and green dots indicate a value above the intercept.
